# Generative and interpretable machine learning for aptamer design and analysis of in vitro sequence selection

**DOI:** 10.1101/2022.03.12.484094

**Authors:** Andrea Di Gioacchino, Jonah Procyk, Marco Molari, John S. Schreck, Yu Zhou, Yan Liu, Rémi Monasson, Simona Cocco, Petr Šulc

**Author notes:** these authors contributed equally to this work and are ordered alphabetically (Andrea Di Gioacchino, Jonah Procyk). joint last authors (Rémi Monasson, Simona Cocco).

## Abstract

Selection protocols such as SELEX, where molecules are selected over multiple rounds for their ability to bind to a target molecule of interest, are popular methods for obtaining binders for diagnostic and therapeutic purposes. With the increasing amount of such high-throughput experimental data available, machine learning techniques have become increasingly popular for molecular datasets analysis. Here, we show that Restricted Boltzmann Machines (RBMs), a two-layer neural network architecture, can successfully be trained on sequence ensembles from SELEX experiments for thrombin aptamers, and used to estimate the fitness of the sequences obtained through the experimental protocol. As a direct consequence, we show that trained RBMs can be exploited to classify as well as generate novel molecules. To confirm our findings, we experimentally verify the generated sequences from RBM.

## 1 Introduction

Discovery and design of molecules that can specifically bind a given target molecule is a key problem in diagnostics, therapeutics and molecular biology in general. Multiple different experimental approaches exist to select specific molecular target binder such as antibodies, short peptides, proteins or small molecules. Single stranded oligonucleotides (DNA or RNA) have also been shown to be able to specifically bind with high affinity to a plethora of various targets, including small metabolites, proteins, nucleic acids, viruses, exosomes, and cells of specific tissue [27, 25, 41, 40, 37, 30, 36, 10, 7, 18], showing promise for applications that range from diagnostics to targeted disease therapy [51]. These short oligonucleotides, called aptamers, are selected from an initial pool of sequences by a procedure known as Systematic Evolution of Ligands by Exponential Enrichment (SELEX) [49, 12]. This method consists of multiple rounds of selection, where aptamers that bind strongly enough to the protein target are selected and amplified for the next round, until few strong binders are obtained. The advantages of using DNA or RNA include low cost of synthesising these molecules and relative ease of their manipulation in the laboratory setting as opposed to other selection methods such as peptide or antibody selection [43, 35]. Oligonucleotides can be denatured and refolded many times, allowing for multiple selection rounds. On the other hand, as they are composed of four possible types of bases (A, C, G and T/U), they do not offer such chemical diversity as antibodies. Thus, the range of targets that aptamers can be selected to bind strongly to is limited to some extent. However, chemical modifications of the nucleic bases can increase the chemical space of the aptamers and provide diverse sequence libraries from which strong binders can be selected against a variety of targets [13].

With the advance of next generation sequencing and high-throughput biological and molecular dataset production, various machine learning methods have been used to process biological sequences datasets, with applications including classifications, binding prediction, and molecular design [11]. While a significant improvement has recently been achieved in using deep learning for protein or RNA structure predictions [20, 46], predictions of binding interactions and *de novo* design of molecular binders remain outstanding significant challenges. So far, it is primarily the prediction of interaction between a small molecule ligand and a target protein that has received attention from the machine learning community, as such approaches are at the basis of the drug screening pipeline [3]. Motif-finding and clustering-based methods, combined with secondary structure prediction tools, have been previously developed for processing SELEX datasets [17, 44, 1, 1, 2, 19]. Currently, the SELEX dataset processing typically involves clustering and identifying a common motif in aligned sequences and then selecting representative aptamers from the last round of selection and verifying their binding affinity to the target.

A challenging task in the analysis of SELEX experiments is the quantification of the aptamer fitness, which determines the sequence landscape evolution at each selection round. Several approaches have been introduced in the past, based on *in silico* molecular dynamics simulations [54, 53], on clustering in sequence space together with enrichment measurements [33], and on additional, direct fitness estimation experiments [34]. These methods proved useful to estimate the fitness of a limited number of selected sequences or of large classes of similar sequences, but they seem unable to assign in a reliable way a fitness score to each molecule observed in final rounds of SELEX.

Over the last decade, deep neural networks (DNN) have become a popular machine learning tool in many areas, such as image recognition or natural language processing, and are now increasingly applied in chemical and biological data processing workflow [22, 56, 23, 6]. However, training DNNs typically requires large datasets, which can be challenging and expensive to obtain from biological experiments. DNNs have many free parameters, which makes it difficult to identify and interpret particular features of the molecule that are attributed to its ability to bind a given target. The presence of errors in the the sequence dataset, coming e.g. from experimental error in affinity measurements or sequencing errors, adds further difficulties to training as well as to interpretability. Machine-learning methods for sequence ensembles include inverse models from statistical physics, such as direct coupling analysis (DCA) methods [8], which have been previously successfully used to infer native contacts and guide folding of RNA and proteins based on homologous sequence alignment [9, 28], as well as to generate functional enzymes based on functional protein alignments [39] and protein recognizing RNA [52]. They infer parameters of maximum-entropy models, which are fixed by the requirement that the conservation of single residues and pairs of residues given by the model match the values observed in the sequence alignment. More recently, Restricted Boltzmann Machine (RBM) architectures, a neural network with a bipartite graph structure, have been successfully applied as a generative model for protein domain sequences [47], as well as a predictor of peptides that will be presented on MHC complexes [5]. They present an intermediate level of complexity between the direct coupling models and DNNs, as they can be trained to recognize multi-residue coupling as opposed to pairwise interactions, but due to limited number of weights between the two neuron layers, the parameters can still be interpreted and rationalized.

Here, we apply RBM models to a set of DNA sequences obtained from the prior experimental work of some of us that used SELEX method to obtain thrombin aptamers [55] (Figure 1). We show that the sequence likelihood assigned by the RBM can be directly related to the fitness of that sequence in the experimental selection. Moreover an RBM model that is trained on an earlier round of the selection is able to predict fitness of sequences in the next rounds not seen during the training, showing remarkable generalization capabilities. We further show that we can identify the sequence motifs conferring large likelihood to an aptamer sequence and that RBM’s hidden unit input can be used to cluster sequences. We show the capability of the RBM to predict binding affinity and generate new monovalent aptamers, which are good binders to one of the two thrombin binding sites, by gel shift assays. We investigate how taking into account the individual sequence counts from the experiment in the training data changes the properties of the inferred RBM model. Lastly, we also explore several supervised learning approaches that include random forest and various DNN architectures, but find them difficult to train and with poor generalization performance on our dataset.

**Figure 1:**
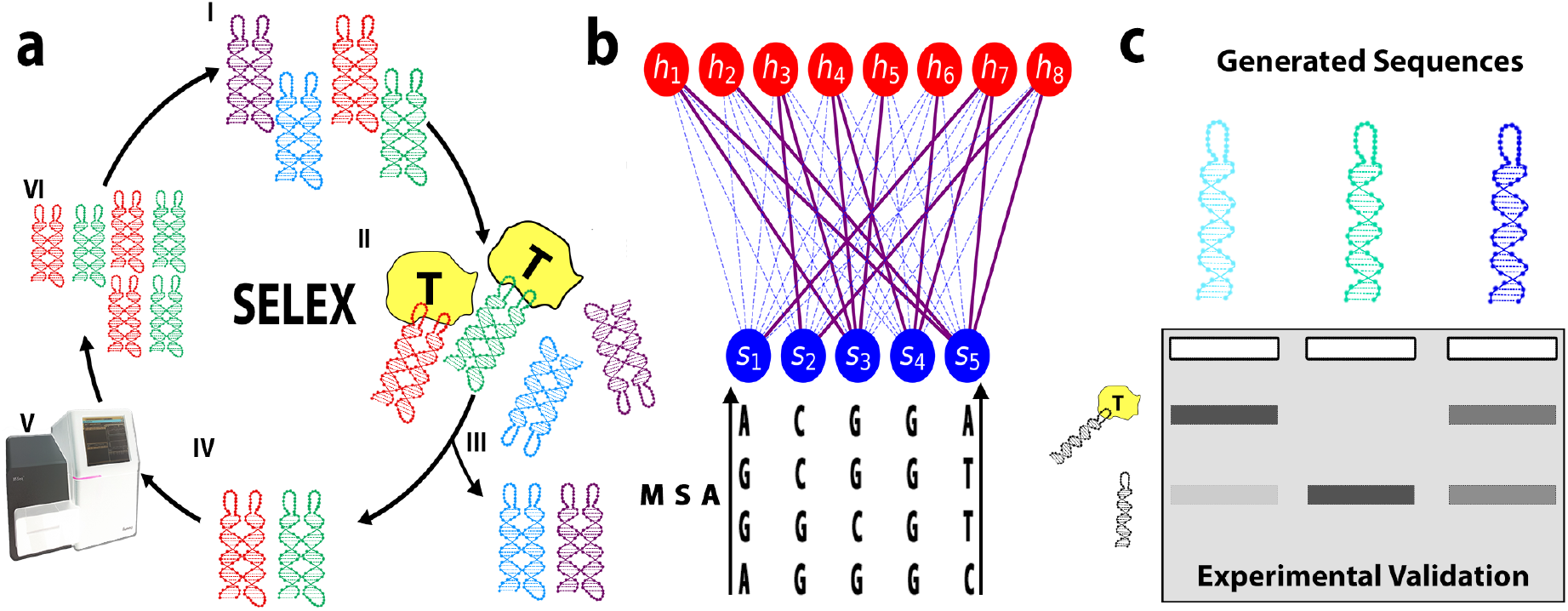
Schematic view of the SELEX experiment and the RBM-based analysis. **a:** The SELEX procedure used to obtain DNA aptamers that bind to thrombin consists of the following steps: I) We start with an initial library of DNA sequences. II) DNA aptamers compete with each other to bind to thrombin. III) Sequences that are unbound (or bound too weakly) are washed away. IV) Remaining bound sequences dissociate after the sample is heated up. V) Binding sequences are sequenced. VI) Using polymerase chain reaction (PCR), multiple copies are made of the remaining sequences, resulting into a new library of aptamers for the next round of selection. **b:** The sequenced aptamers from respective rounds of the SELEX protocol are used to train the parameters of the Restricted Boltzmann Machine model. In this unsupervised neural network architecture, a layer of visible units carry the aptamer sequence, while the layer of hidden units extract representations. The weighted connections between the two layers are learned through maximization of the log-likelihood of the sequences obtained through SELEX. **c:** Single loop sequences generated using the Restricted Boltzmann Machine model are experimentally validated using gel assays.

## 2 Results

### 2.1 Dataset obtained from SELEX Procedure

In a prior work [55], some of us used the SELEX method to obtain a bivalent DNA nanostructure that binds to a thrombin protein. In this DNA SELEX procedure, an initial library of about 10^15^ unique DNA sequences with all about the same length were exposed to the target tethered to a surface. The non-binding sequences were then washed away, while the binding sequences were collected (and optionally also sequenced). After amplification with PCR they served as the sequence library for the next cycle of SELEX. Cycles were repeated until binders of the desired binding affinity were found. The washing intensity was increased in later rounds to obtain stronger binders. In the particular experimental dataset used in Ref. [55], the SELEX procedure was performed on a DNA nanotile (Fig. 1), consisting of a joined-double helix region with two loops of 20 nucleotides each. While the double-helix nanotile structure was conserved across all DNA structures, the two respective loops were variable, starting from the initial random library. The SELEX procedure is schematically shown in Fig. 1 and consisted of eight selection rounds. The binding molecules were sequenced in rounds 5 (891959 sequences out of which 891914 unique), 6 (736436 sequences out of which 735974 unique), 7 (750926 sequences out of which 744597 unique) and 8 (725431 sequences out of which 719413 unique), and form the datasets we use here for training our models.

For each round, our dataset includes the sequence of the two (left and right) respective variable loop regions of the DNA nanotile, as well as the number of counts of the two-loop sequence, corresponding to the number of times it was sequenced in the experiment. In typical SELEX protocols, the sequences with the largest number of counts in the last rounds are considered the best binders.

### 2.2 Restricted Boltzmann Machine model

We use a Restricted Boltzmann Machine (RBM) to learn the probability distribution over the set of aptamers based on the sequences collected through the SELEX procedure. An RBM is a probabilistic model, represented by a bipartite graph consisting of *L* “visible” and *M* “hidden” units (shown schematically in Fig. 1b). It assigns a probability *p*(***s**, **h***) to a system state, given by two parts: the configuration of visible units, ***s*** = (*s*_1_,…, *s*_L_), where *s_i_* = A, C, G or T are the nucleotides on site i along the aptamer sequence, and the configuration of the hidden units, ***h*** = (*h*_1_,…, *h_M_*), meant to extract latent factors of variation in the visible configurations. The likelihood of a sequence **s** is formally obtained by marginalizing over all possible latent configurations (not observed in the data), *p*(***s***) = ∫ *dh p*(***s***, ***h***). The number *L* of visible units can be set to 40 to model full two-loop sequences or restricted to 20 to describe each loop independently. These two possibilities will be referred to as, respectively, D (Double loop) and S (Single loop) in the following.

Training a RBM consists in finding the parameters (in particular, the couplings between the layers) so that the log-likelihood of the observed data,

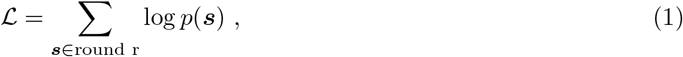

is maximized. Here the sum over ***s*** is over the sequences observed at a fixed selection round, say, *r*, of the SELEX experiment. Each sequence may therefore appear multiple times, depending on the number of its counts. We will denote this model with C (Count). An alternative is to include in the sum in Eq. (1) unique sequences only. The resulting model, labelled with U (Unique), has different properties, which we will discussed below.

The maximization of 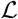 is a computationally difficult problem, but several effective techniques to obtain good parameter values have been developed, for instance contrastive divergence [16] and persistent contrastive divergence [45]. As described in Methods Sec. 4.2, we train, following Ref. [47], the RBM using persistent contrastive divergence and using double Rectified Linear hidden units, with a 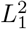 regularization scheme. This regularization favors sparse weights, and enhances interpretability of the trained model.

### 2.3 RBM’s log-likelihood is an accurate predictor of the aptamer’s fitness

Fig. 2a shows the distributions of log-likelihoods of sequences collected at SELEX rounds *r* = 5 to 8, estimated with an RBM trained on double-loop aptamer sequences with counts measured at round 6 (RBM-DC, see Suppl. Sec. S3). At round 5 three peaks are apparent. The logos of the sequences in each peak are shown in Fig. 2b. The peak at low log-likelihoods is characterized by highly variable sequences, weakly enriched in C, G nucleotides. The peak at intermediate values correspond to sequences with a structured loop (the left one, for most sequences), including a G-quadruplex motif. In the high loglikelihood peak a similar G-quadruplex motif appears on both left and right loops (for more details, see also Sec. 2.4). From round 6 to 8 the peaks at low and intermediate log-likelihood values are progressively depleted, and the peak at high log-likelihood gets more and more populated. This enrichment strongly suggests a positive correlation between the score assigned by the RBM and the fitness.

**Figure 2:**
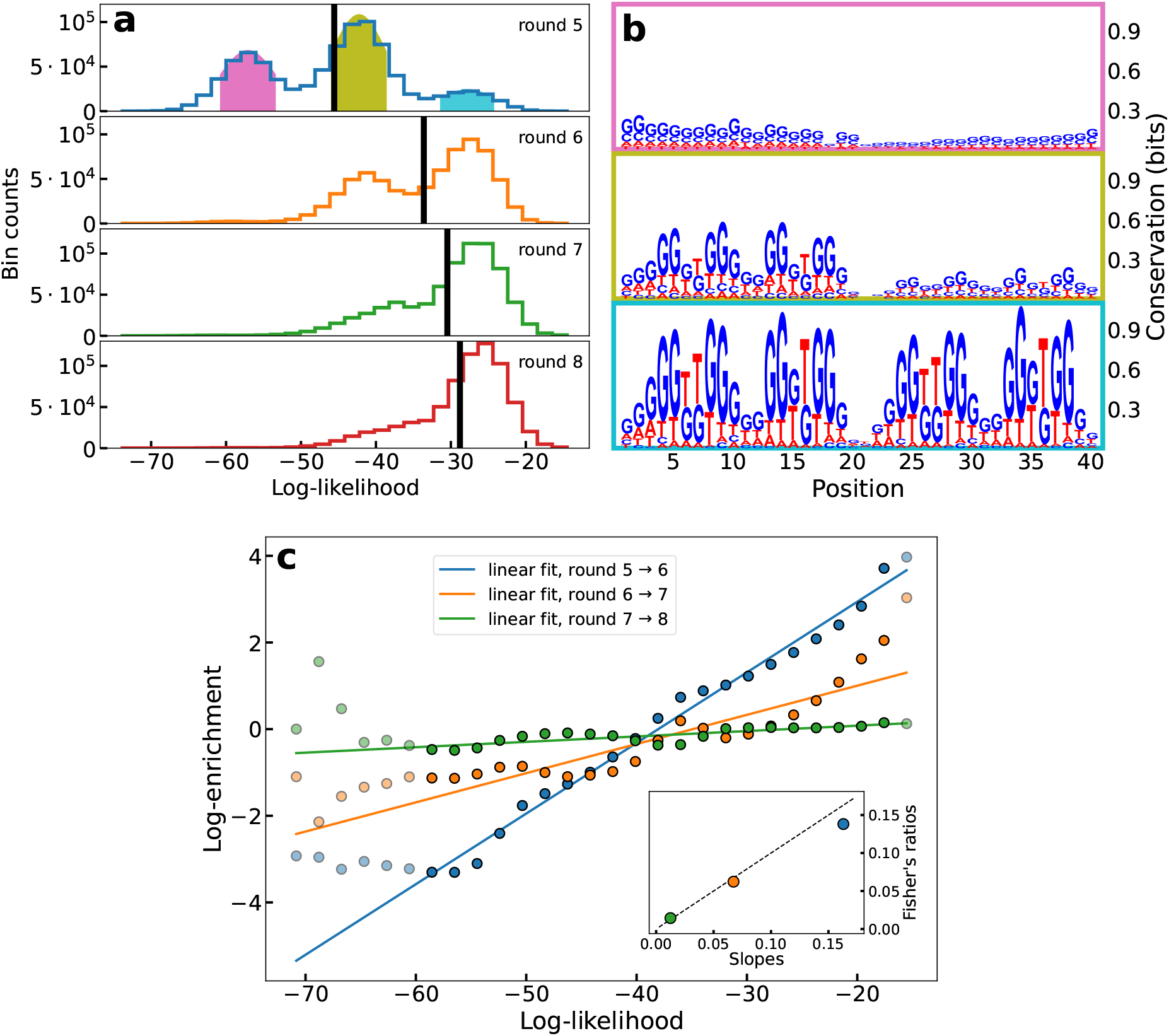
The RBM log-likelihood is strongly correlated to sequence fitness. **a**: Histograms of the log-likelihood of all sequences in the dataset at different rounds, obtained through RBM-DC trained on the sequences from round 6. The black line denotes the average log-likelihood. **b**: Logos of the sequences in each colored-shaded peak of the log-likelihood observed at round 5. **c**: Enrichment log-ratios 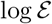 vs. log-likelihoods log*p* averages over the sequences in each bin of the histograms of panel a. The three sets of points corresponds to rounds *r* = 6, 7, 8. Linear fit are estimated from points with log-likelihood in the interval (−60, −17) (not shaded in the plot) only, to exclude undersampled bins cumulatively representing 0.5%, 0.3%, and 0.3% of the sequences, respectively at rounds 6, 7 and 8. Inset: scatter plot of the slopes of the linear fits (x-axis) and of the log-likelihood Fisher ratios (y-axis); linear dashed line: *y* = *x*.

In a population genetic framework, the fraction *q* of aptamers with sequence ***s*** changes from round *r* – 1 to round *r* according to

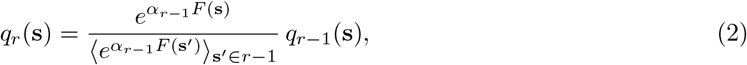

where 〈*O*(**s**′)〉_*s*′_∈_*r*-1_ = ∑_**s**′_ *q*_*r*–1_(**s′**)*O*(**s′**) denotes the average of the observable *O* over the distribution of sequences at round *r* – 1. The fitness *F*(**s**) encompasses the capability of an aptamer ***s*** of binding its target, as well as other chemical properties, such as its affinity to PCR amplification. Parameter *α*_*r*–1_ represents the selection strength from round *r* – 1 to *r*, which can be tuned in practice e.g. by varying with the intensity of washing in SELEX selection.

According to Eq. (2), formally valid for an infinite-size population only, the fitness *α*_*r*–1_ *F*(**s**) is, up to a sequence-independent additive constant, equal to the logarithm of the enrichment ratio 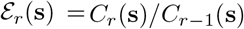, where *C*_r_(**s**) is the number of counts of sequence **s** at round *r*. However, the extreme subsampling of sequences at each round in our dataset prevents us from using empirical enrichment ratios 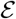 to estimate the fitnesses, and their correlation with log-likelihoods, see Suppl. Fig. S19. For instance, only *f*_shared_ = 0.5% of the sequences observed in round 7 or round 8 are present in both rounds, and among these sequences, about *f*_1_ = 70% have count *C* = 1 in both rounds. In earlier rounds, e.g. 5 and 6, the situations is even worse, with fractions *f*_shared_ = 0.01% and *f*_1_ = 93%.

To obtain more reliable enrichment ratios we gather all sequences **s** having similar log-likelihoods log*p*(***s***), and introduce their cumulative number of counts, *C*(ℓ, *r*). More precisely, *C*(ℓ, *r*) is defined as the number of counts in the *ℓ^th^* bin of the histogram of log-likelihoods in Fig. 2a. We then define the effective enrichment ratio of bin *ℓ* through 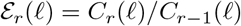. Fig. 2c shows the scatter plots of the enrichment log-ratios log 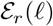 vs. the log-likelihoods *ℝ*, for rounds *r* = 6, 7, 8. Very strong correlations are observed, with coefficients of determination *R*^2^ = 0.99, 0.83 and 0.66 and slopes 0.16, 0.07, 0.01 for, respectively, the pairs of rounds 5 → 6, 6 → 7, and 7 → 8. The smaller values of the slopes of the linear regressions at later rounds suggests that the effective selection strength *α*_*r*–1_ appearing in Eq. (2) is weaker in the last SELEX rounds than in the previous ones. This interpretation is supported by the fact that the 10 different single-loop aptamers with largest count numbers at round 8 do not increase exponentially in the last rounds considered here, as shown in Suppl. Fig. S15.

The linear relationship between the RBM log-likelihood log*p*(***s***) and the sequence fitness *F*(***s***) suggests an alternative way to estimate the selection strengths *α_r_*. Fisher’s fundamental theorem (see for instance [29] for a review) postulates that the selection strength can be estimated through the ratio of the increase of the average fitness and of the its variance, *α*_*r*–1_ = (〈*F*〉_*r*_ – 〈*F*〉_*r*–1_var(*F*). We compute these Fisher’s ratios using log *p* as a proxy for *F* to estimate the selection strengths at the various rounds. Results are shown in the inset of Fig. 2c, and agree with those obtained directly from the slopes of the linear regressions. The precise relation between the fitness and the log-likelihood is further examined in Discussion section.

### 2.4 The log-likelihoods of the aptamers can be explained by the additive contributions of their left and right loops

To examine the cooperative binding of the left and right loops of the aptamer nanostructure at a given round of SELEX, we have trained RBM models on the 20 nucleotide-long single loops only. In practice, RBM-SC trained on all left (L) loop subsequences, on all right (R) loop subsequences, or on both of them show very similar properties (Suppl. Fig. S16), and we hereafter report results with the latter model. We show in Fig. 3a the log-likelihoods of the L and R loops for all aptamers at round 5. We observe the presence of four peaks in the joint distribution, corresponding to all possible combinations of the two peaks at, respectively, low 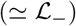 and high 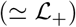 log-likelihoods present in the marginal distributions for L or R loops.

**Figure 3:**
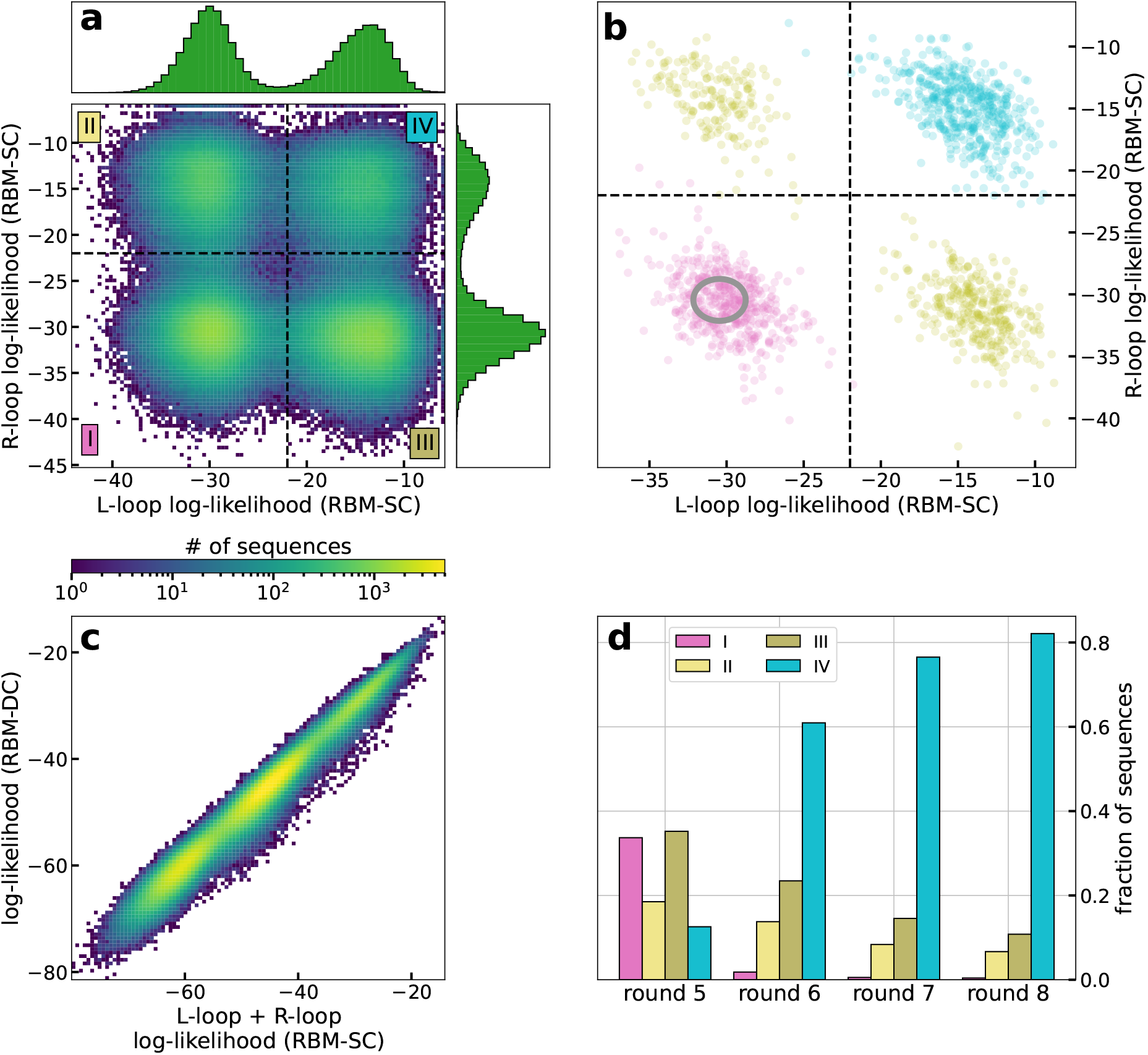
Contribution of left and right loops in the RBM log-likelihood. **a**: Joint distribution of the log-likelihoods of the L and R loop subsequences at round 5, estimated with RBM-SC trained on subsequences at round 8. The insets show the marginal distributions for both loops. **b**: Same as panel a for 2000 aptamer sequences attached to each of the three colored peaks in Fig. 2a (same color code). The gray ellipse shows the distribution of the log-likelihoods of uniform random sequences (center: mean values, ellipse: 2 standard deviations from the mean). **c**: Log-likelihoods of the aptamers in round 5 (estimated with RBM-DC see Suppl. Sec. S3) vs. sums of the log-likelihoods of their L and R loops (estimated with RBM-SC). **d**: Fractions of sequences in regions I to IV of panel a at rounds 5, 6, 7 and 8, estimated with the same RBM-SC model as in panel a.

As shown in Fig. 3b, aptamer sequences previously characterized as having low (in pink), intermediate (in olive) and high (in turquoise) log-likelihoods, see Fig. 2a, occupy the four corners of the joint-distribution plot. Therefore, high-log-likelihood aptamers have both L and R loops with high loglikelihoods 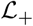, while the L and R loops of the low-log-likelihood aptamers have both low log-likelihoods 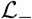. Intermediate aptamers have one loop, either L or R, with high loglikehood value 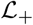 and the other with low log-likelihood 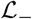.

Fig. 3c shows the scatter plot of the log-likelihoods of the full aptamers (estimated with RBM-DC) vs. the sums of the log-likelihoods of their L and R loops (estimated with RBM-SC). We observe an excellent linear correlation (*R*^2^ = 0.99), indicating that both loops contribute additively to the score of the full aptamer. This linearity also explains the three peak structure of the aptamer log-likelihoods in Fig. 2a, approximately located at 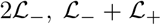, and 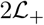. Moreover, thanks to this linearity, the selection of the aptamer population from one SELEX round to the next one (Fig. 2) can be predicted also at the level of single-loop aptamers (see Suppl. Fig. S27).

Fig. 3d shows the fractions of sequences in the four regions labelled I to IV of the L and R loglikelihoods at successive rounds of selection, see Fig. 3a. As observed in Fig. 2a for the full aptamer sequences we see a progressive enrichment in sequences for which both L and R loops have high loglikelihoods. However, we also observe a substantial fraction of sequences (> 15%) at round 8, in which one loop only has high log-likelihood. The cognate 20-nucleotide sequences, with low log-likelihood on the other loop, will be called parasite in the following, as they are likely to be selected only due to the ability of the other loop to bind thrombin. To check this hypothesis we generate random aptamer sequences, in which the 40 nucleotides are drawn uniformly at random. As shown in Fig. 3b these random aptamer sequences are located in the 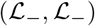 corner, and do not differ much from the pink sequences in terms of log-likelihood, see gray ellipse in Fig. 3b. Notice that removing the parasite sequences from the training set of RBM-SC does not significantly modify the estimation of log-likelihoods, see Supp. Fig. S17, which shows the robustness of the RBM model against the presence of random sequences in the data. The identification of parasite sequences has important consequences for the design of new aptamers based on the RBM model, as discussed in the next section.

### 2.5 RBM parameters reveal functional features of the aptamer sequences

We next extract the features that contribute the most to the likelihood of the sequences by studying weights between hidden units and visible layer (Fig. 1b). To enhance the interpretability of the RBM weights connecting input and and hidden layers we enforce their sparsity through appropriate regularisation (see Methods Sec. 4.2 and Ref. [47]). Figs. 4a-c (left) show the sequence logo of the three weights of RBM-DC with largest Frobenius norms (Suppl. Fig. S20); the height of nucleotide symbol *s* in position *i* for hidden unit *μ* represents the value of the weight *w_μi_*(*s*).

**Figure 4:**
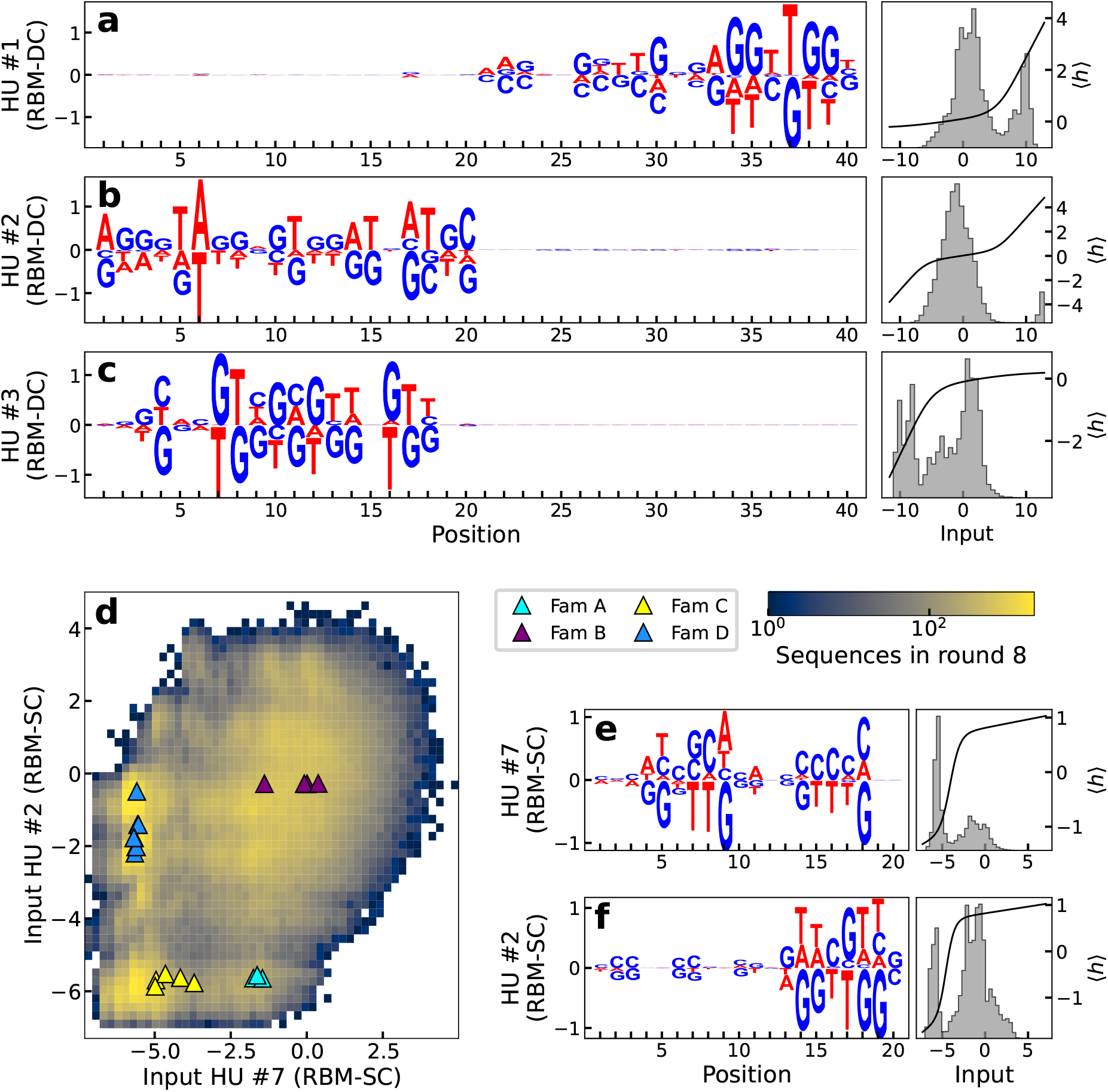
Weights learned by the RBMs have biological interpretations. **a-c**: Left: logos of three weights *μ* = 1, 2, 3 with largest Frobenius norms for RBM-DC trained on round 8 aptamer data; Right: histograms of the inputs *I_μ_* = ∑_*i*_ *w_μi_*(*s_i_*) (where *w_μi_* is the weight of the connection between hidden unit *μ* and visible unit *i* for nucleotide s¿) to the corresponding hidden units for the sequences **s** in the dataset (gray) and average activity (black). **d:** The four families identified in [55] are separated in different clusters in the two-dimensional subspace spanned by the inputs to hidden units 2 and 7 of RBM-SC (trained on loop subsequences at round 8). **e, f**: Logo, distribution of inputs and average activity of the same hidden units as in panel d.

We first observe that the weights are strongly localized either on the left or the right loop. The lack of correlation between the left and right loop sequences holds for all weights (Suppl. Fig. S18), and is compatible with the additivity of their contributions to the aptamer log-likelihood in Fig. 3c.

A closer look at the sequence-dependence of the logos in Fig. 4a-c shows they are G-rich and match parts of G-quadruplex motifs. For instance, the hidden unit focusing on the right loop in Fig. 4a, is strongly activated when the motif AGGTTGG is present on the L loop in positions 33-39. Other L subsequences lead to much weaker activities (in absolute value), see right subpanel in Fig. 4a. A similar observation holds the left loop in Fig. 4c, with the motif GNNTGGTGTGGNTGG in positions 4-18 which is compatible with a G-quadruplex structure. Other features are also detected by the RBM. As an example the weight logo in Fig. 4b is identifying long-range correlations across positions 1-20 associated consisting in a AT-rich motif and is present in some of the training sequences (see histogram in right subpanel).

We then explore the capability of RBM to provide low-dimensional representation of sequences. Prior experimental work [55] have identified four different families of thrombin-binding aptamers (named A, B, C and D), based on sequence alignment and manual curation. We show in Fig. 4d the value of inputs *I_μ_* of two hidden units of single-loop RBM-SC, ranked 2 and 7 in terms of weight Frobenius norms able to cluster these four families. Each hidden unit’s activity (see Methods) has a bimodal distribution (Figs. 4e,f), and the combinations of these modes identify the four families.

### 2.6 RBM trained from unique sequences generate diverse aptamers capable of binding thrombin

After having established that the RBM log-likelihoods and the fitnesses of the aptamers in our dataset are strongly interrelated, we now use the RBM model to generate new sequences *in silico* (see Methods Sec. 4.2). Note that the number of available sequences at any round, < 10^6^, is much smaller than the number of possible sequences over 20 nucleotides, 4^20^ ≃ 10^12^. Hence, it is a non trivial problem to reconstruct the full likelihood landscape from such undersampled data, and use it to generate new binders.

Sampling RBM-SC trained on round-8 data reveals a lack of diversity in the sampled sequences: all the generated sequences with high log-likelihoods are already present in the dataset (Fig. 5a). RBM-SC rightly assigns high scores to the strong binders present at the end of SELEX procedure, but is unable to generate diverse sequences with high scores (Fig. 5b).

**Figure 5:**
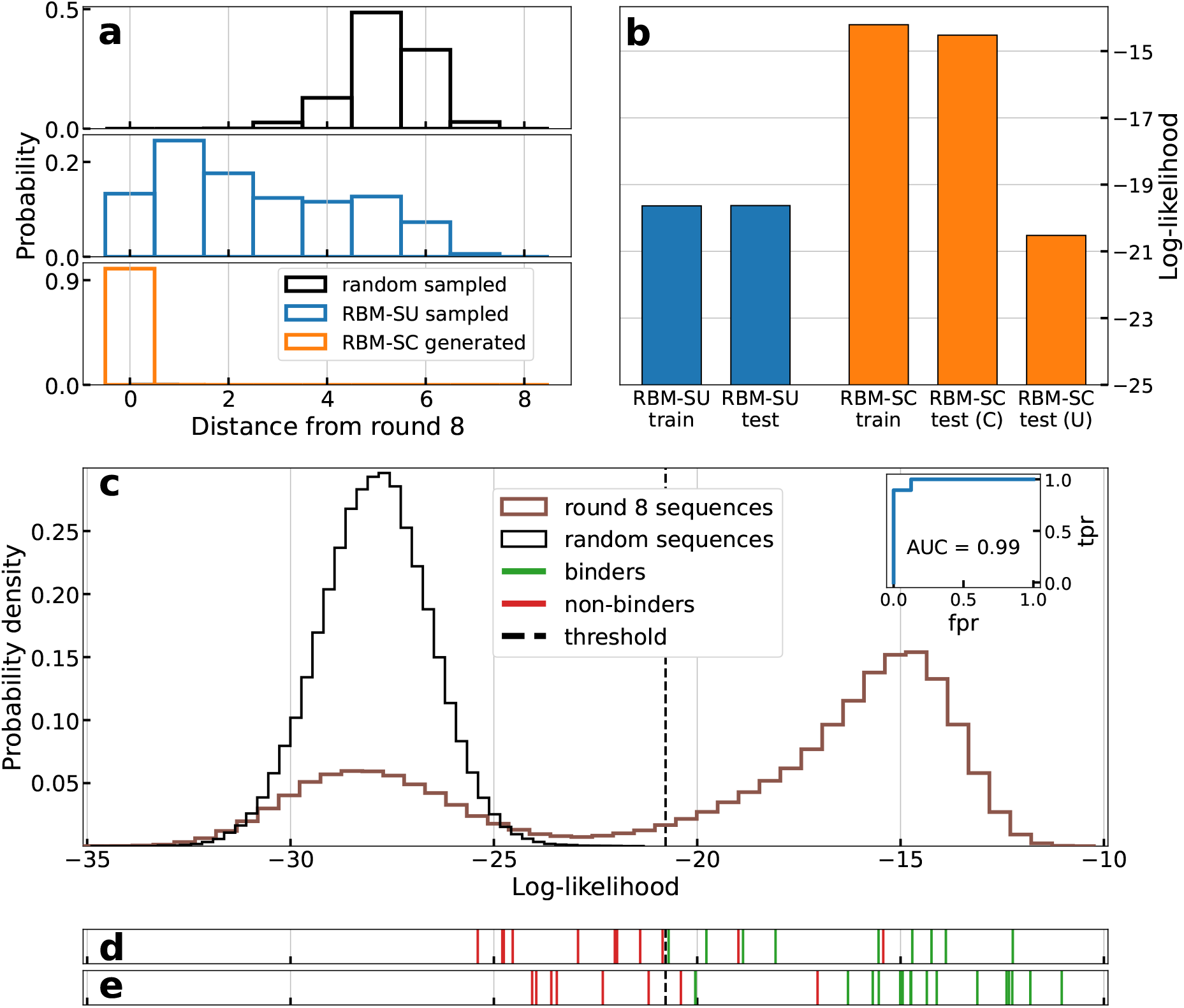
RBM can be used to design new aptamers binding thrombin. **a**: Histograms of distances to dataset of the best (top 5% in terms of log-likelihoods) sequences generated by RBM-SC (orange) and RBM-SU (blue) trained on round 8 data. The black histograms show the distribution obtained with sequences generated uniformly at random. **b**: Average log-likelihoods of training (90% of the unique sequences observed at round 8, chosen at random) and test (remaining 10% of unique sequences) sets for RBM-SU and RBM-SC (after re-introduction of counts). For RBM-SC, the average log-likelihood of the test set is shown when weighting each sequence with (C) or without (U) its counts. **c**: Histogram of the log-likelihoods of all unique aptamers observed in the last round (blue line) and of uniformly random sequences (orange line), computed with RBM-SU trained on the same data (blue line). Inset: AUC computed on the sequences generated by the RBM-SC model (panel e). **d**: Vertical lines locate the log-likelihoods of sequences experimentally validated to be binders (green) or non binders (red). Sequences taken from a preliminary set described in Suppl. Table S5. Results allow us to determine the binding/non binding threshold, located with the black dashed line. **e**: Same as panel d, for sequences designed with the RBM-SU model trained on round-8 sequences, as described in Sec. 2.6, see Table 1.

We then train another model, called RBM-SU, by maximizing the sum of the log-likelihoods of unique sequences in round 8 dataset (composed of 382094 unique single-loop sequences), see Eq. (1). Details of the training procedure are given in Suppl. Sec. S3. The rationale for this approach is two fold. First, 8^th^ round data are expected to include better binders and much less parasite sequences than earlier rounds. Second, discarding the sequence counts prevents the model from being dominated by few very good binders to thrombin.

The effective diversity of training data is reflected in the generated sequences from RBM-SU model. A large fraction of sequences generated by RBM-SU with top log-likelihoods are not present in the dataset, contrary to what found with RBM-SC, see Fig. 5a. In addition, about 30% of generated sequences are 4 or more nucleotides away from the dataset, as is the case for the majority of randomly generated sequences of length 20 nucleotides. Furthermore, we show in Fig. 5b that RBM-SU exhibits excellent generalization properties. The log-likelihood of test data (unique sequences present at round 8 but not used for training) is very close to the one of the training data. On the contrary, RBM-SC essentially assigns high scores to high-count sequences in the training data, and shows poor generalization.

We have next experimentally tested the binding to thrombin of some aptamer sequences to validate the ability of the RBM-SU to predict binding and to generate *de novo* binders. The 20-nucleotide DNA sequences are first inserted into the loop of a hairpin with fixed 18 base-pair-long stem. To estimate the binding affinity to thrombin we use native gel shift assay, where we incubate the thrombin protein with the hairpin aptamer, see Methods Sec. 4.4 and Supp. Inf. Sec. S1.

A set of 16 sequences listed in Suppl. Table S5 (excluding the control sequences listed in the Table), together with 4 binders, experimentally validated in [55] and named ThA, ThB, ThC and ThD, is first used to estimate the log-likelihood threshold above which a sequence is predicted to bind thrombin, see Fig. 5d, where the log-likelihoods of tested sequences are represented as vertical red and green lines, for verified non-binders and binders respectively.

We then propose a set of 27 sequences to test (r1-r27 in Table 1): 2/3 of them are de novo designed sequences generated from the RBM-SU model, and the remaining 1/3 are present in round 8. *De novo* sequences are chosen to test the power of the RBM model to produce good thrombin binders, or to predict critical mutations transforming binders into non binders. Sequences already present in the round-8 data are chosen to test non trivial RBM predictions, *e.g.* sequences with low or high counts having, respectively, high or low log-likelihoods. The detailed description of these sequences and of the design criteria is found in Method Sec. 4.3.

**Table 1:**
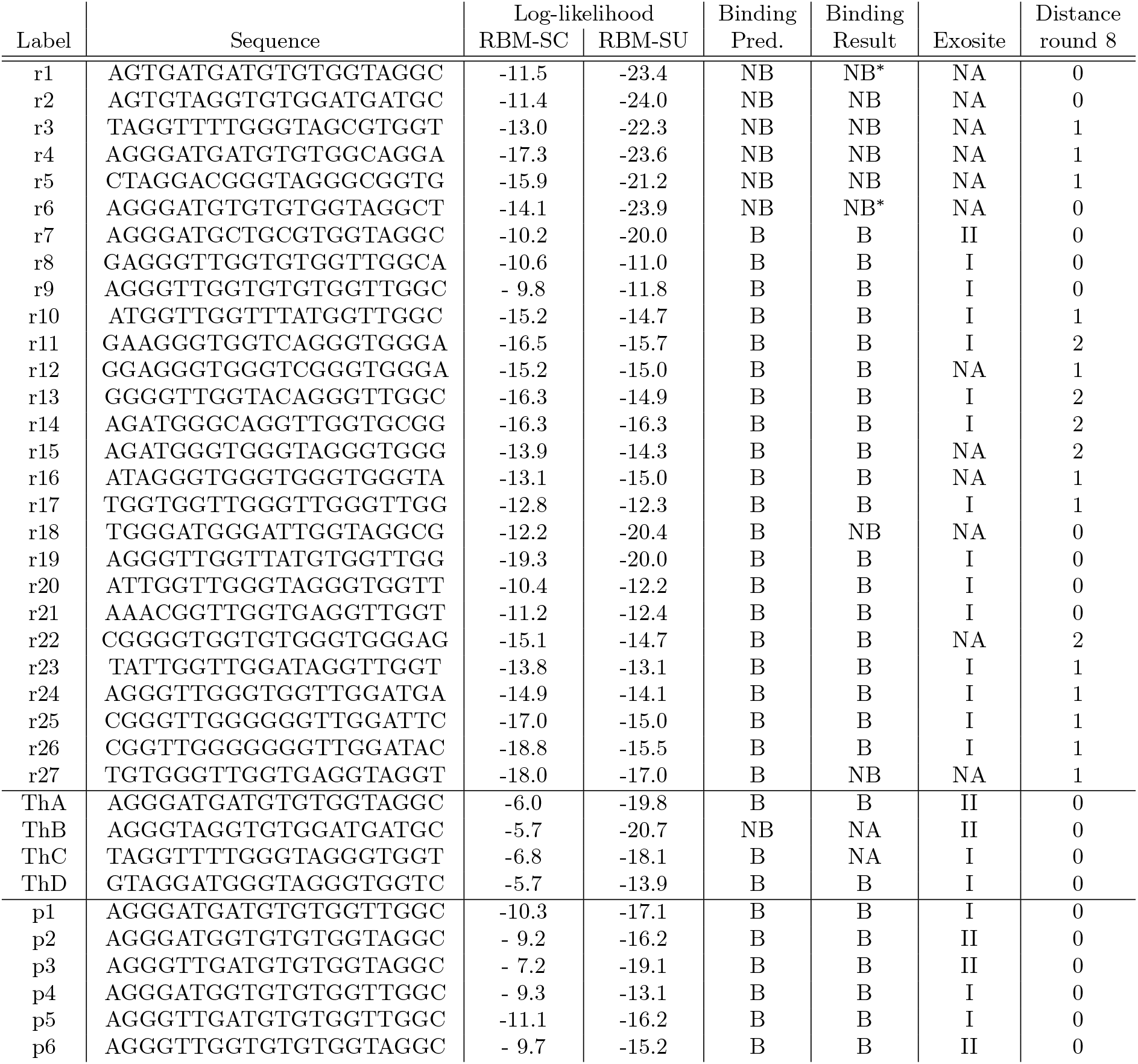
Sequences generated from RBM-SU, log-likelihoods, binding predictions (based on the comparison of the RBM-SU log-likelihood and the threshold in Fig. 5b), and results from gel shift assay (B for binders, NB for non binders) and exosite binding assays. For comparison, data for ThA, ThB, ThC and ThD sequences from Ref. [55] are shown. ThB and ThC have not been tested for binding with our experimental setup (so NA is used in the corresponding column), although they are expected to bind thrombin given the results obtained in Ref. [55]. * Aptamers and r1 and r6 did not show thrombin binding gel band, but their pattern indicates a possible weak interaction with thrombin.

Over the 27 sequences to test, 21 sequences were above threshold, and therefore predicted as binders and 6 sequences below threshold, predicted as non binders. The experimental gel assays are shown in Fig. 6. Overall, 93% of the RBM predictions (binder or non binder) are confirmed by experiments. The log-likelihoods of the tested sequences, along with the RBM predictions and the experimental findings are reported in Table 1 and represented with the experimental results in Fig. 5e.

**Figure 6:**
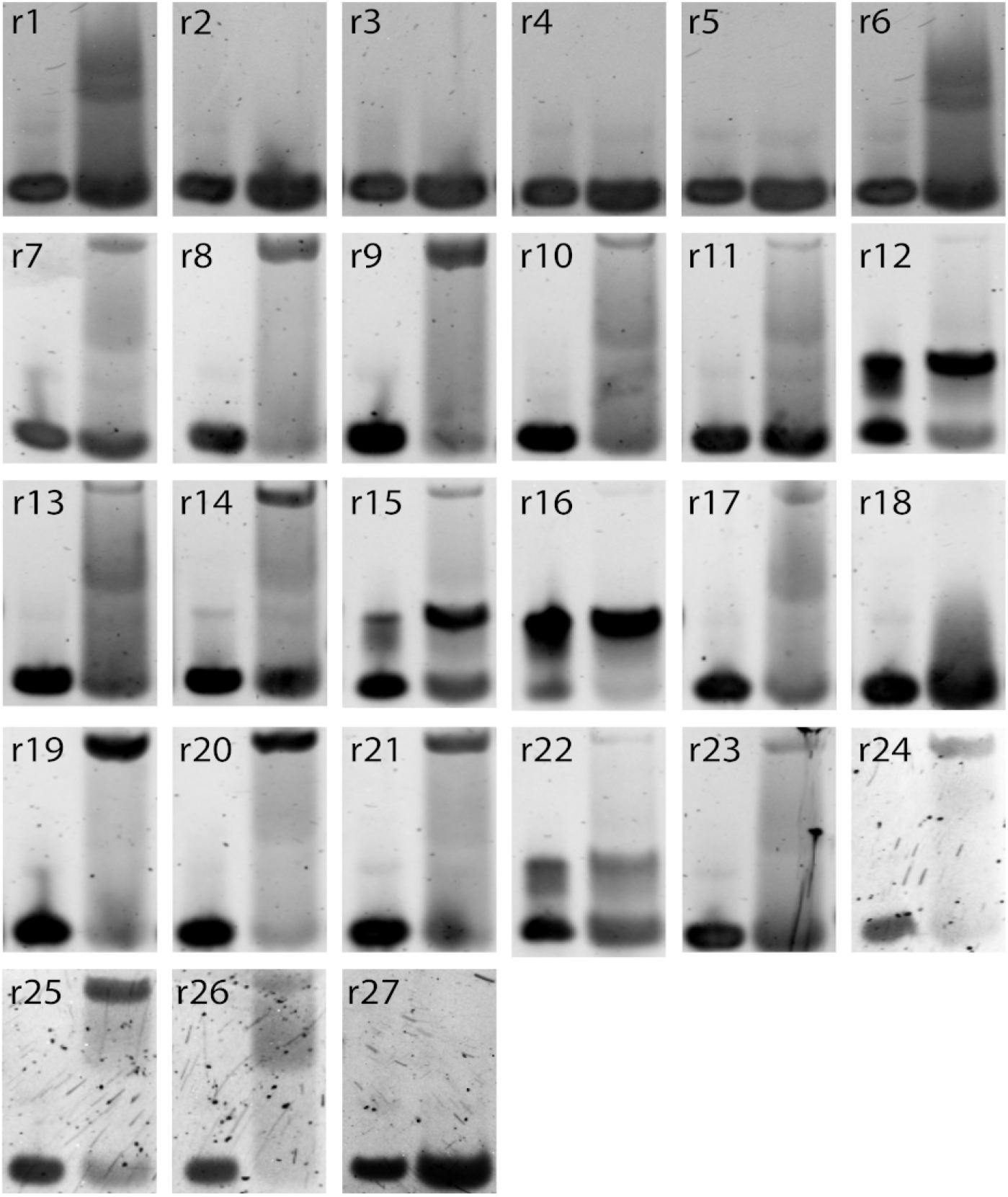
Experimental measurements of binding of respective designed sequences (r1 to r27) to thrombin. 5% native gel assay at 15 °C of stem loops (1-27) alone in the presence of Mg^2+^/K^+^ (lane 1) and allowed to mix with *α*-thrombin for 30 minutes at 25 °C on the bench (lane 2). r12, r15, r16, and r22 aptamers were forming dimers with themselves but upon using samples without K^+^, they were found to bind thrombin. Their entries above display the successful attempt(see Methods for further details). Aptamers r1 and r6 did not show a clear upper band that is indicative of thrombin-aptamer dimer, but the observed smear might indicate weak interactions with the thrombin.

These results show that the log-likelihood provided by RBM-SU is an accurate predictor of the capability to bind thrombin. We show in the inset of Fig. 5c the receiver operating characteristic (ROC) curve and the corresponding area under the curve (AUC=0.99) for RBM-SU-generated sequences. Let us stress that RBM-SC, shows poor performance in discriminating good from bad binders among these sequences, see Suppl. Fig. S26. This failure is expected from the poor generalization abilities of RBM-SC for sequences with low counts (Fig. 5b).

### 2.7 Competition assay for exosite binding site and binding strength measurements

Thrombin has two exosites, referred to as I and II, which can be bound by aptamers, *e.g.* ThA is known to bind exosite II, while ThD binds exosite I [55]. We first identify the target exosite for all the binding aptamers among the r1-r27 by testing each of them (aside from those which were found to form dimer states, see Table 1) against ThA and ThD, see Methods Sec. 4.5. In such a competition assay, the designed aptamers are preincubated with thrombin and are put in competition with a small amount of fluorescently labelled ThD or ThA [55]. If the pre-incubated and fluorescent strand bind the same exosite a thrombin/fluorescent strand complex is observed in the same position as in the thrombin binding assay. However, if the pre-incubated and fluorescent strands target different exosites thrombin is bound twice, causing a downward shift in the observed band (Suppl. Fig. S2). As shown in Fig. 7 and Table 1 we find that all thrombin-binding aptamers among sequences r1-r27 bind exosite I, except one.

**Figure 7:**
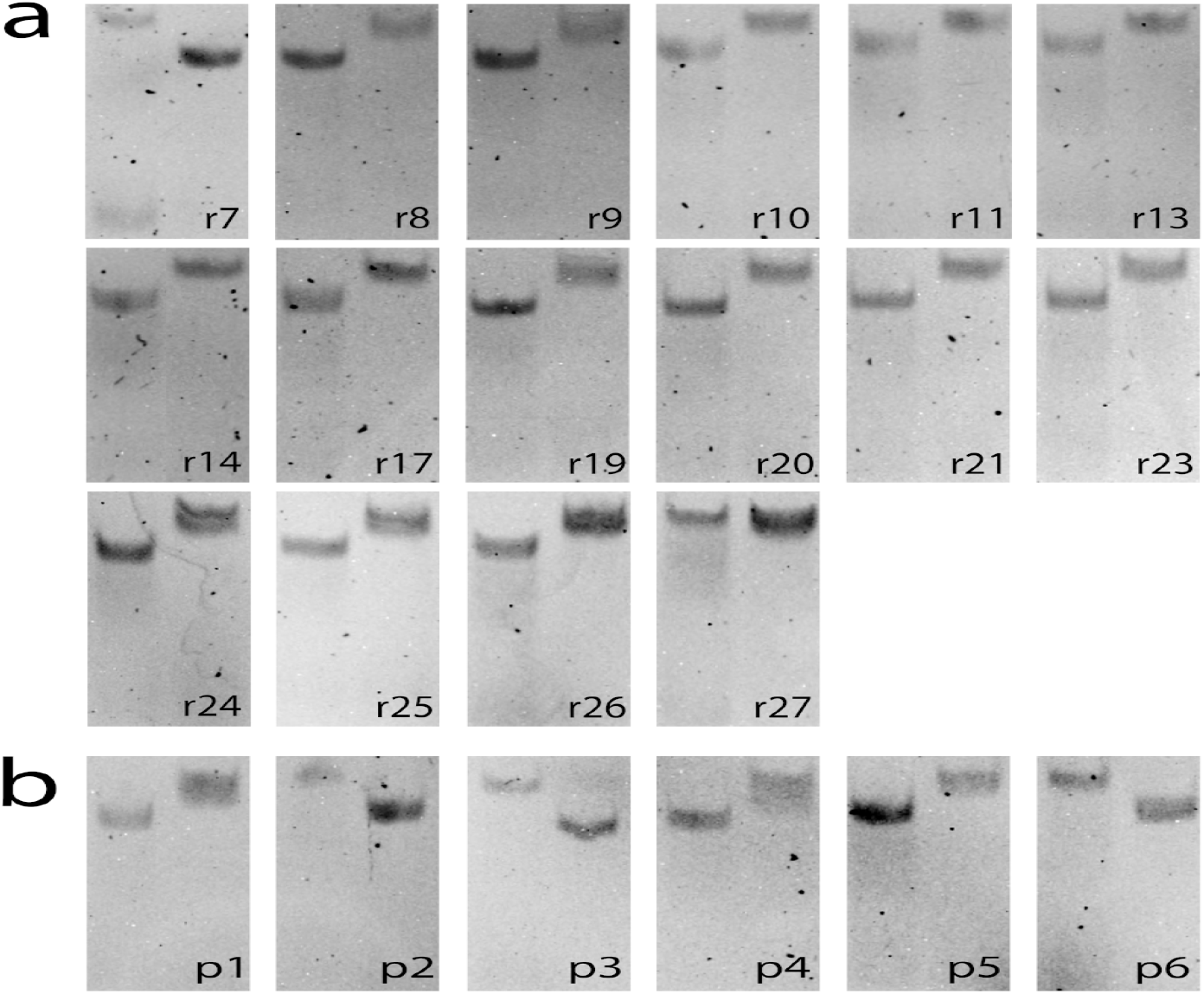
**a**: Binding site assay of all binding sequences and r27 (a nonbinder control) in the RBM generated dataset. **b**: Binding site assay of the 6 sequences that make up the sequence space between ThA and test sequence r9. For all gels, Lane 1 shows addition of ThA and lane 2 shows addition of ThD to the thrombin preincubated strand (labeled in black). Results are reported in Table 1.

As we noticed that sequence r9, which is an exosite-I binder, is only 3 mutations away from ThA, which binds exosite II, we decided to test all six intermediate sequences, labelled as p1-p6 in Table 1. One mutation (Adenine vs. Thymine on site 17) seems to control the exosite binding preference along the mutational path, see Table 1 and Suppl. Fig. S25. Analysis of the RBM-SU weights confirms that position 17 is particularly relevant on the aptamer sequence: many weights have non-zero values on this site (Suppl. Fig. S21). To understand if the presence of A on site 17 (rarely encountered in round 8 sequences) is sufficient to guarantee binding to exosite II we specifically design four sequences (r24 to r27) with this feature and log-likelihoods above threshold, see Methods 4.3 and Table 1. As reported above none of these sequence turns out to bind exosite II (while 3 out of 4 bind exosite I), showing that binding specificity is generally controlled by multiple-nucleotide motifs along the sequence.

Next we test if any of the *de novo* generated aptamer sequences with high RBM-SU log-likelihoods are stronger binders than previously identified ThD and ThA aptamers, the binders with the largest number of counts at the end of SELEX [55]. To determine the strongest binder using competition assays, thrombin is mixed with a mixture of the control and the test aptamers at equal ratios, with the control strand being fluorophore labelled (details in Supp. Mat. Sec. S1.4). The stronger binder is considered to be the control or the test aptamer when fluorescence is observed, respectively, in the thrombin-aptamer gel band or in the stem loop band (the unbound aptamer), see Suppl. Figs. S4 and S3. We observe that none of the designed aptamers binds thrombin more strongly than ThA to exosite II binders, or than ThD to exosite I binders. This result is expected: given the size of the original library (~ 10^15^) virtually all possible sequences of 20-nucleotide aptamer are initially present, so it is unlikely that SELEX misses stronger binders than ThA and ThD.

We then ask whether the outcomes of competition assays for the best binders could be predicted from the comparisons of their log-likelihoods. RBM-SC-based predictions have 100% success with respect to the above competition assays, always assigning larger scores to ThA and ThD than to the competing aptamers. Conversely, RBM-SU underestimates ThA and ThD binding strength, assigning, in particular, low log-likelihood to ThA and having a global performance of 38% on performance of RBM-generated sequences in the competition assays with ThA and ThD. However, for competitive assays between sequences r1-r27, RBM-SU scores are slightly more predictive than their RBM-SC counterparts, with fractions of successful predictions equal to, respectively, 67% and 59%. Interestingly RBM-SU and RBM-SC also depart from one another in their estimates of the log-likelihoods of exosite I and II binders. We observe in Fig. 8 that aptamers binding exosite I have higher scores than their exosite II counterparts, explaining the overwhelming presence of exosite I binders among RBM-SU generated sequences. On the opposite, RBM-SC generally assigns higher log-likelihoods to exosite-II binders. The differences in the behaviours of these models are further examined in Discussion.

**Figure 8:**
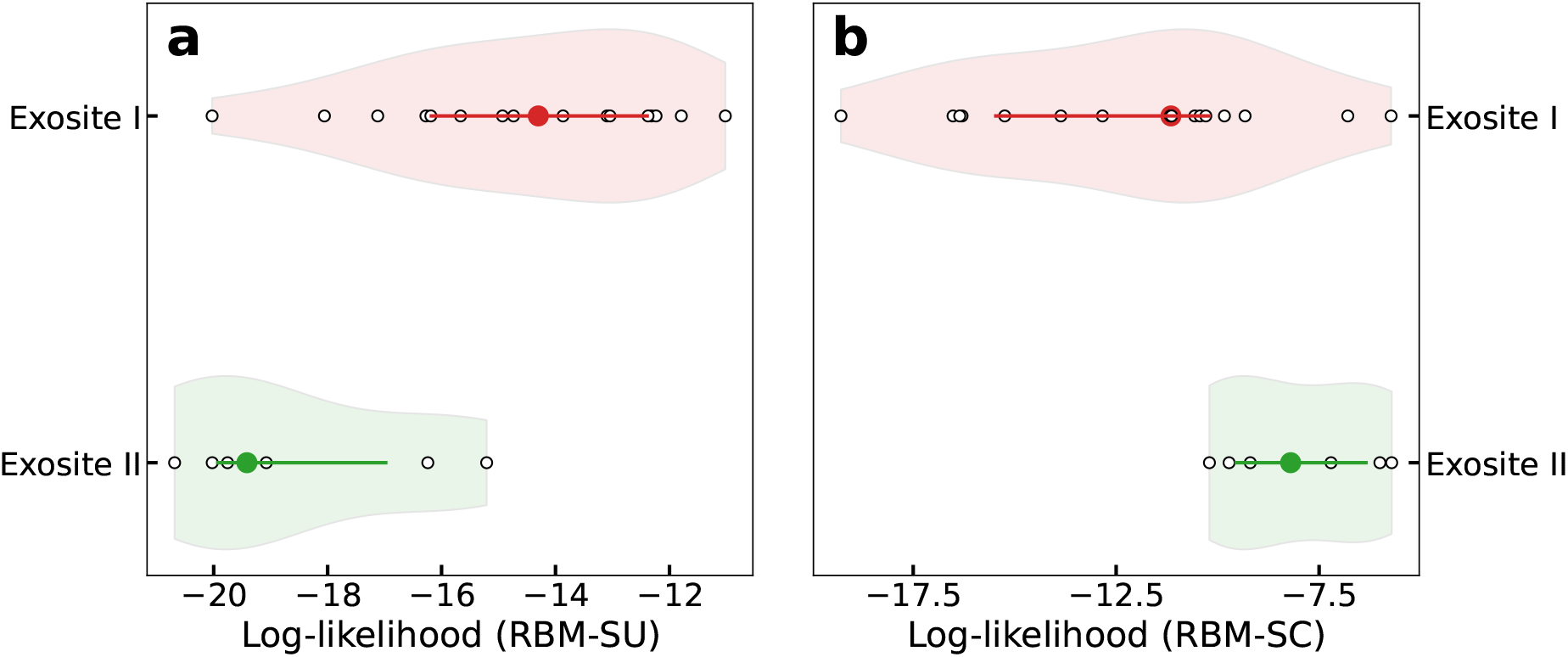
Aptamers binding to exosite I have larger log-likelihoods with RBM-SU, lower log-likelihood with RBM-SC. Violin plots showing the log-likelihoods of exosite I (light orange violin) and exosite II (light green violin) binders. Circles in darker colors denote the average log-likelihood over the class, lines denotes 25- and 75-percentiles, and white points corresponds to the log-likelihoods of the generated sequences. In panel a RBM-SU is used, while in panel b RBM-SC is used.

### 2.8 Supervised learning approach

We also explored supervised learning approaches to train from the aptamer datasets. We considered several DNN architectures (ResNet, Siamese Network and variational autoencoder) as well as traditional methods (random forest and gradient boosted tree) that we trained to classify sequences as binders or non-binders (see Suppl. Sec. S4). Training was complicated by the fact that the aptamer dataset only contained positive examples (binders from different selection rounds with their respective counts obtained from the sequencing step). Hence, we either classified sequences with low counts as non-binders, or we generated random sequences not present in the dataset and treated them as non-binders. The first approach achieved between 70% to 84% accuracy on the validation dataset. The second approach had at least 99% accuracy on the validation dataset for all models. However, when evaluating models against the test set (sequences from Table 1), we observed 30% to 74% accuracy for the first approach, and 70% to 89% for the second approach, as the test set is heavily biased to binding sequences, and methods with high accuracy classified most non-binders as false positives. These results indicate that SELEX datasets are challenging for the commonly used supervised learning methods.

## 3 Discussion

In this work we proposed data-driven models of aptamer sequences obtained at different stages of directed evolution for thrombin binding. Our models are based on Restricted Boltzmann Machines (RBM), the simplest neural network architecture embedding the notion of representation (or latent factors) of sequence data.

One of our main findings is that the score (log-likelihood) assigned by the model to a sequence ***s*** was linearly related to its fitness *F*(***s***) in the SELEX experiment. More precisely, repeated applications of Eq. (1) at previous rounds of selection imply that the likelihood of a sequence ***s*** at round r is related to its fitness through

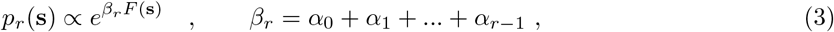

where *α*_*k*–1_ is the intensity of selection from round *k* – 1 to *k,* see Eq. (1), and the initial library is assumed to be roughly uniform over the sequence space (*β*_0_ = 0). This equation can be conveniently rephrased in the language of statistical physics. The rounds of SELEX selection shape a Boltzmann-like distribution over the aptamer sequences, corresponding to an effective energy equal to minus the fitness, –*F*. The effective inverse temperature *β_r_* at round *r* is the sum of the intensities of selection at the previous rounds, and measures the cumulative effect of these previous selections. As more rounds are carried out, the effective temperature 1/*β_r_* diminishes, and the distribution of sequences concentrates around the fittest aptamers, *i.e.* the sequences ***s*** maximizing *F*(***s***), see Fig. 9. As more and more rounds *r* of SELEX are applied to the aptamer population the cumulative selection strength *β_r_* seem to saturate, a phenomenon compatible with previous theoretical works [15] and observed in other SELEX experiments [54].

**Figure 9:**
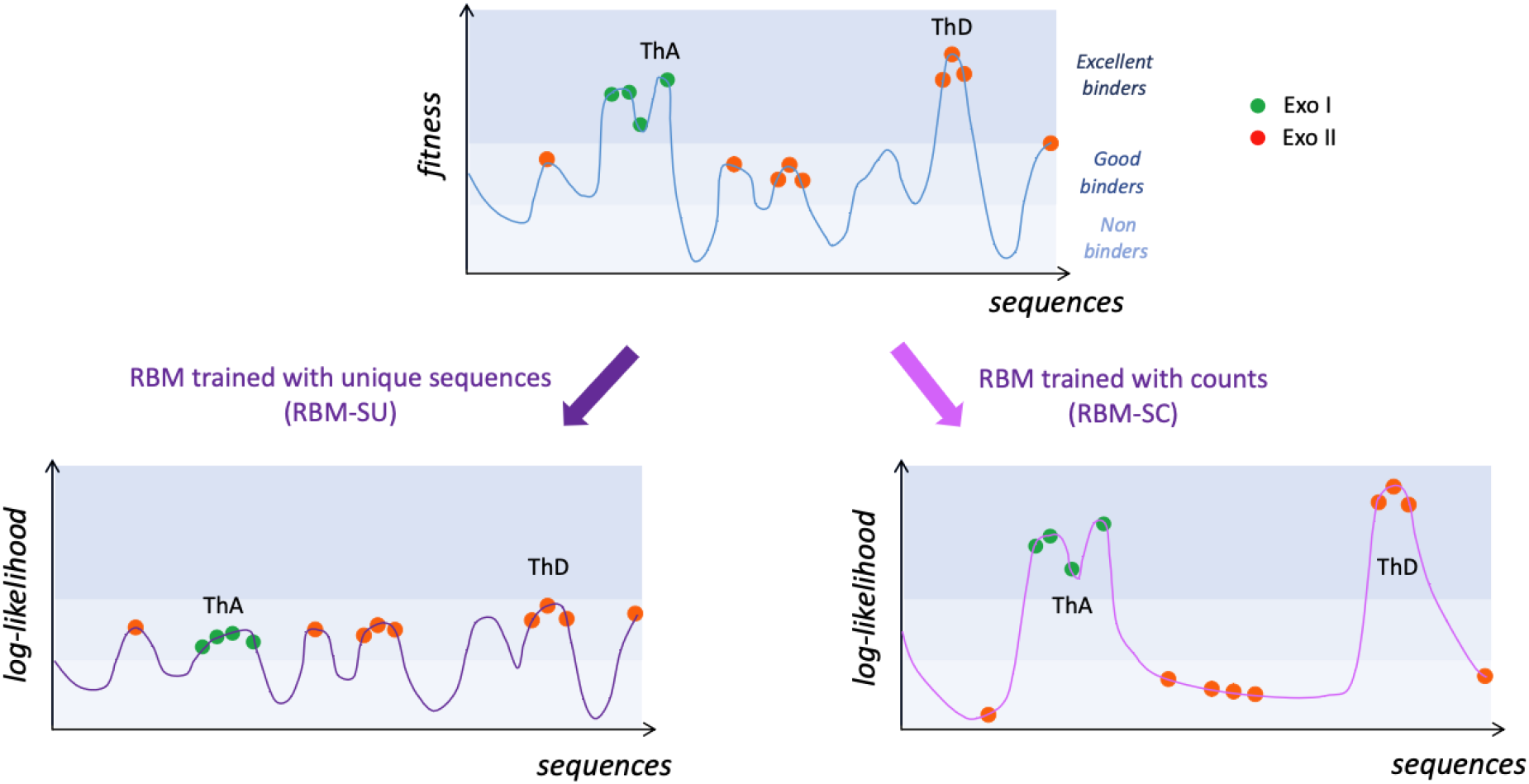
Sketches of the fitness and inferred landscapes. Top: fitness of the aptamer sequences as estimated by the SELEX experiment. After some rounds of selection, most sequences are good binders to thrombin and have low counts (very often, *C* =1), while some are excellent binders and have large counts. Two excellent binders, ThA and ThD, are schematically shown. Bottom: log-likelihood landscapes defined by the RBM models, trained from unique sequences (RBM-SU, left) or taking into counts (RBM-SC, right). RBM-SU is able to capture the statistical features of the many good binders, but does not reproduce well the few high-fitness peaks. It can be used to generate new sequences (empty peak in the landscape). Conversely, RBM-SC accurately models the high peaks in the fitness landscape, but is unable to reproduce the detailed structure of the landscape at lower levels. It cannot be used to generate new binders.

The values of the selection strengths *α_r_* and of the cumulative selection strengths *β_r_* can be extracted from our analysis; for definiteness we arbitrarily choose *β*_6_ = 1 to fix the scale of the fitness *F*, as Eqs. (2) and (3) are obviously unchanged under the rescaling *α_r_, β_r_* → λ *α_r_, β_r_*, *F* → *F*/λ. First, we report in Suppl. Fig. S24 the scatter plots of the log-likelihoods of the sequence data with models trained at different rounds, say, *r* and *r*′; the slopes of these scatter plots give access to the ratios *β_r_/β_r′_* according to Eq. (3). Second the linear fits of the log-likelihood (estimated with the RBM trained on round-6 data) vs. log. enrichment ratios, as well as the Fisher ratios shown in Fig. 2c provide estimates of the ratios *α_r_/β*_6_. The outcome for *α_r_, β_r_* can be found in Suppl. Table S8, and allows us to accurately quantify the amount of selection on the aptamers throughout the SELEX process.

The double-loop nature of the aptamer sequences studied here is at the origin of two interesting phenomena. First, we find that log*p_r_*(***s***) and, consequently, the fitness *F*(***s***) are, to a very good accuracy, equal to the sum of two contributions coming from the left and from the right loops. This additivity property suggests a mechanistic picture of the binding of aptamers to thrombin. The enrichment factor of the set of molecules carrying the sequence ***s*** is proportional to the probability *p*_bind_ that they bind thrombin and to their amplification factor through PCR. Hence, log*p*_bind_ is proportional to the fitness and addivity of the latter implies that *p*_bind_ is the product of the binding probabilities of the left and right loops. The two loops of aptamers are thus progressively required, through successive SELEX rounds, to bind the thrombin target. While double-loop aptamers with one binding loop and one parasite subsequence exist in early rounds, they progressively disappear (Fig. 2a). The bivalence of aptamers in the final rounds likely reflects the strong selection pressure imposed by SELEX.

The RBM model also allows for identification of the nucleotide motifs in the aptamer sequence that contribute most to the sequence likelihood, or, equivalently, to its fitness. Such motifs are indicative of a G-quadruplex group, a known functional motif in the DNA aptamers that bind thrombin [31]. Other RBM motifs could also allow one to help identify clusters of sequences (subfamilies), investigated in prior works through sequence alignments and manual curation.

A second major finding is that the RBM model is capable of generating new sequences, not present in the dataset, with good binding properties. We have generated 27 aptamer sequences from the RBM that were either predicted to bind or not bind to thrombin. Out of 21 sequences that were thought to be binders, 19 were confirmed to bind thrombin, and all 6 sequences generated as non-binders were rightly predicted so. These non-binder sequences were generated under the non-trivial constraint to differ as little as possible (in terms of mutated nucleotides) from known good binders.

We stress that the capability of RBM models to generate diverse aptamers crucially depends on how they are trained. Standard training, where the counts of sequences are taken into account result in models giving very high scores to the very best binders in the dataset, but unable to generalize beyond these few sequences (Fig. 5b). On the contrary, discarding the count information and maximizing the log-likelihood of the set of unique sequences produces models with very good generalization properties, and able to design new and diverse binders, as confirmed in the experiments reported above. The choice of considering unique sequence is partially reminiscent of the reweighting procedure used in sequencebased modeling of proteins [8, 47], and allows the inferred log-likelihoods to reflect more accurately the probabilities for sequences with low number of counts, see Fig. 9. Notice that, while unique-sequencebased training could *a priori* be sensitive to sequencing errors we estimate that the probability *ϵ* of misreading a nucleotide is < 10^-3^ (see Methods Sec. 4.1 and Suppl. Sec. S2), in agreement with error rates with next generation sequencing methods [32]. As a result spurious sequences are < 0.5% of all unique sequences in the dataset, and have only marginal impact on the trained model. However, ensembles in other SELEX experiments using modified bases might experience higher sequencing error rates, which our approach would allow to identify and correct for (Methods Sec. 4.1).

The properties of the two models are graphically summarized in Fig. 9. RBM-SC, which takes into account counts, accurately models the high peaks of the fitness landscape, but discards the smaller peaks. It rightly assigns very high log-likelihoods to the excellent binders, such as ThA or ThD. However, at this level of fitness, the diversity of the sequences that can be generated is very poor. Conversely, RBM-SU, captures the statistical features of sequences at a much lower level of fitness. Many varied sequences can then be generated, the majority of which are good binders. RBM-SU is therefore able to generate more diverse and less strong binders, which makes particularly appropriate for the design of evolvable aptamers [50]. In principle, RBM-SC inferred from sequences collected in an early round would have had similar properties to RBM-SU inferred from round-8 data. However, in the specific problem of double-loop aptamers we consider here, the presence of a large number of parasite single-loop sequences at the beginning of SELEX evolution could also affect the generative power of models trained at early rounds.

We next used a competitive binding assay both to first classify the binding site of the generated sequences and, in a second step, to assess the strength of binding to a given exosite. We find that the majority of sequences generated with RBM-SU preferentially bind to exosite I. In addition, sequences binding exosite I have on average higher log-likelihoods than the few exosite-II binding sequences. In particular, ThA, an exosite-II binder with a large number of counts in the SELEX experiment is not among the sequences with highest RBM-SU log-likelihoods. Furthermore direct competition experiments between the highest log-likelihood sequences and ThA or ThD (binding exosite I and having a large number of counts) showed that the latter aptamers outperform the former in terms of binding affinity. These apparently paradoxical results can be explained in two ways. First, the log-likelihoods were estimated with the model used for generating sequences, that is, RBM-SU. This model is very good at generate diverse binders, but is not trained to reproduce counts. The absence of correlation between RBM-SU log-likelihoods and counts or binding affinities is therefore not surprising, whereas RBM-SC high scores show a good correlation with large counts as expected (Suppl. Fig. S23,). Second, these results are compatible with a selection mechanism involving binding to the two sites of thrombin. Binding to exosite II has been shown to facilitate binding to exosite I, presumably through allosteric structural change [55]. Due to this allostery mechanism, when exosite II is loaded (even with a different molecule), hairpin with a low-affinity loop to exosite I could be selected. This mechanism could produce a rather subtle parasitism, where only the best exosite II binders in a quasi-monoclonal population (few sequences with largest counts) are under strong selection, and allow for the presence of a more diverse exosite-I binder population. Further experimental investigations combined with theoretical analysis, *e.g.* using concepts developed in ecosystems dynamics in presence of parasite populations, could help to further investigate the selection dynamics.

Finally, we note that our RBM represents a higher level of complexity than the direct contact analysis methods (DCA) that have also been recently applied to protein ensemble selection experiments [4]. While the DCA method trained using the pseudo-likelihood method was not able to correctly predict binders and non-binders for our dataset, when we used contrastive divergence training for DCA, the assigned scores from the trained DCA model showed correlation with our trained RBM (see Supp. Inf. Sec. S5). As opposed to DCA, which infers pairwise interactions, the RBM model’s hidden units can be used for clustering of sequences or identification of multi-nucleotide motifs, such as G-quadruplexes, making them more readily interpretable. We have also explored using supervised learning models, including DNNs, on our datasets predicting binders and non-binders, but as further detailed in Supp. Inf. Sec.S4, we did not obtain good prediction accuracy for the outcomes of our experiments with designed sequences.

We anticipate that RBMs will be also useful for the modeling of other aptamer datasets, including competition assays where aptamers are selected to bind to a desired target, *e.g.* cancerous tissues, and at the same time not to bind to the control, *e.g.* healthy tissue. We believe our approach has the potential to generate alternative or better binders for these complex targets, as well as to unveil the sequence motifs that are enriched or avoided in these high-quality aptamers. The same approach can be also useful to model RNA and DNA regulatory sequences and their interaction with proteins in the key processes such as transcription regulation [52, 26, 56, 23]. Lastly, our modeling and design methods are also readily applicable to other selection-amplification protocols, such as phage display for antibody discovery [14, 24] or directed protein evolution studies [4, 42], which have much larger space of possible sequences (20^*L*^ for length *L*) compared to aptamers (4^*L*^).

## 4 Methods

### 4.1 Estimation of sequencing error probability

Sequencing errors are potentially harmful, as they could lead to more unique sequences in the dataset and possible biases in the RBM models. We introduce an inference approach to estimate the sequencing error rate, based on the presence of spurious single-site mutations of sequences with high number of counts. In practice the method consists in selecting a subset of sequences with high number of counts, referred to as “peak” sequences, and in comparing the expected number of sequences one mutation away from these peaks due to sequencing errors to the actual number in the data. Our analysis, detailed in Supplementary Sec. S2, indicates that the error rate (per nucleotide) is smaller than *ϵ*\* ~ 10^-3^.

We use this bound to estimate the expected number of spurious sequences present in the dataset. We obtain *N*_spurious_ ~ 1000 unique sequences (see Supplementary Sec. S2), corresponding to ~ 0.5% of the total number of unique sequences present in the data.

### 4.2 Restricted Boltzmann Machine: definition, training, sampling

The probability of a visible and hidden units state in an RBM model is defined by

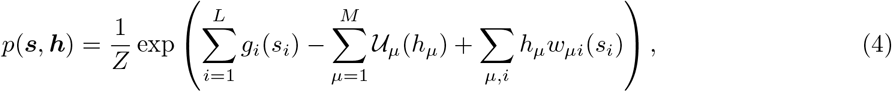

where *Z* is the normalization, *g_i_*, and *w_μi_* are parameters to be inferred from the data during training, and

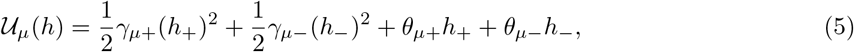

where *h*_+_ = max(*h*, 0), *h*_–_ = min(*h*, 0) and *γ_μ+_, γ_μ–_, θ_μ+_, θ_μ–_* are again model parameters to be inferred from the data during training. This specific form of the function 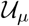, which is called “double Rectified Linear Unit” combines the usage of a relatively low number of parameters with the possibility of learning high-order correlations in the data [47]. An advantage of choosing Double ReLU potentials is that the likelihood log*p*(***s***) of a sequence ***s***, obtained by marginalizing *p*(***s, h***) over ***h***, has an explicit analytical expression in terms of error functions.

It has been suggested that, for RBMs, sparsity of the weight parameters, together with a high number of hidden units, can improve the generative properties of the machine and its interpretability [48, 47]. We resort to a 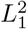 regularization scheme, which consists in adding to the log-likelihood of the data, 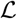 in Eq. (1), a term of the form [47, 5]

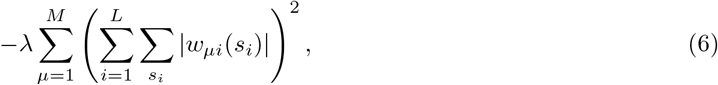

hence enhancing sparsity homogeneously across hidden units. The value of the hyperparameter λ must be, in general, chosen carefully to balance model interpretability (obtained for sparse weights, *i.e.* large λ) and expressivity (to learn data features). We observed little effects of changes in hyperparameters (see also Suppl. Fig. S22), provided that they are not too different from the one given in Suppl. Sec. S3. This is also the case for the number M of hidden units chosen: we used *M* ≃ 70 for RBMs with *L* = 20 visible units, and *M* ≃ 90 for *L* = 40. Precise values of *M* are given, for each RBM used, in Suppl. Sec. S3, but we noticed that using different numbers have little effects on the results discussed in this work (see also Suppl. Fig. S22).

Once the parameters in Eq. (4) are obtained, we can sample from the marginal distribution *p*(***s***) to generate new sequences. Sampling can be done in several ways [38]. Here we use alternate Gibbs sampling (AGS), which consists in sampling the RBM’s visible units while keeping the hidden units fixed and vice-versa, in an alternate manner, until the Monte Carlo Markov Chain equilibrates. To increase the probability of sampling high log-likelihood sequences we can sample from *p*(***s***)^2^ instead of *p*(***s***) using the so-called duplication trick [47]. We write

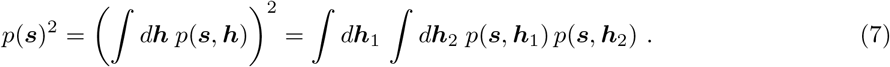

This squared likelihood distribution can therefore be sampled with standard AGS after duplication of the hidden layer of the trained RBM model.

The average hidden unit *μ*’s activity for a given sequence **s** is defined as 〈*h_μ_*) = ∫ *d***h** *h_μ_ p*(**h**|**s**). Note that 〈*h_μ_*〉 only depends on the sequence **s** through the input *I_μ_* = ∑_*i*_ *w_μi_*(**s**_*i*_). When the average activity is close to 0, the corresponding hidden unit has vanishing contribution to the sequence log-likelihood, while for both large negative or positive values of average activity the contribution of the hidden unit to the log-likelihood is positive. Therefore the sign of the weights *w_iμ_* assigned to a particular sequence motif is not indicative itself of the presence or absence of a given pattern, as the contribution in *p*(**h**|**s**) depends on the product *h_μ_I_μ_* and can only be null or positive.

### 4.3 Design of single-loop aptamers with RBM

The RBM-SU distribution *p*(***s***) can be sampled to generate sequences ***s*** of interest, and test the validity of the model. We describe below how we generated sequences in Table 1.

#### Determination of threshold

We fix the threshold, which allows us to distinguish good from bad binders based on their log-likelihoods to minimize the number of misclassified sequences among the preliminary set of sequences given in Suppl. Table S5. As a range of possible values are possible, we actually take the median of this interval.

#### Sequences with high likelihoods

We first sample through AGS (see Sec. 4.2) 4000 sequences from *p*(***s***) and from *p*(***s***)^2^. We then choose 10 among these sequences (named r9 to r17 and r22, r23 in Table 1), which have both high log-likelihood and large distances (numbers of different nucleotides) to round 8 data. In practice these sequences are at Hamming distance 1 or 2 from the closest sequences in the original dataset, since further away sequences have substantially lower log-likelihoods. All 10 generated sequences are experimentally confirmed to be good binders (Table 1), and are indicated as green lines in Fig. 5c.

#### Sequences with critical mutations for binding/non-binding status

We next use our RBM to predict critical mutations capable of changing the binder/non-binder status of aptamers. First we exhaustively look for the smallest possible number of mutations leading to a substantial decrease of the log-likelihood of known good binders. In particular, sequence r1 has 1 mutation with respect to a control sequence that we tested for binding (named d10 in Suppl. Table S5), r2 and r3 are both 1 mutation away from, respectively ThB and ThC, both identified as good binders in Ref. [55]. All these generated sequences are confirmed to be unable to stably bind thrombin after this single-point mutation (Table 1 and Fig. 6) and they correspond to red vertical lines to the left of the threshold in Fig. 5c. All these mutations removed a G from the sequence, and G nucleotides are necessary to form G-quadruplex motifs, known to be important for thrombin aptamers. To show that our RBM can also identify other positions in the aptamer that are key to thrombin binding, we also design two more sequences, r4 and r5, which have 2 mutations with respect to aptamers found in the SELEX dataset and validated as good binders (respectively, d10 and d18, see Suppl. Table S5). The mutations are again chosen so that the log-likelihood is decreased as much as possible, but without removing G nucleotides from the original sequences. We find the sequences lost their ability to bind thrombin after the 2 mutations, as predicted by the RBM (Fig. 6), so they correspond to two vertical red lines to the left of the threshold in Fig. 5c.

#### Sequences in dataset with mismatches between counts and log-likelihoods

We further test the performance of the RBM model by searching for sequences with (1) relatively low log-likelihoods but with large numbers of counts (139 or more, see Suppl. Tab. S7) in the SELEX experimental data from Ref. [55], of for sequences with high log-likelihood but with few counts (11 or less, see Suppl. Tab. S7). The sequences chosen in case (1) are r6, r7, r18, r19 (see Table 1); one of them (r6) is below, and the other 3 are slightly above the identified log-likelihood threshold. Sequences chosen in case (2) are r8, r9, r20, r21 (Table 1). The RBM predictions are confirmed in all cases but one (r18), which corresponds to the red vertical line at the right of the threshold in Fig. 5c.

#### Sequences sharing a rare mutation with ThA, a strong exosite-II binder

Last of all we design de novo sequences (r24 to r27 in Table 1) under the following two-fold criterion. First these sequences are required to have Adenine in position 17, which is uncommon in the training dataset (A is the second least common nucleotide in that position, being present in about 13% of the sequences in round 8; it is found in ThA, which strongly binds exosite II). Second, the sequences are required to have large log-likelihoods, exceeding the threshold value. Remarkably, the only non-binder among r24-r27 is the one with lowest log-likelihood, r27. However, while mutating away from A in ThA change the binding specificity from exosite II to I (Fig. 8) sequences r24 to r27 are all exosite-I binders, showing that the presence of A17 is not sufficient for exosite-II specificity.

### 4.4 Thrombin binding assay

All RBM designed sequences were first assessed for their ability to bind either of the cationic exosites of human alpha-thrombin. Each sequence was placed as the loop of a 18 bp stem loop with the full sequences reported in the Suppl. Table S1. As done previously [55], we used a 5% native gel shift assay to qualitatively assess the binding of each stem loop to thrombin. Each sequence was tested with two gel lanes, the first lane always corresponding to the stem loop without thrombin and the second lane consisting of equimolar amounts (500 nM) of thrombin and the stem loop. The presence of an upper band, consisting of a stem loop bound to thrombin complex, in the second lane indicates a binding sequence. Sequences without the upper band (nonbinding sequences) either very weakly interact with thrombin, characterized by a smear but no band in the second lane, or do not interact with thrombin at all matching their negative control lane. Sequences ThA and ThD were selected from the previous study as positive controls for their high affinity for thrombin and known binding sites [55]. Results for all RBM generated sequences are shown in Figure 6 and summarized in Table 1. Results for all DCA generated sequences are shown in Suppl. Fig. S12 and summarized in Suppl. Table S5. To quantify the interaction of the stem loop and thrombin, we tested both control sequences independently and together in varying concentrations of thrombin (Suppl. Fig. S2). The results clearly indicate the stem loop/thrombin band occurs from a 1:1 interaction of thrombin and each stem loop, and the simultaneous binding of two stem loops on opposite exosites of thrombin downshifts the stem loops/ thrombin band from the singular case.

A secondary band prominently appeared among four of the sequences during the binding assay, (r12, r15, r16, and r22). These sequences showed no binding to Thrombin at first. Upon further investigation, the secondary band was found to most likely be a dimer state of the DNA loop from interaction of the G-quartet motifs. The four sequences have a higher G-content than all other RBM-generated sequences. Additionally, a G-quadruplex dimer would require K^+^ cations to form, indicating a testable transition from the single loop to dimer state. The sequences’ Thrombin binding ability was re-assessed by the same experiment, with two small changes. The first was remaking the DNA samples without K^+^ in their buffer, so their transition from single stem loop to a dimer state could be observed[21]. The second change was the heating the DNA samples to 90 “C for 5 minutes before immediately chilling them in ice. Samples (r12, r15, r16, r22) in Figure 6 show the results of this final experiment, with all dimer-susceptible sequences showing an ability to bind Thrombin. Accordingly, we classify these sequences as binders and suggest their absence from the original dataset is due to G-quadruplex dimer formation during the original SELEX procedure. A clear shift from the monomer state in 1x TAE Mg^2+^ (no K^+^) buffer (lane 1) to the dimer state upon addition of buffer with K+ (lane 2) is also observed for all dimer-susceptible sequences. Note this transition still contains some fraction of the dimer state in lane 1 where the sample contains no K+. This is due to presence of K+ in the gel matrix itself as well as the running buffer.

### 4.5 Exosite binding assay

RBM-generated sequences that were able to bind to thrombin were tested to determine which exosite (I or II) of thrombin they bind to. Each aptamer was pre-incubated with thrombin for 30 minutes at 25 °Cat an equimolar ratio in two separate samples. Small amounts (1/10th the pre-incubated strand) of fluorescent labeled exosite II binder ThA [55] was added to the first sample and fluorescent labeled exosite I binder ThD to the second. Using the same strategy as our thrombin binding assay, our samples were run in a 5% native gel with 5 mM K^+^ for proper DNA/thrombin binding. If the pre-incubated strand bound the same exosite as the fluorescent strand, the thrombin/fluorescent strand complex band would be observed in the same position as seen in our thrombin binding assay. However, if the pre-incubated strand bound the opposite exosite as the fluorescent strand, both strands bind thrombin causing the same downward shift as observed for our exosite verified control strands mixing (Suppl. Fig. S2). Accordingly, sequences with no binding affinity to thrombin matched control samples with no test strand. By comparing the outcome of both lanes for a sample we are able to firmly assign the binding site of our test sequences. The gel results are shown in Fig. 7 and summarized in Table 1.

## 5 Acknowledgement

We thank G. Mayer and M. Famulok for helpful discussions and comments. P.S. and J.P. acknowledge the use of the Extreme Science and Engineering Discovery Environment (XSEDE), which is supported by National Science Foundation grant number TG-BIO210009. We further acknowledge Research Computing at Arizona State University for providing HPC resources that have contributed to the research results reported within this paper. S.C., A.D.G. and R.M. are partially supported by the ANR-17 RBMPro CE30-0021-01 and ANR-19 Decrypted CE30-0021-01 grants.

## 6 Data Availability

The source codes used for training and sampling from RBM, as well as the trained supervised learning models, is available at github.com/adigioacchino/RBMsForAptamers. The data used for this work is publicly available through the Zenodo service, with the following Digital Object Identifier (DOI): 10.5281/zenodo.6341687.

## References

[1] K. K. Alam, J. L. Chang, and D. H. Burke. Fastaptamer: a bioinformatic toolkit for high-throughput sequence analysis of combinatorial selections. Molecular Therapy-Nucleic Acids, 4:e230, 2015.

[2] T. L. Bailey, J. Johnson, C. E. Grant, and W. S. Noble. The meme suite. Nucleic acids research, 43(W1):W39–W49, 2015.

[3] P. Bannigan, M. Aldeghi, Z. Bao, F. Häse, A. Aspuru-Guzik, and C. Allen. Machine learning directed drug formulation development. Advanced Drug Delivery Reviews, 2021.

[4] M. Bisardi, J. Rodriguez-Rivas, F. Zamponi, and M. Weigt. Modeling sequence-space exploration and emergence of epistatic signals in protein evolution. Molecular biology and evolution, 39(1):msab321, 2022.

[5] B. Bravi, J. Tubiana, S. Cocco, R. Monasson, T. Mora, and A. M. Walczak. RBM-MHC: A semisupervised machine-learning method for sample-specific prediction of antigen presentation by hla-i alleles. Cell systems, 12(2):195–202, 2021.

[6] D. H. Bryant, A. Bashir, S. Sinai, N. K. Jain, P. J. Ogden, P. F. Riley, G. M. Church, L. J. Colwell, and E. D. Kelsic. Deep diversification of an aav capsid protein by machine learning. Nature Biotechnology, 39(6):691–696, 2021.

[7] L. Civit, S. M. Taghdisi, A. Jonczyk, S. K. Haßel, C. Gröber, M. Blank, H. J. Stunden, M. Beyer, J. Schultze, E. Latz, et al. Systematic evaluation of cell-selex enriched aptamers binding to breast cancer cells. Biochimie, 145:53–62, 2018.

[8] S. Cocco, C. Feinauer, M. Figliuzzi, R. Monasson, and M. Weigt. Inverse statistical physics of protein sequences: a key issues review. Reports on Progress in Physics, 81(3):032601, 2018.

[9] E. De Leonardis, B. Lutz, S. Ratz, S. Cocco, R. Monasson, A. Schug, and M. Weigt. Direct-coupling analysis of nucleotide coevolution facilitates RNA secondary and tertiary structure prediction. Nucleic acids research, 43(21):10444–10455, 2015.

[10] V. Domenyuk, Z. Gatalica, R. Santhanam, X. Wei, A. Stark, P. Kennedy, B. Toussaint, S. Levenberg, J. Wang, N. Xiao, et al. Poly-ligand profiling differentiates trastuzumab-treated breast cancer patients according to their outcomes. Nature communications, 9(1):1–9, 2018.

[11] S. D’Souza, K. Prema, and S. Balaji. Machine learning models for drug–target interactions: current knowledge and future directions. Drug Discovery Today, 25(4):748–756, 2020.

[12] A. D. Ellington and J. W. Szostak. In vitro selection of rna molecules that bind specific ligands. Nature, 346(6287):818–822, 1990.

[13] J. P. Elskens, J. M. Elskens, and A. Madder. Chemical modification of aptamers for increased binding affinity in diagnostic applications: Current status and future prospects. International Journal of Molecular Sciences, 21(12):4522, 2020.

[14] C. M. Hammers and J. R. Stanley. Antibody phage display: technique and applications. The Journal of investigative dermatology, 134(2):e17, 2014.

[15] D. L. Hartl, D. E. Dykhuizen, and A. M. Dean. Limits of adaptation: The evolution of selective neutrality. Genetics, 111(3):655–674, nov 1985.

[16] G. E. Hinton. Training products of experts by minimizing contrastive divergence. Neural Computation, 14(8):1771–1800, 2002.

[17] J. Hoinka, E. Zotenko, A. Friedman, Z. E. Sauna, and T. M. Przytycka. Identification of sequence–structure rna binding motifs for selex-derived aptamers. Bioinformatics, 28(12):i215–i223, 2012.

[18] T. Hornung, H. A. O’Neill, S. C. Logie, K. M. Fowler, J. E. Duncan, M. Rosenow, A. S. Bondre, T. Tinder, V. Maher, J. Zarkovic, et al. Adapt identifies an escrt complex composition that discriminates vcap from lncap prostate cancer cell exosomes. Nucleic acids research, 48(8):4013–4027, 2020.

[19] P. Jiang, S. Meyer, Z. Hou, N. E. Propson, H. T. Soh, J. A. Thomson, and R. Stewart. Mpbind: a meta-motif-based statistical framework and pipeline to predict binding potential of selex-derived aptamers. Bioinformatics, 30(18):2665–2667, 2014.

[20] J. Jumper, R. Evans, A. Pritzel, T. Green, M. Figurnov, O. Ronneberger, K. Tunyasuvunakool, R. Bates, A. Žídek, A. Potapenko, A. Bridgland, C. Meyer, S. A. A. Kohl, A. J. Ballard, A. Cowie, B. Romera-Paredes, S. Nikolov, R. Jain, J. Adler, T. Back, S. Petersen, D. Reiman, E. Clancy, M. Zielinski, M. Steinegger, M. Pacholska, T. Berghammer, S. Bodenstein, D. Silver, O. Vinyals, A. W. Senior, K. Kavukcuoglu, P. Kohli, and D. Hassabis. Highly accurate protein structure prediction with AlphaFold. Nature, 596(7873):583–589, 2021.

[21] M. Kogut, C. Kleist, and J. Czub. Why do g-quadruplexes dimerize through the 5’-ends? driving forces for g4 dna dimerization examined in atomic detail. PLoS computational biology, 15(9):e1007383, 2019.

[22] P. K. Koo, P. Anand, S. B. Paul, and S. R. Eddy. Inferring sequence-structure preferences of RNA-binding proteins with convolutional residual networks. BioRxiv, page 418459, 2018.

[23] P. K. Koo and M. Ploenzke. Deep learning for inferring transcription factor binding sites. Current opinion in systems biology, 19:16–23, 2020.

[24] T. Kretzschmar and T. Von Rüden. Antibody discovery: phage display. Current opinion in biotechnology, 13(6):598–602, 2002.

[25] S. Lennarz, T. C. Alich, T. Kelly, M. Blind, H. Beck, and G. Mayer. Selective aptamer-based control of intraneuronal signaling. Angewandte Chemie, 127(18):5459–5463, 2015.

[26] T.-F. Lou, C. A. Weidmann, J. Killingsworth, T. M. T. Hall, A. C. Goldstrohm, and Z. T. Campbell. Integrated analysis of rna-binding protein complexes using in vitro selection and high-throughput sequencing and sequence specificity landscapes (seqrs). Methods, 118:171–181, 2017.

[27] G. Mayer, A. Lohberger, S. Butzen, M. Pofahl, M. Blind, and A. Heckel. From selection to caged aptamers: identification of light-dependent ssdna aptamers targeting cytohesin. Bioorganic & medicinal chemistry letters, 19(23):6561–6564, 2009.

[28] F. Morcos, A. Pagnani, B. Lunt, A. Bertolino, D. S. Marks, C. Sander, R. Zecchina, J. N. Onuchic, T. Hwa, and M. Weigt. Direct-coupling analysis of residue coevolution captures native contacts across many protein families. Proceedings of the National Academy of Sciences, 108(49):E1293–E1301, 2011.

[29] R. A. Neher and B. I. Shraiman. Statistical genetics and evolution of quantitative traits. Reviews of Modern Physics, 83(4):1283–1300, nov 2011.

[30] A. D. Ortega, V. Takhaveev, S. R. Vedelaar, Y. Long, N. Mestre-Farras, D. Incarnato, F. Ersoy, L. F. Olsen, G. Mayer, and M. Heinemann. A synthetic rna-based biosensor for fructose-1, 6-bisphosphate that reports glycolytic flux. Cell Chemical Biology, 2021.

[31] K. Padmanabhan, K. Padmanabhan, J. Ferrara, J. E. Sadler, and A. Tulinsky. The structure of alpha-thrombin inhibited by a 15-mer single-stranded dna aptamer. Journal of Biological Chemistry, 268(24):17651–17654, 1993.

[32] F. Pfeiffer, C. Gröber, M. Blank, K. Händler, M. Beyer, J. L. Schultze, and G. Mayer. Systematic evaluation of error rates and causes in short samples in next-generation sequencing. Scientific reports, 8(1):1–14, 2018.

[33] A. Pressman, J. E. Moretti, G. W. Campbell, U. F. Müller, and I. A. Chen. Analysis of in vitro evolution reveals the underlying distribution of catalytic activity among random sequences. Nucleic Acids Research, 45(14):8167–8179, jun 2017.

[34] A. D. Pressman, Z. Liu, E. Janzen, C. Blanco, U. F. Müller, G. F. Joyce, R. Pascal, and I. A. Chen. Mapping a systematic ribozyme fitness landscape reveals a frustrated evolutionary network for self-aminoacylating RNA. Journal of the American Chemical Society, 141(15):6213–6223, mar 2019.

[35] D. Proske, M. Blank, R. Buhmann, and A. Resch. Aptamers—basic research, drug development, and clinical applications. Applied microbiology and biotechnology, 69(4):367–374, 2005.

[36] C. Renzl, A. Kakoti, and G. Mayer. Aptamer-mediated reversible transactivation of gene expression by light. Angewandte Chemie, 132(50):22600–22604, 2020.

[37] M. Rosenthal, F. Pfeiffer, and G. Mayer. A receptor-guided design strategy for ligand identification. Angewandte Chemie International Edition, 58(31):10752–10755, 2019.

[38] C. Roussel, S. Cocco, and R. Monasson. Barriers and dynamical paths in alternating gibbs sampling of restricted boltzmann machines. Physical Review E, 104(3):034109, sep 2021.

[39] W. P. Russ, M. Figliuzzi, C. Stocker, P. Barrat-Charlaix, M. Socolich, P. Kast, D. Hilvert, R. Monasson, S. Cocco, M. Weigt, et al. An evolution-based model for designing chorismate mutase enzymes. Science, 369(6502):440–445, 2020.

[40] A. Schmitz, A. Weber, M. Bayin, S. Breuers, V. Fieberg, M. Famulok, and G. Mayer. A sars-cov-2 spike binding dna aptamer that inhibits pseudovirus infection by an rbd-independent mechanism. Angewandte Chemie International Edition, 60(18):10279–10285, 2021.

[41] A. Schüller, D. Matzner, C. E. Lünse, V. Wittmann, C. Schumacher, S. Unsleber, H. Brötz-Oesterhelt, C. Mayer, G. Bierbaum, and G. Mayer. Activation of the glms ribozyme confers bacterial growth inhibition. Chembiochem, 18(5):435–440, 2017.

[42] L. Sesta, G. Uguzzoni, J. Fernandez-de Cossio-Diaz, and A. Pagnani. Amala: Analysis of directed evolution experiments via annealed mutational approximated landscape. International journal of molecular sciences, 22(20):10908, 2021.

[43] M. Sola, A. P. Menon, B. Moreno, D. Meraviglia-Crivelli, M. M. Soldevilla, F. Cartón-García, and F. Pastor. Aptamers against live targets: is in vivo selex finally coming to the edge? Molecular Therapy-Nucleic Acids, 21:192–204, 2020.

[44] J. Song, Y. Zheng, M. Huang, L. Wu, W. Wang, Z. Zhu, Y. Song, and C. Yang. A sequential multidimensional analysis algorithm for aptamer identification based on structure analysis and machine learning. Analytical chemistry, 92(4):3307–3314, 2019.

[45] T. Tieleman. Training restricted boltzmann machines using approximations to the likelihood gradient. In Proceedings of the 25th International Conference on Machine Learning, ICML ’08, page 1064–1071, New York, NY, USA, 2008. Association for Computing Machinery.

[46] R. J. Townshend, S. Eismann, A. M. Watkins, R. Rangan, M. Karelina, R. Das, and R. O. Dror. Geometric deep learning of rna structure. Science, 373(6558):1047–1051, 2021.

[47] J. Tubiana, S. Cocco, and R. Monasson. Learning protein constitutive motifs from sequence data. eLife, 8:e39397, mar 2019.

[48] J. Tubiana and R. Monasson. Emergence of compositional representations in restricted boltzmann machines. Phys. Rev. Lett., 118:138301, Mar 2017.

[49] C. Tuerk and L. Gold. Systematic evolution of ligands by exponential enrichment: Rna ligands to bacteriophage t4 dna polymerase. Science, 249(4968):505–510, 1990.

[50] A. Wagner. Robustness and evolvability in living systems. Princeton university press, 2013.

[51] J. Zhou and J. J. Rossi. Cell-type-specific, aptamer-functionalized agents for targeted disease therapy. Molecular Therapy-Nucleic Acids, 3:e169, 2014.

[52] Q. Zhou, N. Kunder, J. A. De la Paz, A. E. Lasley, V. D. Bhat, F. Morcos, and Z. T. Campbell. Global pairwise rna interaction landscapes reveal core features of protein recognition. Nature communications, 9(1):1–10, 2018.

[53] Q. Zhou, X. Sun, X. Xia, Z. Fan, Z. Luo, S. Zhao, E. Shakhnovich, and H. Liang. Exploring the mutational robustness of nucleic acids by searching genotype neighborhoods in sequence space. The Journal of Physical Chemistry Letters, 8(2):407–414, jan 2017.

[54] Q. Zhou, X. Xia, Z. Luo, H. Liang, and E. Shakhnovich. Searching the sequence space for potent aptamers using SELEX in silico. Journal of Chemical Theory and Computation, 11(12):5939–5946, nov 2015.

[55] Y. Zhou, X. Qi, Y. Liu, F. Zhang, and H. Yan. Dna-nanoscaffold-assisted selection of femtomolar bivalent human alpha-thrombin aptamers with potent anticoagulant activity. ChemBioChem, 20(19):2494–2503, 2019.

[56] J. Zrimec, F. Buric, M. Kokina, V. Garcia, and A. Zelezniak. Learning the regulatory code of gene expression. Frontiers in Molecular Biosciences, 8, 2021.

