## Supplementary Information for "Generative and interpretable machine learning for aptamer design and analysis of in vitro sequence selection"

#### S1 Experiments

##### S1.1 Gel Electrophoresis

The formation of the Thrombin/DNA complex is sensitive to temperature as well as the presence of $K^+$. Therefore, each 5% native gel incorporated $K^+$ into its matrix and was run at 15C for 90 minutes at 200V. Running buffer was 10mM $K^+$ 7mM $Mg^{2+}$ 1x TAE at 8 pH. Each gel was stained with SYBR gold prior to imaging at 300nm. If using a fluorophore labeled strand, the gel was was not stained prior to imaging.

##### S1.2 DNA Sample Preparation

Each RBM-generated 20nt sequence was supplemented with two complementary 18nt regions to form a stem loop structure (56nt total). The RBM-generated stem loops were designed and their secondary structure predicted (see Figure S1) using NUPACK's webserver [8]. NUPACK results showed no other complex formation except for the desired stem loop. The sequences were ordered, HPLC purified from IDT and re-suspended in 10mM $K^+$ 7mM $Mg^{2+}$ 1x TAE. The stem loops were annealed for 12hrs to ensure proper secondary structure formation, and their concentrations standardized to 500nM by measure of the 260nm absorbance using a Nanodrop Spectrometer. All DCA-generated sequences were originally designed to form the nanotile from the SELEX experiment which generated our dataset [10]. Using each loop individually resulted in a 15nt stem loop with non-pairing regions. These sequences were ordered in a plate from IDT with their standard desalting. Each DCA-generated sequence was purified by using a 5% or 6% denaturing gel (depending on the sequence size) in 1x TBE buffer, cutting the resulting band and precipitating the DNA out with ethanol. The stem loops were annealed for 12hrs to ensure proper secondary structure formation, and their concentrations standardized to 500nM by measure of the 260nm absorbance using a Nanodrop Spectrometer. All sequences used throughout the main text are shown in Table S1 except for any 5' 6FAM modifications which are marked in any figures in which they are used.

##### S1.3 Control Sequence Verification

To confirm the binding band and establish the interaction between the stem loop sequences and thrombin, the control strands ThA and ThD were exposed to varying concentrations of thrombin shown in Figure S2b,c. For both ThA and ThD, the almost complete uptake of the stem-loop from the starting position (Figure S2) to the stem-loop / complex band at a ratio of 1:1.08 indicates the stem loop / complex band interaction is made up of a single stem loop binding to a single molecule of thrombin. Further, the combination of the two stem loops binding to thrombin at the same concentrations (Fig S2d) confirmed the cooperative binding seen in previous experiments as well as indicated a downshift of the stem-loop / protein band upon 2 stem-loops binding to thrombin.

##### S1.4 Competition Assays

Competition assays were performed by mixing equimolar amounts (2.5um) of a fluorophore labeled DNA strand and non-labeled DNA strand that bind to the same Thrombin exosite. The reverse is simultaneously tested, where the fluorophore labeled version of the non-labeled strand is substituted

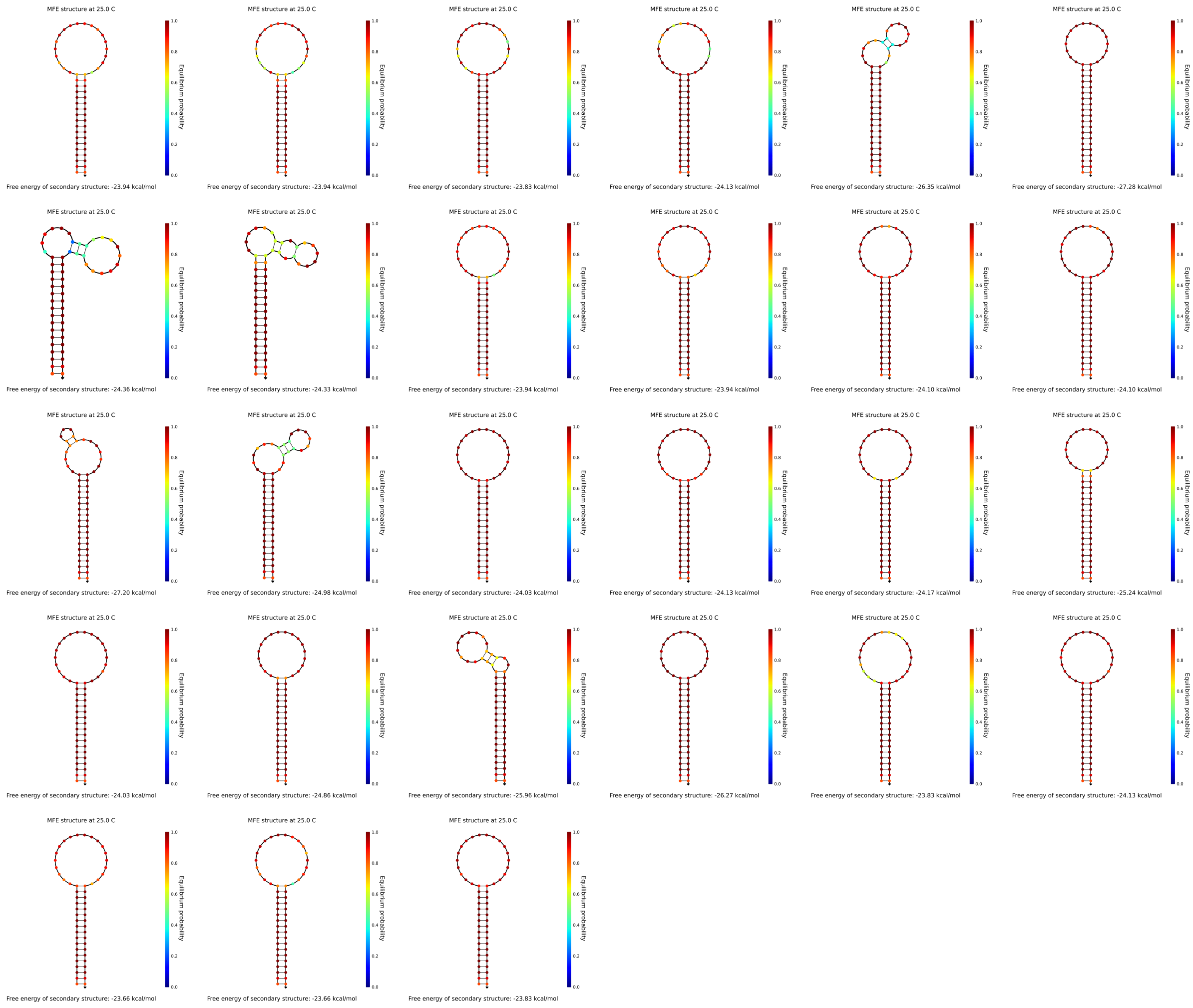

Figure S1: Nupack Predictions of the minimum free energy structure (MFE) of each DNA stem-loop at 25C. Figures start from r1 in the top left corner to r27 in the bottom right corner.

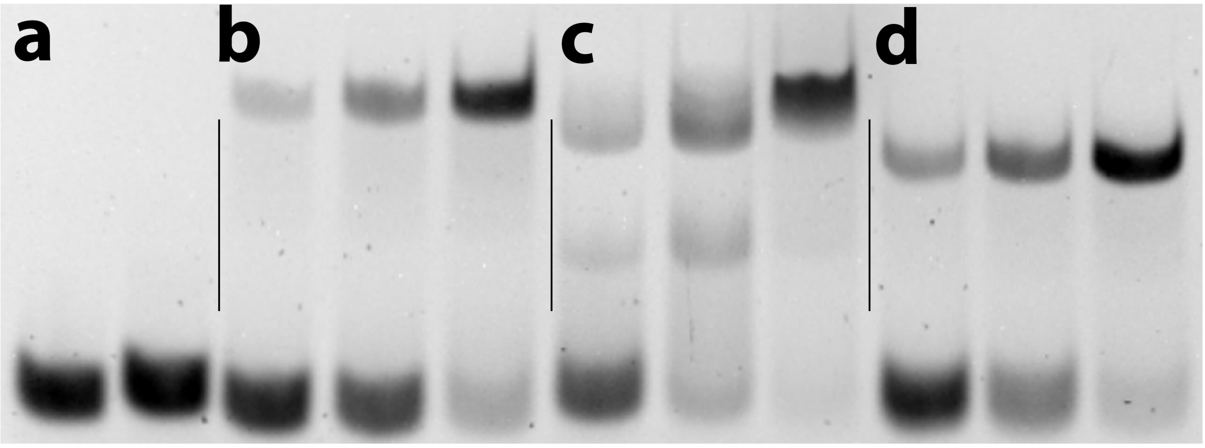


Figure S2: Panel a: lane 1 contains 5' 6FAM labeled control sequence ThA, lane 2 contains 5' 6FAM labeled control sequence ThD.
Panel b: ThA mixed in varying ratios with Thrombin (1:0.32, 1:0.64, 1:1.08).
Panel c: ThD mixed in varying ratios with Thrombin (1:0.32, 1:0.64, 1:1.08).
Panel d: ThA + ThD mixed in varying ratios with Thrombin (1:1:0.32, 1:1:0.64, 1:1:1.08).

| Sample Name | Full Reported Sequence |
|---|---|
| r1 | CTCGAGAGTTGCAGAAGTAGTGATGATGTGTGGTAGGCACTTCTGCAACTCTCGAG |
| r2 | CTCGAGAGTTGCAGAAGTAGTGTAGGTGTGGATGATGCACTTCTGCAACTCTCGAG |
| r3 | CTCGAGAGTTGCAGAAGTTAGGTTTTGGGTAGCGTGGTACTTCTGCAACTCTCGAG |
| r4 | CTCGAGAGTTGCAGAAGTAGGGATGATGTGTGGCAGGAACTTCTGCAACTCTCGAG |
| r5 | CTCGAGAGTTGCAGAAGTCTAGGACGGGTAGGGCGGTGACTTCTGCAACTCTCGAG |
| r6 | CTCGAGAGTTGCAGAAGTAGGGATGTGTGTGGTAGGCTACTTCTGCAACTCTCGAG |
| r7 | CTCGAGAGTTGCAGAAGTAGGGATGCTGCGTGGTAGGCACTTCTGCAACTCTCGAG |
| r8 | CTCGAGAGTTGCAGAAGTGAGGGTTGGTGTGGTTGGCAACTTCTGCAACTCTCGAG |
| r9 | CTCGAGAGTTGCAGAAGTAGGGTTGGTGTGTGGTTGGCACTTCTGCAACTCTCGAG |
| r10 | CTCGAGAGTTGCAGAAGTATGGTTGGTTTATGGTTGGCACTTCTGCAACTCTCGAG |
| r11 | CTCGAGAGTTGCAGAAGTGAAGGGTGGTCAGGGTGGGAACTTCTGCAACTCTCGAG |
| r12 | CTCGAGAGTTGCAGAAGTGGAGGGTGGGTCGGGTGGGAACTTCTGCAACTCTCGAG |
| r13 | CTCGAGAGTTGCAGAAGTGGGGTTGGTACAGGGTTGGCACTTCTGCAACTCTCGAG |
| r14 | CTCGAGAGTTGCAGAAGTAGATGGGCAGGTTGGTGCGGACTTCTGCAACTCTCGAG |
| r15 | CTCGAGAGTTGCAGAAGTAGATGGGTGGGTAGGGTGGGACTTCTGCAACTCTCGAG |
| r16 | CTCGAGAGTTGCAGAAGTATAGGGTGGGTGGGTGGGTAACTTCTGCAACTCTCGAG |
| r17 | CTCGAGAGTTGCAGAAGTTGGTGGTTGGGTTGGGTTGGACTTCTGCAACTCTCGAG |
| r18 | CTCGAGAGTTGCAGAAGTTGGGATGGGATTGGTAGGCGACTTCTGCAACTCTCGAG |
| r19 | CTCGAGAGTTGCAGAAGTAGGGTTGGTTATGTGGTTGGACTTCTGCAACTCTCGAG |
| r20 | CTCGAGAGTTGCAGAAGTATTGGTTGGGTAGGGTGGTTACTTCTGCAACTCTCGAG |
| r21 | CTCGAGAGTTGCAGAAGTAAACGGTTGGTGAGGTTGGTACTTCTGCAACTCTCGAG |
| r22 | CTCGAGAGTTGCAGAAGTCGGGGTGGTGTGGGTGGGAGACTTCTGCAACTCTCGAG |
| r23 | CTCGAGAGTTGCAGAAGTTATTGGTTGGATAGGTTGGTACTTCTGCAACTCTCGAG |
| r24 | CTCGAGAGTTGCAGAAGTAGGGTTGGGTGGTTGGATGAACTTCTGCAACTCTCGAG |
| r25 | CTCGAGAGTTGCAGAAGTCGGGTTGGGGGGTTGGATTCACTTCTGCAACTCTCGAG |
| r26 | CTCGAGAGTTGCAGAAGTCGGTTGGGGGGGTTGGATACACTTCTGCAACTCTCGAG |
| r27 | CTCGAGAGTTGCAGAAGTTGTGGGTTGGTGAGGTAGGTACTTCTGCAACTCTCGAG |
| ThA | CTCGAGAGTTGCAGAAGTAGGGATGATGTGTGGTAGGCACTTCTGCAACTCTCGAG |
| ThD | CTCGAGAGTTGCAGAAGTGTAGGATGGGTAGGGTGGTCACTTCTGCAACTCTCGAG |
| p1 | CTCGAGAGTTGCAGAAGTAGGGATGATGTGTGGTTGGCACTTCTGCAACTCTCGAG |
| p2 | CTCGAGAGTTGCAGAAGTAGGGATGGTGTGTGGTAGGCACTTCTGCAACTCTCGAG |
| p3 | CTCGAGAGTTGCAGAAGTAGGGTTGATGTGTGGTAGGCACTTCTGCAACTCTCGAG |
| p4 | CTCGAGAGTTGCAGAAGTAGGGATGGTGTGTGGTTGGCACTTCTGCAACTCTCGAG |
| p5 | CTCGAGAGTTGCAGAAGTAGGGTTGATGTGTGGTTGGCACTTCTGCAACTCTCGAG |
| p6 | CTCGAGAGTTGCAGAAGTAGGGTTGGTGTGTGGTAGGCACTTCTGCAACTCTCGAG |
| d1 | TCAGGCTCTCGAGAGTTGCAGAAGTAGGGTAGGTGTGGGGTATGCACTTCTGCCTGCATCGAGACA |
| d2 | TCAGGCTCTCGAGAGTTGCAGAAGTAGGGTAGATGTGTAGGATGCACTTCTGCCTGCATCGAGACA |
| d3 | TCAGGCTCTCGAGAGTTGCAGAAGTAGGGATGATGGTTGGTAGGCACTTCTGCCTGCATCGAGACA |
| d4 | TCAGGCTCTCGAGAGTTGCAGAAGTAGGGATGATGTGGATTAGGCACTTCTGCCTGCATCGAGACA |
| d5 | TCAGGCTCTCGAGAGTTGCAGAAGTAGGGTGGGAGCGGGGGACGCACTTCTGCCTGCATCGAGACA |
| d6 | TCAGGCTCTCGAGAGTTGCAGAAGTCGGGTAGGTGTGGATTATGCACTTCTGCCTGCATCGAGACA |
| d7 | TCAGGCTCTCGAGAGTTGCAGAAGTGTAGGACGGGTAGGGCGGTCACTTCTGCCTGCATCGAGACA |
| d8 | TCAGGCTCTCGAGAGTTGCAGAAGTGGGGGTTGGGCGGGATGGGCACTTCTGCCTGCATCGAGACA |
| d9 | TCAGGCTCTCGAGAGTTGCAGAAGTGCGGGTTGGGCAGGATCAGCACTTCTGCCTGCATCGAGACA |
| d10 | TCAGGCTCTCGAGAGTTGCAG AAGTAGGGATGATGTGTGGTAGGCACTTCTGCCTGCATCGAGACA |
| d11 | /5PHOS/CCAGTTTTTCTGGTGAGCTAGTGCAGACATGATCGTAGGATGGGTGGGGTGGGAGATCATGTAACTCCTAGCTGCCTGA |
| d12 | /5PHOS/CCAGTTTTTCTGGTGAGCTAGTGCAGACATGATCGTAGGATGGGTAGGGTGGTAGATCATGTAACTCCTAGCTGCCTGA |
| d13 | /5PHOS/CCAGTTTTTCTGGTGAGCTAGTGCAGACATGATCCTAGGTTGGGTAGGGTGGTGGATCATGTAACTCCTAGCTGCCTGA |
| d14 | /5PHOS/CCAGTTTTTCTGGTGAGCTAGTGCAGACATGATCCTAGCATGGGTAGGGTGGTGGATCATGTAACTCCTAGCTGCCTGA |
| d15 | /5PHOS/CCAGTTTTTCTGGTGAGCTAGTGCAGACATGATCGTAGCATGGGTAGGGTGGTCGATCATGTAACTCCTAGCTGCCTGA |
| d16 | /5PHOS/CCAGTTTTTCTGGTGAGCTAGTGCAGACATGATCTTGGGTGGTGTAGGTTGGCGGATCATGTAACTCCTAGCTGCCTGA |
| d17 | /5PHOS/CCAGTTTTTCTGGTGAGCTAGTGCAGACATGATCTTGGGTGGTGCAGGTTCGCGGATCATGTAACTCCTAGCTGCCTGA |
| d18 | /5PHOS/CCAGTTTTTCTGGTGAGCTAGTGCAGACATGATCCTAGGATGGGTAGGGTGGTGGATCATGTAACTCCTAGCTGCCTGA |

Table S1: The full sequences from all experiments carried out in this work, with their loop region underlined for easy identification. r1-27 correspond to sequences generated from sampling our RBM. All sequences with p labels (p1-p6) are along the mutation pathway from sequence ThA to r9. Sequences d1-d9 and d11-d17 are were generated from sampling from the DCA parameters. Sequences d10, d18, ThA, and ThD were used as controls throughout.

with a fluorophore labeled version and the previously non-labeled strand is substituted for a fluorophore labeled version. In both, Thrombin is added in a 1:2 ratio (2.5um) and allowed to mix at 25°C for 30 min. Comparing the results of the assays yields a conclusive ranking of the relative binding affinity of the two sequences. Competition assays using 5' 6FAM modified sequences are depicted in Fig. S3.

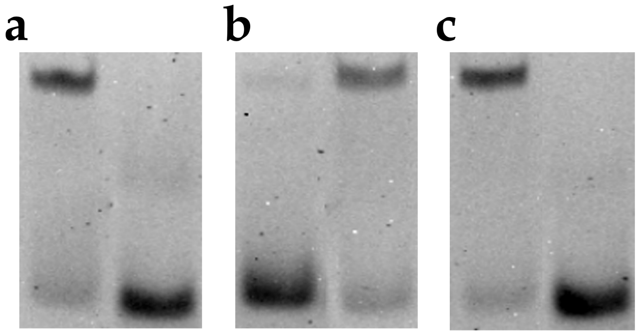


Figure S3: Competition assay of r8F vs r14 and r14F vs r8 (panel a). r8F vs r19 and r19F vs r8 (panel b), and r19F vs r14 and r14F vs r19 (panel c). The F suffix indicates the strand is fluorohore labeled with a 5' 6FAM modification.

Additionally, one-sided competition assays for all exosite-I binders and all sequences between r9 and ThA were tested against fluorophore-labeled versions of the best binding aptamers from the previous study (ThA and ThD) to assess whether any novel binder performed better. From Fig. S4b,c we see that no exosite-II binding aptamer was found which bound better than ThA and no exosite-I binding aptamer was found which bound better than ThD. Additionally, the tested exosite-I binders Fig. S4(a) are worse binders than r8 and r19 but better binders than r14.

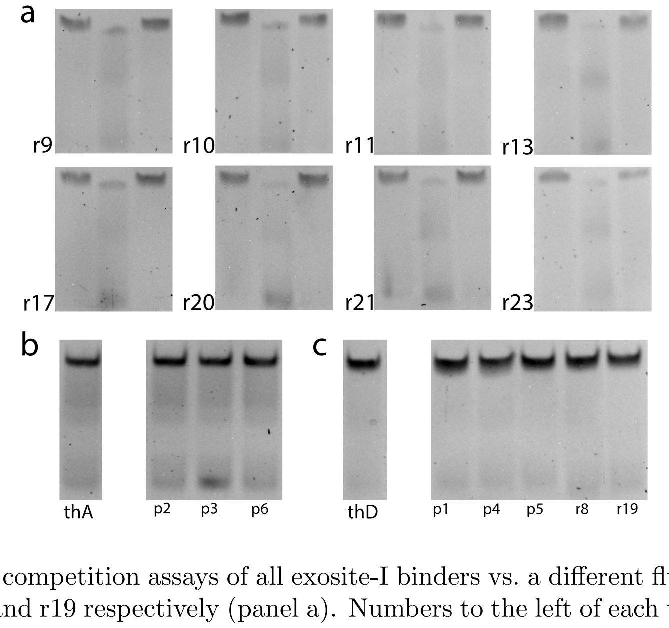


Figure S4: One sided competition assays of all exosite-I binders vs. a different fluorophore labeled strand in each well, r8, r14, and r19 respectively (panel a). Numbers to the left of each trial indicate the identity of the non-labeled strand. Additionally exosite-II binders were tested against fluorophore labeled ThA with negative control labeled ThA (panel b) and select exosite-I binders were tested against ThD with negative control labeled ThD (panel c).

#### S1.5 Thrombin Sample Preparation

We used 1 mg Human $\alpha$-thrombin manufactured by Haematologic Technologies Inc. and purchased from Fisher Scientific Co. Concentrations were assessed by 280nm absorbance using a Nanodrop Spectrometer. The stock was stored at -20C. Sample concentrations were made at 500nm and 250nm in 1x PBS 10mM $K^+$ 7mM $Mg^{2+}$. Each sample was made fresh prior to being used in an assay.

#### S2 Inference of sequencing error probability

We describe here our method for inferring the single-site sequencing error probability. The analysis here discussed is based on sequences from the left loop, collected at the last selection round (round 8). Repeating the analysis on the right loop provides analogous results.
Given a sequence $\sigma$ with high copy number $n_\sigma \gg 1$, the method uses as signal the number $\mu_\sigma$ of sequences that are at Hamming distance 1 from $\sigma$ and are never observed in the dataset. This number depends on the error rate, since a higher error rate is expected to cause more of these sequences to be detected. Since we consider only the left loop, sequences have a length $L = 20$ nt.
In Fig. S5A we provide a representation of the sequence space around $\sigma$ = `AGGGATGATGTGTGGTAGGC`, which is the sequences with highest copy number $n_\sigma = 8034$ in our dataset. The dots in a circle around $\sigma$ represent the $3 \times L = 60$ sequences that belong to the neighborhood of the main sequence $\mathcal{N}(\sigma)$, with color encoding their copy number. Some of these sequences are present $> 100$ times, and are unlikely to be an artifact of sequencing error. Other are present 1-2 times and can potentially be generated by sequencing errors. Finally, a number $\mu_\sigma = 12$ of sequences are absent in the sample (red crosses). These are mostly related to mutations removing one Guanine from the sequence, which might be related to a loss of fitness. While it is not possible to know with certainty whether one of the present neighbouring sequences with low copy-number was originated by sequencing error, the fact that some of these sequences are absent implies that $\sigma$ was never mis-read into these sequences. This information will be used in our inference. We start by selecting a number of sequences with high copy number. In fig. S5B we plot the number of sequences that have copy-number higher than a given threshold, as a function of the threshold. For our analysis we select as "peaks" all sequences with $n_\sigma > 1000$ (21 such sequences in the dataset). In Fig. S5C we report the Hamming distance matrix for the selected sequences. As can be expected peaks tend to cluster together, with most of the peaks having at least one other peak in their neighbourhood. This can potentially increase the bias in our upper bound for the sequencing error probability. We will later introduce a correction to reduce this bias.

As a next step we define a probability for $\mu_\sigma$ as a function of the reading error probability. We call $\epsilon$ the probability of mis-reading a single nucleotide in the sequence. We consider this probability to be uniform along the sequence and on the real/read nucleotides, so that the probability of obtaining as outcome of sequencing $\sigma'$, one of the single-site mutations $\mathcal{N}(\sigma)$ of $\sigma$, when in reality reading $\sigma$ is:

$$P(\sigma'|\sigma) = p(\epsilon) = \frac{\epsilon}{3}(1-\epsilon)^{L-1}. \tag{S1}$$

The real copy-number $\widetilde{n}_\sigma$ of $\sigma$ in the sample might be slightly different from the observed copy number $n_\sigma$, due to sequencing error. If we call $P(\sigma|\sigma) = (1-\epsilon)^L$ the probability of correctly reading $\sigma$, then for a small enough error, we can approximate

$$n_\sigma \simeq \widetilde{n}_\sigma P(\sigma|\sigma) + p(\epsilon) \sum_{\sigma' \in \mathcal{N}(\sigma)} \widetilde{n}_{\sigma'} \simeq \widetilde{n}_\sigma P(\sigma|\sigma). \tag{S2}$$

For any given sequence $\sigma' \in \mathcal{N}(\sigma)$, the probability of never mis-reading $\sigma'$ when in reality sequencing $\sigma$ is given by:

$$P(n_{\sigma'} = 0) = (1 - p(\epsilon))^{\widetilde{n}_\sigma} = q(\epsilon, n_\sigma). \tag{S3}$$

Finally, the probability that in the neighbourhood of $\sigma$ a number $\mu_\sigma$ of sequences are never observed, provided that in reality they were never present, is:

$$P(\mu_\sigma|n_\sigma, \epsilon) = \mathrm{Binom}\big[|\mathcal{N}(\sigma)|, q(\epsilon, n_\sigma)\big](\mu_\sigma) = \binom{|\mathcal{N}(\sigma)|}{\mu_\sigma} \big(q(\epsilon, n_\sigma)\big)^{\mu_\sigma} \big(1 - q(\epsilon, n_\sigma)\big)^{|\mathcal{N}(\sigma)| - \mu_\sigma}, \tag{S4}$$

where $|\mathcal{N}(\sigma)| = 60$ is the size of the neighbourhood of $\sigma$. When writing this equation we are making a number of simplifications. On one hand we are considering that all sequences in $\mathcal{N}(\sigma)$ were originally absent in the sample. Moreover we are neglecting the probability that reads of these sequences might be generated from the sequencing of other sequences different from $\sigma$ (e.g. other peaks). All of these effects will bias our estimate, but the bias is always in the same direction, leading us to overestimate $\epsilon$. For this reason the result of the inference represents a reliable upper bound.
To reduce the bias we can remove from the total number of trials in the binomial the number of sequences that we are confident to be really present in the original sample. As a simple correction, we substitute the term $|\mathcal{N}(\sigma)| = 60$ in eq. (S4) with $|\{\sigma' \in \mathcal{N}(\sigma) \text{ s.t. } n(\sigma') \leq 10\}|$, i.e. the number of sequences in the neighbourhood with no more than 10 counts. That is to say we consider all sequences with more than

10 counts to be really present in the original sample. We perform the inference both with and without this correction (cf. fig. S5D).
At this point we can write the total log-likelihood of our data as a function of the error probability $\epsilon$ as:

$$\log \mathcal{L}(\text{data}|\epsilon) = \sum_{\sigma \in \text{peaks}} \log P(\mu_\sigma | n_\sigma, \epsilon) \ \propto \ \log \mathcal{L}(\epsilon|\text{data}), \tag{S5}$$

where the inversion was operated using Bayes theorem with an uniform prior for $\epsilon$. In fig. S5D we display the behavior of the log-likelihood as a function of $\epsilon$ for the two cases, with and without correction. Numerical maximization of these functions yields values of $\epsilon^* \sim 10^{-3}$ as an upper bound for the error probability. To obtain a confidence interval on these bounds one can perform a Gaussian fit on the likelihood (i.e. a quadratic fit of the log-likelihood) around its maximum, and use the variance of the inferred Gaussian to obtain a confidence interval for the inferred value. In this case we obtain a standard deviation of the order of $4 \times 10^{-5}$.
In conclusion, we are confident that the single-site sequencing error probability in our dataset is smaller than $10^{-3}$.

#### Estimation of number of sequencing error artifacts in the dataset

We can make use of the previously derived upper bound for $\epsilon$ to provide an upper bound for the number of unique sequences in our dataset that could be generated by sequencing error.
Since we expect double errors to be sufficiently rare in our dataset (for $\epsilon^* \sim 10^{-3}$ the probability of having more than 1 error is $\sim 2 \times 10^{-4}$), we can consider that in order to be an error, all the reads of a sequence $\sigma$ in our dataset must be generated by sequences in its neighbourhood $\mathcal{N}(\sigma)$, with the probability of mis-reading being equal to $p(\epsilon)$ (see Eq. (S1)). For each sequence we define:

$$N_\sigma = \sum_{\sigma' \in \mathcal{N}(\sigma)} n_{\sigma'}. \tag{S6}$$

This is the total number of sequences in the neighborhood of $\sigma$. Because of sequencing error the real number might be slightly higher, and as done for $n_\sigma$ one can introduce the correction $\widetilde{N}_\sigma = N_\sigma/(1-\epsilon)^L$. We can take as an upper bound for the probability of $\sigma$ to be an artifact of sequencing error, the probability that by reading $\widetilde{N}_\sigma$ sequences in the neighbourhood of $\sigma$, we read $\sigma$ a number of time equal or greater than the observed copy-number $n_\sigma$:

$$P(n_{\text{err}} \geq n_\sigma) = \pi(\sigma, \epsilon) = \sum_{k=n_\sigma}^{\infty} \text{Binom}[N_\sigma, p(\epsilon)](k) \tag{S7}$$

We numerically evaluate this probability for every sequence $\sigma$. The value of $N_\sigma$ is efficiently computed by generating all possible single mutations $\sigma' \in \mathcal{N}(\sigma)$, and quickly recovering their copy-number using a hash table.
In Fig. S6 we report the distribution of $\pi(\sigma, \epsilon^* = 10^3)$ for all of the sequences in our dataset. For the great majority of the sequences this probability is very low. From the procedure we employ it follows that sequences with the highest probability of being errors are ones that have very low $n_\sigma$ and with a highly populated neighbourhood (high $N_\sigma$). By treating the reality of each unique sequence as a Bernoulli random variable, the mean and variance for the total number of unique sequences that we expect to be an artifact of sequencing error can be expressed as:

$$E[N_{err}] = \sum_\sigma \pi(\sigma, \epsilon^*) \quad Var[N_{err}] = \sum_\sigma \pi(\sigma, \epsilon^*)(1 - \pi(\sigma, \epsilon^*)) \tag{S8}$$

This gives an estimate $N_{err} \sim 941 \pm 28$. Since our dataset is composed of roughly $2 \times 10^5$ unique sequences this upper bound represents only 0.5% of the total dataset, and it is not expect to meaningfully impact the training of our models.

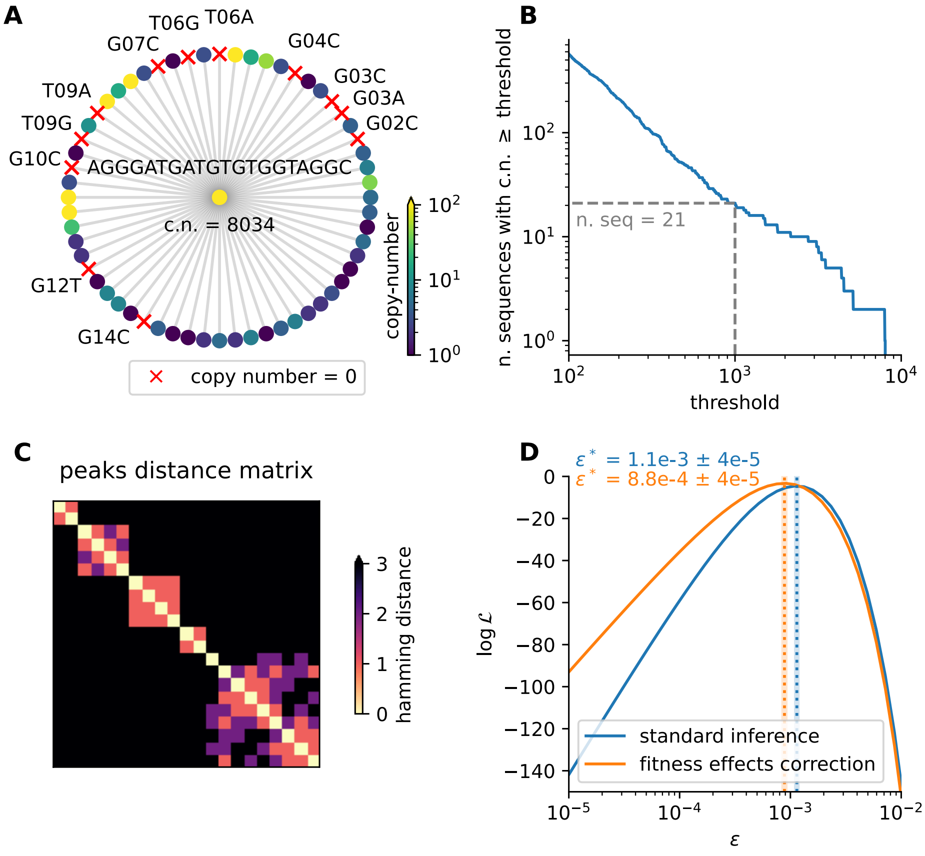


Figure S5: inference of an upper bound for the sequencing error probability in our sample, using sequences from the left loop in the 8th round.

Panel A: example of sequence space around the most abundant sequence in the dataset. The main sequence is represented as a dot in the center, and the full DNA sequence and copy number (c.n.) are reported. Dots around it represent sequences at hamming distance 1, with color encoding their copy number. Sequences that were never detected in the sample are indicated with red crosses. For these sequences we report the difference from the main sequence as a triplet (original nucleotide, position, substituted nucleotide). Notice how some of the neighboring sequences have high copy number, indicating probable fitness effects. Most of the non-detected sequences are associated with removal of a guanine, which might decrease binding affinity.

Panel B: number of sequences with copy-number greater than a given threshold. For our analysis we select only sequences with c.n. $\geq 1000$ (21 such sequences in the dataset). These sequences are referred to as "peaks" in the analysis.

Panel C: relative Hamming distance between peak sequences. High-copy-number sequences tend to cluster together. This can cause a less precise estimation of the inferred sequencing error upper bound, since the neighbourhood of a peak can be populated by other high-fitness sequences. To correct for this we introduce a correction that removes sequences with c.n. $> 10$ from the expression of the likelihood.

Panel D: log-likelihood of the single-site sequencing error probability $\epsilon$. The inference was performed in two ways: either using the standard approach (blue) or introducing the correction for fitness effects (orange). In each case we mark the inferred value $\epsilon^*$ with vertical dotted lines. The thin shaded area represent the confidence interval, that was derived through a Gaussian fit of the log-likelihood in proximity of its maximum.

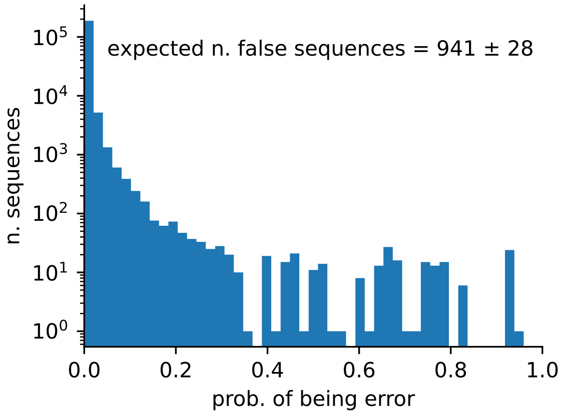


Figure S6: Distribution of inferred single-sequence error probabilities. For each sequence in the considered dataset (round 8, left loop) we infer the probability of being an artifact of sequencing error, using the approach described in Methods S2. In the inference the single-site error probability was set equal to the upper bound $\epsilon^* = 10^{-3}$. The vast majority of sequence have a zero or low probability of being sequencing error artifacts. From this distribution one can evaluate the mean and standard deviation of the total number of artifacts. This gives an upper bound of $N_{err} = 941 \pm 28$, which corresponds to a 0.5% of the total number of unique sequences in the dataset considered.

#### S3 Details of RBMs' training

We trained several RBMs which have been used for the analysis presented in this manuscript. For the training of each RBM, we used 90% of the dataset as training set and 10% of the dataset as validation set to check that no overfitting is observed. For RBMs trained using information on counts, the training dataset is obtained by sampling from the dataset of unique sequences many sequences (below the exact number for each RBM is given), with a probability proportional to each sequence's count (this gives the same results as long as the size of the re-sampled dataset is large enough, and allows to avoid having too large dataset which considerably slow down the RBM training). The training set was divided in mini-batches and for each epoch each mini-batch was used to perform an update of the parameters, using the persistent contrastive divergence algorithm with few (below the exact numbers are given) number of Monte-Carlo steps for each update of the paramters. In all cases the training stopped after 20000 updates of the RBM parameters. Finally, we used a $L_1^2$ regularization of the form given in Eq. (6) to increase sparsity in the weights, which in turn improves the interpretability of the contribution of each hidden unit to a sequence's log-likelihood. Below we give the regularization parameter $\lambda$ used for each RBM trained (see Eq. (6)).

For the full range of explored hyperparameters (size of mini-batches, number of Monte-Carlo steps, regularization strength), we never saw any sign of relevant overfitting, and we motivated this with the very large datasets that are available for training the models.

The code used to train the RBMs can be obtained from https://github.com/jertubiana/PGM.
We now give more details about the training of each RBM model used in this manuscript. To distinguish RBMs trained with sequences observed in different rounds, we will append the round number to the model name. In particular, we used the following RBMs in this manuscript:

- RBM-DC6 (Fig. 2, Suppl. Figs. S19, S27), trained on the double aptamers (40 nucleotides) obtained from the SELEX 6th round. The training set is built by re-sampling 736436 sequences from the dataset of unique double-loop sequences observed in round 6, using their number of counts as weight for the sampling. The parameters are: 40 visible units, 90 hidden units, $\lambda = 0.01$, 10 Monte-Carlo steps for each update of the parameters, mini-batches of size 500.

- RBM-DC8 (Figs. 3, 4, Suppl. Figs. S18, S20), trained on the double aptamers (40 nucleotides) obtained from the SELEX 8th round. The training set is built by re-sampling 719413 sequences from the dataset of unique double-loop sequences observed in round 8, using their number of counts as weight for the sampling. The parameters are: 40 visible units, 90 hidden units, $\lambda = 0.01$, 10 Monte-Carlo steps for each update of the parameters, mini-batches of size 500.

- RBM-SC8 (Figs. 3, 4, 8, Table 1, Suppl. Figs. S16, S17, S23, S25, S26), trained on the single aptamers (20 nucleotides) obtained from the SELEX 8th round. The training set is built by re-sampling 725431 sequences from the dataset of unique single-loop left or right sequences observed in round 8, using their number of counts as weight for the sampling. The parameters are: 20 visible units, 80 hidden units, $\lambda = 0.01$, 2 monte-carlo steps for each update of the parameters, mini-batches of size 1000.

- RBM-SU8 (Figs. 5, 8, Table 1, Suppl. Figs. S14, S20, S21, S23, S25), trained on the single aptamers (20 nucleotides) obtained from the SELEX 8-th round, merging sequences from the left and right loops (unique single-loop sequences: 382094; with counts: 1450862). Multiple copies of the same aptamer are neglected. The parameters are: 20 visible units, 70 hidden units, $\lambda = 0.01$, 4 Monte-Carlo steps for each update of the parameters, mini-batches of size 500.

- RBM-SC5 (Suppl. Figs. S24), trained on the single aptamers (20 nucleotides) obtained from the SELEX 5th round. The training set is built by re-sampling 1375403 sequences from the dataset of unique single-loop left or right sequences observed in round 5, using their number of counts as weight for the sampling. The parameters are: 20 visible units, 70 hidden units, $\lambda = 0.01$, 8 monte-carlo steps for each update of the parameters, mini-batches of size 1500.

- RBM-SC6 (Suppl. Figs. S24), trained on the single aptamers (20 nucleotides) obtained from the SELEX 6th round. The training set is built by re-sampling 598696 sequences from the dataset of unique single-loop left or right sequences observed in round 6, using their number of counts as weight for the sampling. The parameters are: 20 visible units, 80 hidden units, $\lambda = 0.01$, 8 monte-carlo steps for each update of the parameters, mini-batches of size 600.

- RBM-SC7 (Suppl. Figs. S24), trained on the single aptamers (20 nucleotides) obtained from the SELEX 7th round. The training set is built by re-sampling 419934 sequences from the dataset of unique single-loop left or right sequences observed in round 7, using their number of counts as weight for the sampling. The parameters are: 20 visible units, 70 hidden units, $\lambda = 0.01$, 8 monte-carlo steps for each update of the parameters, mini-batches of size 500.
- RBM-SU6 (Suppl. Fig. S27), trained on the single aptamers (20 nucleotides) obtained from the SELEX 6-th round, merging sequences from the left and right loops (unique single-loop sequences: 598696; with counts: 1472872). Multiple copies of the same aptamer are neglected. The parameters are: 20 visible units, 70 hidden units, $\lambda = 0.01$, 4 Monte-Carlo steps for each update of the parameters, mini-batches of size 600.
- RBM-LC8, RBM-RC8 (Suppl. Fig. S16), trained on the single aptamers (20 nucleotides) obtained from the SELEX 8-th round. The training sets of RBM-LC8 (RBM-RC8) is built by re-sampling 177014 (227789) sequences from the dataset of unique left-loop (right-loop) sequences observed in round 8, using their number of counts as weight for the sampling. The parameters of both models are: 20 visible units, 70 hidden units, $\lambda = 0.01$, 4 Monte-Carlo steps for each update of the parameters, mini-batches of size 500.
- RBM-NPU8 (Suppl. Fig. S17), trained on the single aptamers (20 nucleotides) obtained from the SELEX 8-th round, merging sequences from the left and right loops, after excluding parasite sequences. Parasite sequences are obtained here as single-loop sequences with log-likelihood computed by RBM-SU8 lower than -24.8, with the partner loop having log-likelihood computed by RBM-SU8 larger than -24.8 (procedure resulting in 276682 unique single-loop non-parasite sequences). Multiple copies of the same aptamer are neglected. The parameters are: 20 visible units, 70 hidden units, $\lambda = 0.01$, 4 Monte-Carlo steps for each update of the parameters, mini-batches of size 500.
- RBM-NPC8 (Suppl. Fig. S17), trained on the single aptamers (20 nucleotides) obtained from the SELEX 8-th round, merging sequences from the left and right loops, after excluding parasite sequences. Parasite sequences are obtained here as single-loop sequences with log-likelihood computed by RBM-SC8 lower than -26.6, with the partner loop having log-likelihood computed by RBM-SC8 larger than -26.6 (procedure resulting in 274250 unique single-loop non-parasite sequences). The training dataset is built by sampling 246825 non-parasite sequences, using their number of counts as weight for the sampling. The parameters are: 20 visible units, 70 hidden units, $\lambda = 0.01$, 4 Monte-Carlo steps for each update of the parameters, mini-batches of size 500.

All the parameters for the training which are not given here are the default parameters as defined in the code. The trained RBMs are provided in the Github repository (https://github.com/adigioacchino/RBMsForAptamers), together with a jupyter notebook that can be used to re-train them.

As a final remark, we checked that the results obtained here depend very little on the precise values of the hyperparameters used here (see Suppl. Fig. S22). The only notable exception being the usage of counts to weight multiple occurrences of the same aptamer in the dataset. We decided to exclude multiple occurrences from the training to regularize the RBM, as discussed in in details in Suppl. Sec. S26.

### S4 DNN and Traditional Machine Learning

#### S4.1 Dataset preparation

Starting with the raw SELEX data from the 8th round of selection of our previous study, we have both 20nt aptamer sequences in each arm of the DNA scaffold and a copy number, representing the number of times that sequence was observed during sequencing. Any sequence not matching the 40nt length was assumed to have a reading error and excluded from the dataset. Independent counts for each arm of the sequence were generated by counting their occurrence throughout the dataset. Using their individual counts, each 20nt sequence was categorized as either as a "good" (copy number $> 10$) or "bad" (copy number $< 10$) binder. Note that this approach is distinct from our training of the RBMs, where we considered all sequences in the training sample and used counts for weighting the sequences.

Our analysis of the dataset found a subset of bad binder sequences far in sequence space from any other observed sequence in the dataset that were paired with good binder sequences. We concluded that these sequences were most likely carried through the selection process by their good binder and subsequently excluded these sequences from our training set.

Three datasets were generated from the remaining sequences: sequences from the left loop (L), sequences from the right loop (R), and sequences from both loops (B). Each dataset consists of the entire set of good binders from the appropriate loop and 5 randomly sampled bad binders per good binder. Training sets (80% of good binders, $\sim 35k$ in total sequences for L and R, $\sim 70k$ for B) and validation sets (20% of good binders, $\sim 12k$ in total sequences for L and R, $\sim 25k$ for B) were split from our dataset. As further verification of our DNN and traditional ML models we used the experimental results from both the RBM-generated sequences as well as the DCA-model generated sequences to assess our models accuracy. All sequences were one-hot encoded prior to training, validation, or prediction.

As using only sequences from the final round of the SELEX procedure introduces a general bias of all sequences interacting with thrombin, three more datasets were created (GL, GR, and GB) with good binders selected as previously done but bad binders were randomly sampled from a set of random sequences outside the SELEX dataset's sequence space. These datasets had the same amount of sequences as those mentioned previously (L, R, B). We assume that if there is no bias in the initial random library, most of the possible aptamer sequences of length 20 were initially present, and hence a randomly generated sequence which is not encountered in the SELEX dataset is most likely not going to be able to bind to thrombin.

#### S4.2 Model Selection

For the classification task we used 5 different deep learning models: 2 versions of a Variational Auto Encoder [4], 2 versions of a Resnet [3] and a Siamese Network Model [7] outlined in Suppl. Table S2. A schematic description of the DNN model specifications used in this work is provided in Table S2. Additionally we used 3 classic Machine Learning methods: a decision tree, a random forest and a gradient boosted tree classifier to also classify the sequences as binders or non binders.

#### S4.3 DNN Training Specifics

All 5 models were written as pytorch lightning modules and hyperparameter optimization was done using the raytune library. Integration of each pytorch module with raytune enabled simultaneous distributed hyperparameter optimization. All models were trained for either 30 or 50 epochs. No significant performance increase or decrease was observed between models trained for 30 vs 50 epochs.

Hyperparameter optimization was performed using the raytune library. For resnet, we optimized the batch size, learning rate (lr) and dropout (dr) prior to the dense layer and softmax. For variational Auto-Encoders, we optimized the batch size, learning rate, dropout and z_dim (embedding dimension). For the siamese network, we optimized the learning rate, batch size, and distance cutoff (Euclidean distance cutoff, being less means a match while being greater indicates a nonmatch). As a grid search, the AsyncHyperBandScheduler (AHSA) was given 10 trials with the goal to find the model with best accuracy on the validation set. Bayesian Optimization was performed on the same hyperparameters as the ASHA, save the integer valued batch size. Bayesian optimization was given a different directive, to minimize the mean loss (training+validation). Population-based training was only performed on the siamese network with the goal of maximizing the accuracy on the validation set.

| Model | Description |
|---|---|
| Long Resnet | A 152 layer Resnet followed by a dropout layer, a 512 input to 2 output linear layer followed by a softmax layer. Residual networks guarantee performance of subsequent layers in the network by mapping to a residual function F(x) = H(x)-x. This network architecture has been shown to avoid vanishing gradients and accuracy degradation present in traditional network architecture learning [3]. During training, this model used label smoothed [6] (smoothing=0.01) cross entropy as its loss function. |
| Short Resnet | An 18 layer Resnet followed by a dropout layer, a 512 input to 512 output linear layer, a DReLU activation function, a 512 input to 2 output linear layer, and finally a softmax layer. During training, this model used label smoothed (smoothing=0.01) cross entropy as its loss function. |
| Long Variational AutoEncoder | A 2d convolution with ReLU activation function followed by three encoder blocks encoded the embedding. Encoder blocks consisted of a spectral normalized 2d convolution layer [5], followed by 2d batch normalization and a leaky ReLU activation function. The decoder consisted of 4 decoder blocks made up of a transposed 2d convolution followed by 2d batch normalization and a leaky ReLU activation function. Self attention layers were added in between both encoder and decoder blocks [9]. Binary classification of binder vs. nonbinder was performed on each embedding by two fully connected layers (sizes 128 and 64, consisting of: a dropout layer, linear layer, 1d batch norm, and a leaky ReLU) followed by a dropout layer, linear layer (size 2), and a final softmax layer. Similarly mu and logvar were generated by two fully connected layers and a final layer (sizes 128, 112, 100). Variational AutoEncoders are generative models designed to sample across a continuous latent space[4]. During training this model used label smoothed (smoothing=0.01) cross entropy on the predictions and symmetric MSE loss on the decoder's reconstruction. The loss functions were mixed for the total training loss. |
| Short Variational AutoEncoder | A 152 layer Resnet encoder and 2 decoder blocks (separated by an attention layer). A 512 input to 2 output linear layer was trained on each embedding with a log softmax layer on the end to predict a binding vs. nonbinding result. During training this model used label smoothed (smoothing=0.01) cross entropy on the predictions and symmetric MSE loss on the decoder's reconstruction. The loss functions were mixed for the total training loss. |
| Siamese | A Siamese network trained on pairs of sequences to discriminate between binder-binder pairs and nonbinder-binder pairs. The Siamese network used here consisted of a single resnet made up of 4 layers to a 512 input to 256 output linear layer, a sigmoid activation function, and a 256 input to 2 output linear layer following. Each iteration was run individually on pairs of sequences [7]. The Euclidean distance between the resulting embeddings is used to assign our binary classification value. During training this model used contrastive loss as its loss function. |

Table S2: Descriptions of all DNN models used in this work.

#### S4.4 DNN Results

To compare performance of our DNN models, we assessed the accuracy of each model to predict a binder/nonbinder label for each experimentally validated dataset: the RBM generated dataset and the DCA generated dataset. We also calculated the F1 score metric by comparison of each model's prediction with the ground truth. The F1 score is the harmonic mean of precision, the number of true positives divided by the sum of true positives and false positives, and recall, the number of true positives divided by the sum of true positives and false negatives, in a binary classification task. A F1 score was calculated for each dataset and a mean F1 score was determined by weighting each individual F1 score by the number of total sequences in the dataset. Scores for the DNN models are provided in Table S3.

DNN (L, R, B) models (i.e. models trained on L, R or B dataset) failed to generalize to our experimental datasets. In every case, prediction of binding ability on the RBM and DCA datasets results in a significant number of false positives and false negatives. Bayes hyperparameter optimized models were directed to either minimize the loss on the validation dataset or maximize the accuracy on the validation dataset whereas AsyncHyperBandScheduler (AHSA) hyperparmater optimized models were directed to only maximize the accuracy on the validation dataset. The most accurate (L, R, B) models on the RBM generated dataset (Bayes Resnet L and Bayes Resnet L and R) were directed to minimize the loss for hyperparameter optimization and achieved 74.1% (20/27) accuracy on the RBM experimental dataset with poor performance on the DCA generated dataset at 31.3% (5/16) accuracy. A distinct correlation between optimization directive and performance metrics was observed. Models that were optimized to minimize the loss of the validation dataset performed worse in validation set accuracy, better in RBM generated dataset binder prediction, and worse in DCA generated dataset binder prediction to a significant degree than those optimized to maximize the validation set accuracy. From the confusion matrices of loss minimized models on the RBM generated dataset (Fig. S8), we see these models are completely unable to distinguish between nonbinders and binders in both the RBM generated and DCA generated datasets.

DNN (L, R, B) models trained to maximize the accuracy on the validation set performed poorly overall. The best performing of them (ASHA VAE short R) managed the highest mean F1 score, excellent accuracy on the DCA generated dataset at 87.5% (14/16) accuracy but poor performance on the RBM generated dataset with 48.1% (13/27) accuracy. The poor performance of all DNN (L, R, B) models indicates the sequencing info of the last round of selection is not sufficient for DNN models to classify sequences on their ability to bind a target.

DNN (GL, GR, GB) models were trained as a more naive classifier using good binders and randomly generated bad binders for both training and validation. As the random bad binders were guaranteed be to outside the sequence space of the entire 8th round of selection, we would expect these models to over-predict binders in our datasets which contain binders and nonbinders separated by small distances in sequence space. Indeed, all (GL, GR, GB) models have higher accuracy values than their (L, R, B) counterparts, but consistently have little to none false negatives and a large number of false positives on the sequences generated using RBM (Fig. S9). Additionally the higher accuracy scores on the RBM generated dataset and lower accuracy scores on the DCA generated dataset is due to the difference in population group membership (binder vs. nonbinder) of the two datasets. Their ability to predict thrombin binding ability from sequences close in sequence space is subpar due to their overfitting to the aptamer sequence space.

The performance of all DNN models on predicting thrombin binding ability from sequence alone was poor. DNN (L, R, B) models tend to generate a notable amount of false positives and false negatives, while (GL, GR, GB) models generate false positives almost exclusively on the RBM generated dataset. Overall, using the last round of selection for our dataset exclusively (L, R, B) or for just the good binders (GL, GR, GB) did not allow accurate prediction of thrombin binding ability from any of the DNN models.

#### S4.5 Traditional ML Training Specifics

Three traditional models: a single tree, a random forest, and a gradient-boosted forest were used to classify the experimental dataset as binders or nonbinders. The training and validation datasets used were the same as those used for the deep learning models. The scikit-learn python library implementations of each of the three models were used in this work.

#### S4.6 Traditional ML Results

Our traditional ML techniques' performance was measured by the same metrics as for our DNN models, namely the accuracy on the RBM generated sequences, the accuracy on the DCA generated sequences, and the F1 mean of both datasets shown in Table S4.

Traditional (L, R, B) models very rarely predicted a nonbinder correctly in our RBM generated dataset, instead predicting almost every sequence to be a binder. Their validation set accuracy never crossed 30%. Similar to our DNN models, the accuracy on the validation sets of the (GL, GR, GB) models was significantly better than (L, R, B) models due to the difference in sequence space of the bad binders. Traditional (GL, GR, GB) models suffered from the same issue of an overabundance of false positives including the single tree models which had the best performance of any machine learning model besides the RBM. The GR single tree achieved an accuracy of 85.2% (23/27) on the RBM generated dataset and an accuracy of 81.3% on the DCA generated dataset. Despite the high accuracy, these models suffer from the same over-fitting that the DNN (GL, GR, GB) models where binders are over-predicted significantly. The small difference in single tree models GR and GB illustrate how decreasing the amount of false positives by one in the RBM generated set has the effect of predicting almost 20% less binders in the DCA generated dataset. This ability to overestimate binders is especially apparent in the confusion matrices of the random forest (GL, GR, GB) models Fig. S10. The random forest on average performed worse than the single tree, performing as well as most DNN models. This is in stark contrast to our gradient boosted classification tree which performed poorly on every dataset no matter the hyperparameters tried.

#### S4.7 Additional ML Results

The main results for the DNN models and traditional ML models referenced in the main text are shown in Table S3 and Table S4 respectively. Fig. S7 shows the AUC, several binary performance metrics, and the performance diagram for the VAE Long ASHA model in (a-c) respectively, for the six training data sets. Additional ML results in the form of confusion matrices of each model's performance on the RBM-generated sequence dataset are included in Figs. S9, S8 and S10.

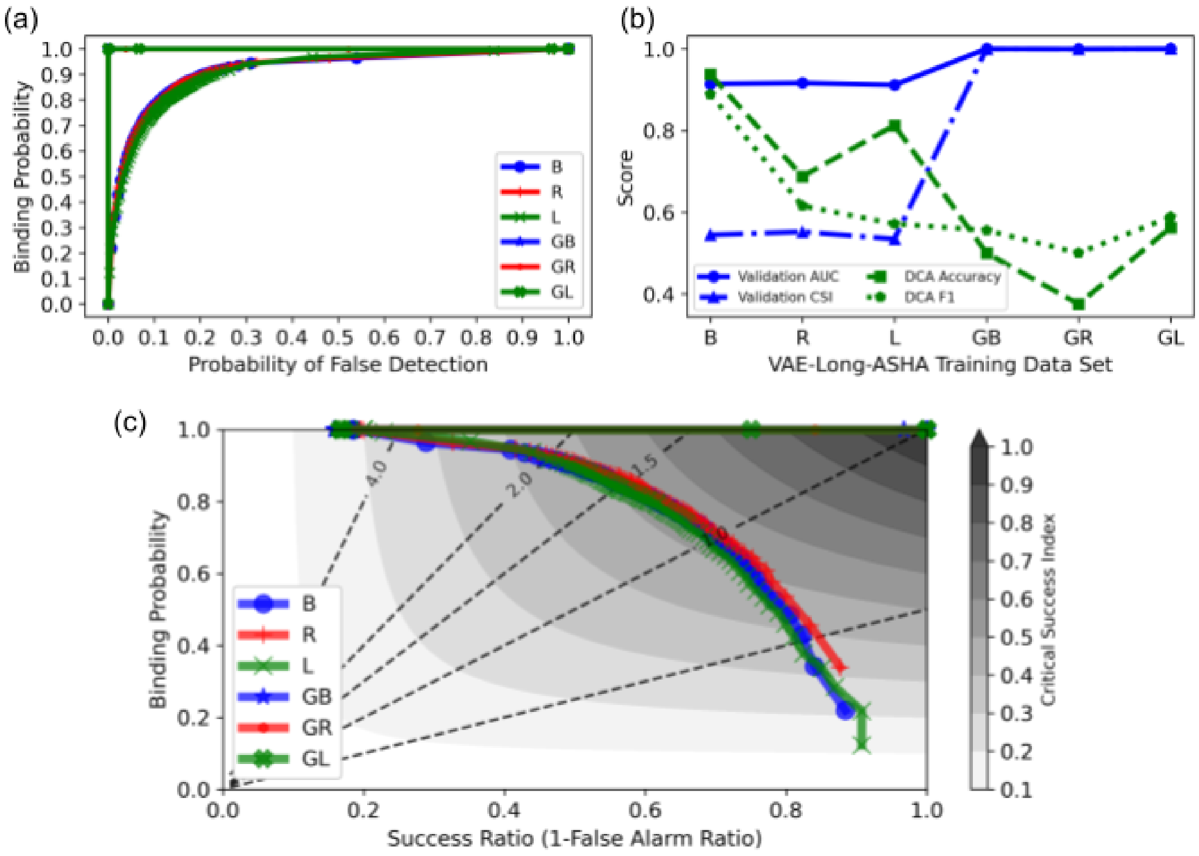


Figure S7: AUC (panel (a)), performance metrics (panel (b)), performance diagram (panel (c)) showing CSI for the VAE Long ASHA model.

| Model | Validation Acc. | RBM Acc. | DCA Acc. | F1 mean |
|---|---|---|---|---|
| **AHSA Resnet Long** | | | | |
| L | 0.792 | 0.333 | 0.750 | 0.428 |
| R | 0.711 | 0.296 | 0.562 | 0.469 |
| B | 0.751 | 0.444 | 0.625 | 0.578 |
| GL | 0.999 | 0.778 | 0.375 | 0.718 |
| GR | 0.999 | 0.889 | 0.562 | 0.790 |
| GB | 0.998 | 0.778 | 0.438 | 0.729 |
| **Bayes Resnet Long** | | | | |
| L* | 0.304 | 0.741 | 0.312 | 0.697 |
| R* | 0.281 | 0.704 | 0.312 | 0.683 |
| B* | 0.280 | 0.741 | 0.312 | 0.697 |
| **AHSA Resnet Short** | | | | |
| L | 0.758 | 0.333 | 0.688 | 0.463 |
| R | 0.789 | 0.407 | 0.750 | 0.586 |
| B | 0.767 | 0.407 | 0.875 | 0.617 |
| GL | 0.998 | 0.778 | 0.438 | 0.729 |
| GR | 0.999 | 0.778 | 0.438 | 0.729 |
| GB | 0.999 | 0.852 | 0.562 | 0.777 |
| **Bayes Resnet Short** | | | | |
| L* | 0.384 | 0.741 | 0.312 | 0.697 |
| R | 0.796 | 0.444 | 0.688 | 0.528 |
| B | 0.758 | 0.333 | 0.625 | 0.470 |
| **AHSA VAE Long** | | | | |
| L | 0.828 | 0.333 | 0.812 | 0.416 |
| R | 0.837 | 0.333 | 0.625 | 0.492 |
| B | 0.819 | 0.333 | 0.875 | 0.510 |
| GL | 1.000 | 0.815 | 0.562 | 0.766 |
| GR | 1.000 | 0.778 | 0.500 | 0.741 |
| GB | 1.000 | 0.778 | 0.562 | 0.754 |
| **Bayes VAE Long** | | | | |
| L | 0.804 | 0.307 | 0.688 | 0.401 |
| R | 0.838 | 0.407 | 0.688 | 0.505 |
| B | 0.829 | 0.296 | 0.750 | 0.450 |
| **AHSA VAE Short** | | | | |
| L | 0.757 | 0.296 | 0.688 | 0.365 |
| R | 0.789 | 0.481 | 0.875 | 0.623 |
| B | 0.776 | 0.407 | 0.812 | 0.574 |
| GL | 1.000 | 0.741 | 0.562 | 0.739 |
| GR | 1.000 | 0.889 | 0.688 | 0.822 |
| GB | 1.000 | 0.889 | 0.562 | 0.790 |
| **Bayes VAE Short** | | | | |
| L | 0.804 | 0.333 | 0.562 | 0.360 |
| R | 0.803 | 0.370 | 0.812 | 0.542 |
| B | 0.801 | 0.407 | 0.875 | 0.581 |
| **PBT Siamese** | | | | |
| L | 0.687 | 0.458 | 0.662 | 0.300 |
| R | 0.598 | 0.491 | 0.600 | 0.391 |
| B | 0.643 | 0.467 | 0.508 | 0.314 |

Table S3: Accuracy Scores for all models trained on the Left Arm (L), Right Arm (R) Both Arms (B), Generated Left Arm (GL), Generated Right Arm (GR) or Generated Both Arms (GB) datasets. Models with a star(*) were optimized to minimize the validation set loss. Validation sets were taken as 10% of the training data, while the experimental datasets consisted of the 27 RBM generated sequences and the 16 DCA generated sequences.

| Model | Validation Acc. | RBM Acc. | DCA Acc. | F1 Mean |
|---|---|---|---|---|
| **Single Tree** | | | | |
| L | 0.116 | 0.778 | 0.313 | 0.708 |
| R | 0.122 | 0.697 | 0.313 | 0.741 |
| B | 0.114 | 0.778 | 0.375 | 0.718 |
| GL | 0.999 | 0.704 | 0.813 | 0.764 |
| GR | 0.999 | 0.852 | 0.813 | 0.832 |
| GB | 0.885 | 0.889 | 0.625 | 0.781 |
| **Random Forest** | | | | |
| L | 0.242 | 0.630 | 0.438 | 0.665 |
| R | 0.270 | 0.630 | 0.313 | 0.651 |
| B | 0.294 | 0.630 | 0.313 | 0.651 |
| GL | 0.950 | 0.741 | 0.375 | 0.684 |
| GR | 0.942 | 0.778 | 0.375 | 0.695 |
| GB | 0.939 | 0.778 | 0.438 | 0.706 |
| **Gradient Boosted Forest** | | | | |
| L | 0.098 | 0.741 | 0.313 | 0.697 |
| R | 0.097 | 0.741 | 0.313 | 0.697 |
| B | 0.099 | 0.741 | 0.313 | 0.697 |
| GL | 0.091 | 0.741 | 0.313 | 0.697 |
| GR | 0.091 | 0.741 | 0.313 | 0.697 |
| GB | 0.167 | 0.741 | 0.313 | 0.697 |

Table S4: Accuracy Scores for single tree, random forest and gradient boosted forest trained on the Left (L), Right (R), Both (B), Generated Left Arm (GL), Generated Right Arm (GR) or Generated Both Arms (GB) datasets. Validation sets were taken as 20% of the training data, while the experimental dataset consisted of the 27 RBM generated sequences and the 16 DCA generated sequences.

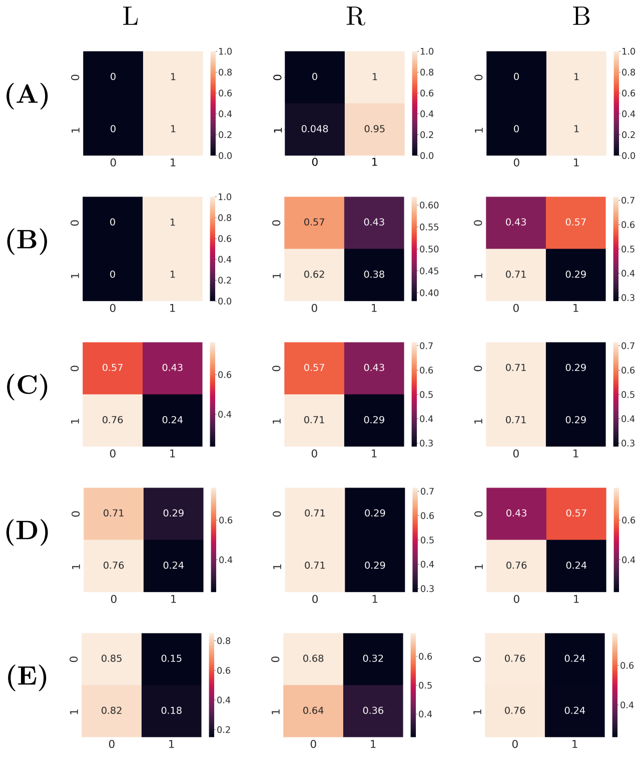


Figure S8: Confusion Matrices of trained loss-minimized or accuracy maximized bayesian optimized hyper-parameters on RBM generated dataset, (A) Long Resnet, (B) Short Resnet, (C) Short VAE, (D) Long VAE, and Population based training of Siamese Network (E). Predicted Label (0) nonbinder or (1) binder is shown on the x-axis with the true label being the y-axis.

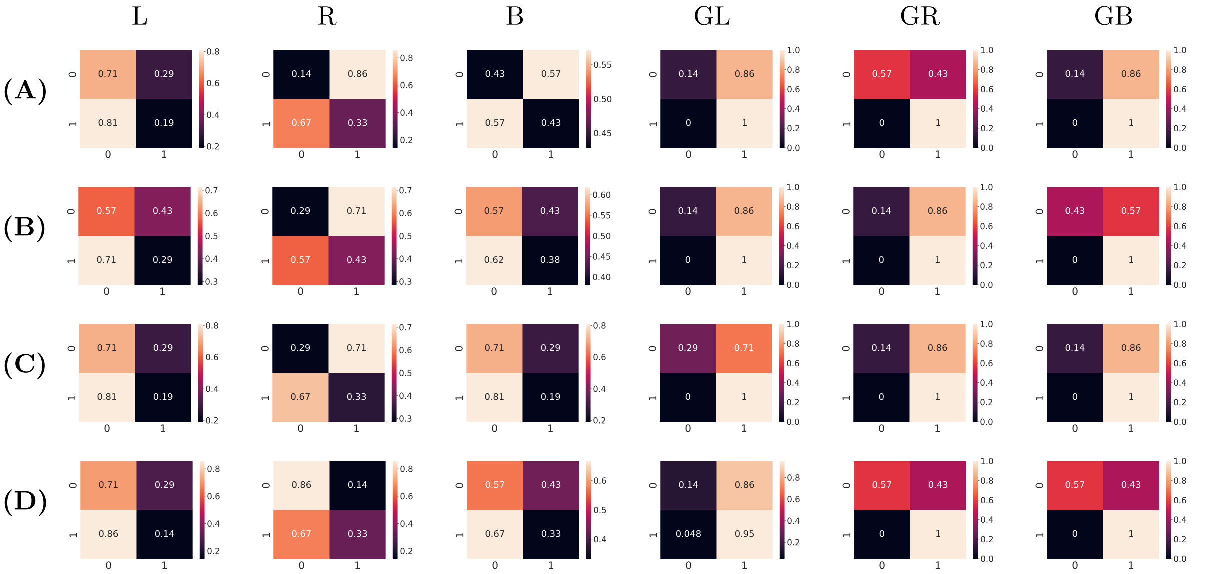


Figure S9: Confusion Matrices of accuracy maximized ASHA scheduler for hyper-parameter optimization using deep learning models: Long Resnet (A), Short Resnet(B), Long VAE (C) and Short VAE (D) on the RBM generated dataset. Predicted Label (0) nonbinder or (1) binder is shown on the x-axis with the true label being the y-axis.

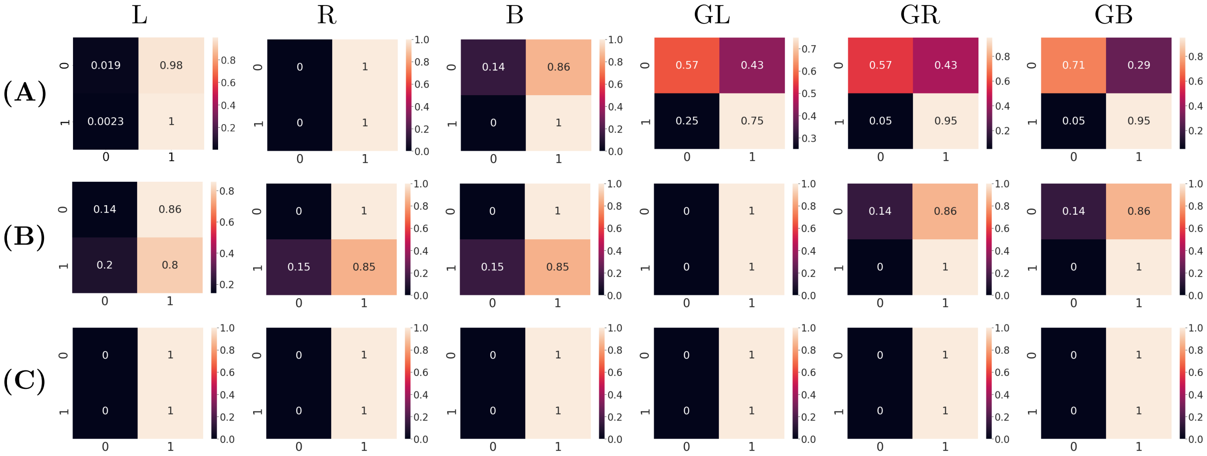


Figure S10: Confusion Matrices of traditional machine learning models: Single Tree, Random Forest and Gradient Boosted Forest on the RBM generated dataset. Predicted Label (0) nonbinder or (1) binder is shown on the x-axis with the true label being the y-axis.

### S5 Direct Coupling Analysis

Direct Coupling Analysis (DCA) is a method of analysis originally used for contact prediction in proteins from sequence alignments of homologues. The basis of this method is that the homologue alignments have the same general native state to carry out their function. Despite their differences in sequence, all homologues will have similar inter-domain contacts. To maintain these contacts, detrimental single site mutations must be offset by compensatory mutations in other parts of the sequence. DCA is a maximum-entropy method, where the model parameters are fixed so that the one- and two-point correlations along the sequences are fixed to those observed in the training homologue-sequence aligment. The sequence probability is given in Eq. (S9) and is dependent on the learned single position parameters ($h$) and pairwise interactions ($J_{ij}$) of the multiple sequence alignment.

$$P(\sigma) = \frac{1}{Z} \exp \left( \sum_{i=1}^{L} h_i(\sigma_i) + \sum_{1 \leq i < j \leq L} J_{ij}(\sigma_i, \sigma_j) \right). \tag{S9}$$

Similar to the protein case, we applied DCA on our aligned DNA aptamer dataset to approximate the aptamer sequence space with the learned single site and pairwise correlations. Sequences unobserved in the original dataset were generated from the learned parameters and tested experimentally.

#### S5.1 DCA Training

The training set used for DCA analysis was a subset (90%) of sequences with copy number $> 1$ from the 8th round of selection. Rather than separate the arms of each nanotile, the DCA model was trained with on 40 nt long sequences containing both arms. The normalization constant Z is difficult to calculate, so we use psuedolikelihood maximzation DCA (plmDCA) [2] to obtain local fields ($h_i$) and pairwise coupling ($J_i j$) for the model, given the aligned aptamer dataset. Monte Carlo sampling was applied across a range of temperatures and mutation steps to sample from the learned parameters. In total $2{*}10^9$ sequences were sampled, and from those 16 sequences shown in Table S5 were selected for experimental validation of the model.

| Label | Sequence | Score | Binder Prediction | Experimental result |
|---|---|---|---|---|
| d1 | AGGGTAGGTGTGGGGTATGC | 86.92 | B | NB |
| d2 | AGGGTAGATGTGTAGGATGC | 87.86 | B | NB |
| d3 | AGGGATGATGGTTGGTAGGC | 84.76 | B | NB |
| d4 | AGGGATGATGTGGATTAGGC | 86.03 | B | NB |
| d5 | AGGGTGGGAGCGGGGGACGC | 75.01 | B | NB |
| d6 | CGGGTAGGTGTGGATTATGC | 77.59 | B | NB |
| d7 | GTAGGACGGGTAGGGCGGTC | 67.57 | NB | NB |
| d8 | GGGGGTTGGGCGGGATGGGC | 72.15 | B | NB |
| d9 | GCGGGTTGGGCAGGATCAGC | 44.58 | NB | NB |
| d10 | AGGGATGATGTGTGGTAGGC | N/A | Cntrl | Cntrl |
| d11 | GTAGGATGGGTGGGGTGGGA | 86.46 | B | B |
| d12 | GTAGGATGGGTAGGGTGGTA | 84.76 | B | B |
| d13 | CTAGGTTGGGTAGGGTGGTG | 75.01 | B | B |
| d14 | CTAGCATGGGTAGGGTGGTG | 77.59 | B | B |
| d15 | GTAGCATGGGTAGGGTGGTC | 65.57 | NB | NB |
| d16 | TTGGGTGGTGTAGGTTGGCG | 72.15 | B | B |
| d17 | TTGGGTGGTGCAGGTTCGCG | 44.58 | NB | NB |
| d18 | CTAGGATGGGTAGGGTGGTG | N/A | Cntrl | Cntrl |

Table S5: Result of thrombin binding assays with all DCA-generated sequences and sequences of exosite I control d18 and exosite II control d10. B indicates a binder while NB indicates a nonbinder.

#### S5.2 DCA Sequence Selection

From the generated sequences, we wanted to find not only novel binders but also verify the learned model parameters. Sequences are scored according to the sum of their single position and pairwise parameters.

A sequence's higher score indicates it is more likely to bind while a lower score indicates it is less to bind. Predicted binders (sequences d1, d2, d3, d4, d5, d11, d12, d13) were selected from the MC-generated sequences by having the highest score while being at least 3 mutations away from anything observed in the entirety of the 8th round of sequencing data. Two predicted nonbinders (d6, d14) were selected for having the lowest score within 2 mutations of the dataset. Rationally designed binders (d7, d8, d9, d15, d16, d17) were generated by randomly selecting a good and bad binder from the original dataset and altering them to either have the highest or lowest score possible by exhaustively calculating the entire sequence space within 3 mutations and finding the variant with the highest or lowest score. Model parameters used to generate all sequences are shown in Fig. S11.

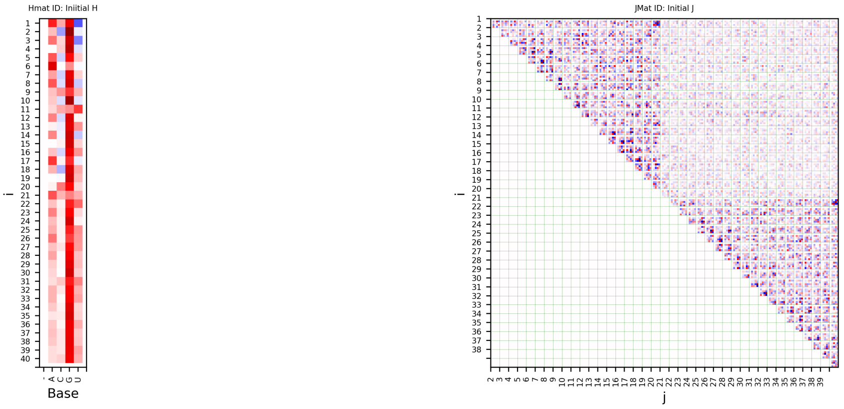


Figure S11: Single position ($H$) and pairwise correlations ($J_{ij}$) learned by the plmDCA model and used in both sampling and sequence selection.

#### S5.3 DCA Gel Shift Assay

Sequences generated using the plmDCA model were tested experimentally for their ability to bind Thrombin. Binding sequences formed a clear protein / stem-loop band. Sequences were tested the same way as done for the RBM-generated sequences in the main text. Fig. S12 shows the experimental results of a gel shift assay for the plmDCA generated sequences.

#### S5.4 DCA Binding Site Assay

Thrombin binding sequences generated via plmDCA (d11, d12, d13, d14, d16) were tested against known binders 5' 6FAM labeled ThA and ThD to determine their binding site as described in the main text. Table S6 contains the results and Fig. S12 shows the gel results.

| Label | Sequence | Binding Site |
|---|---|---|
| d11 | GTAGGATGGGTGGGGTGGGA | exosite I |
| d12 | GTAGGATGGGTAGGGTGGTA | exosite I |
| d13 | CTAGGTTGGGTAGGGTGGTG | exosite I |
| d14 | CTAGCATGGGTAGGGTGGTG | exosite I |
| d16 | TTGGGTGGTGTAGGTTGGCG | exosite I |

Table S6: Exosite prediction of DCA sequences that bound thrombin from our gel shift assays.

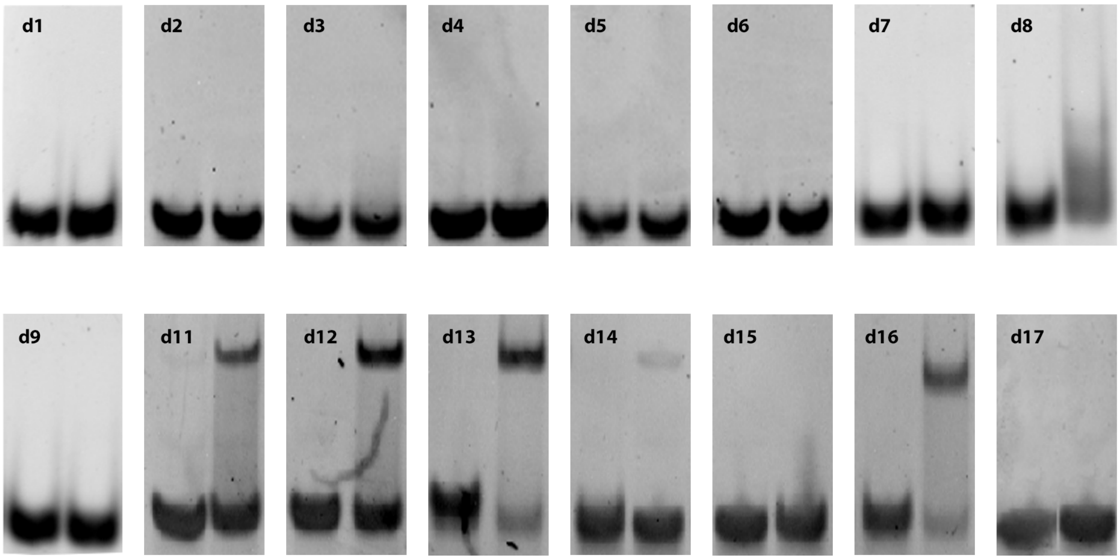


Figure S12: Thrombin binding assay of DCA generated sequences. Lane 1 has the stem loop alone, whereas lane 2 has the same stem loop exposed to thrombin. Binding sequences are indicated by a high visible band in lane 2.

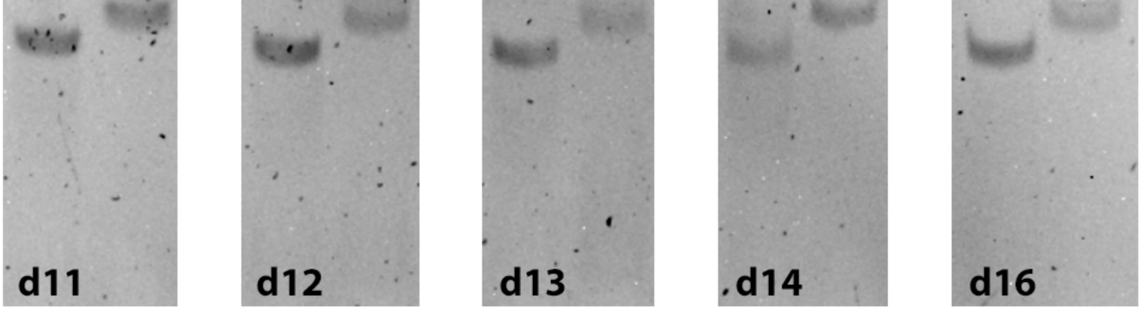


Figure S13: Binding site assay using the same method discussed in the main text. Lane 1 is the result of the preincubated strand exposed to exosite-II binder ThA and lane 2 is the preincubated strand exposed to exosite-I binder ThD.

#### S5.5 DCA Results

The weak pairwise correlations seen in the top right corner of the pairwise correlation matrix ($J_{ij}$) confirm the lack of correlation between the two arms of each nanotile. The plmDCA method did see limited success in generating novel binders (d11, d12, d13, d16) from the right loop sequences but no success in generating binding left loop sequences (d1, d2, d3, d4, d5) (Fig. S13).

We tried also to train a DCA using the same algorithm we used for the RBM models to obtain the model parameters, i.e. the persistent contrastive divergence algorithm. Moreover, building on the results obtained with our RBM models, we decided to use all the available sequences to train the BM model, neglecting the counts. Then we compared, for the obtained DCA model trained with single-loop sequences at round 8, the log-likelihood assigned by the DCA with the one assigned by an RBM trained on the same data. The resulting plot is given in Suppl. Fig. S14, and this test gave a very good linear correlation between the log-likleihoods of the two models (slope of the linear fit: 1.09; $R^2$ score: 0.97), suggesting that the DCA model trained with persistent contrastive divergence has superior generalization capabilities with respect to plmDCA models. This result is compatible with what observed in [1].

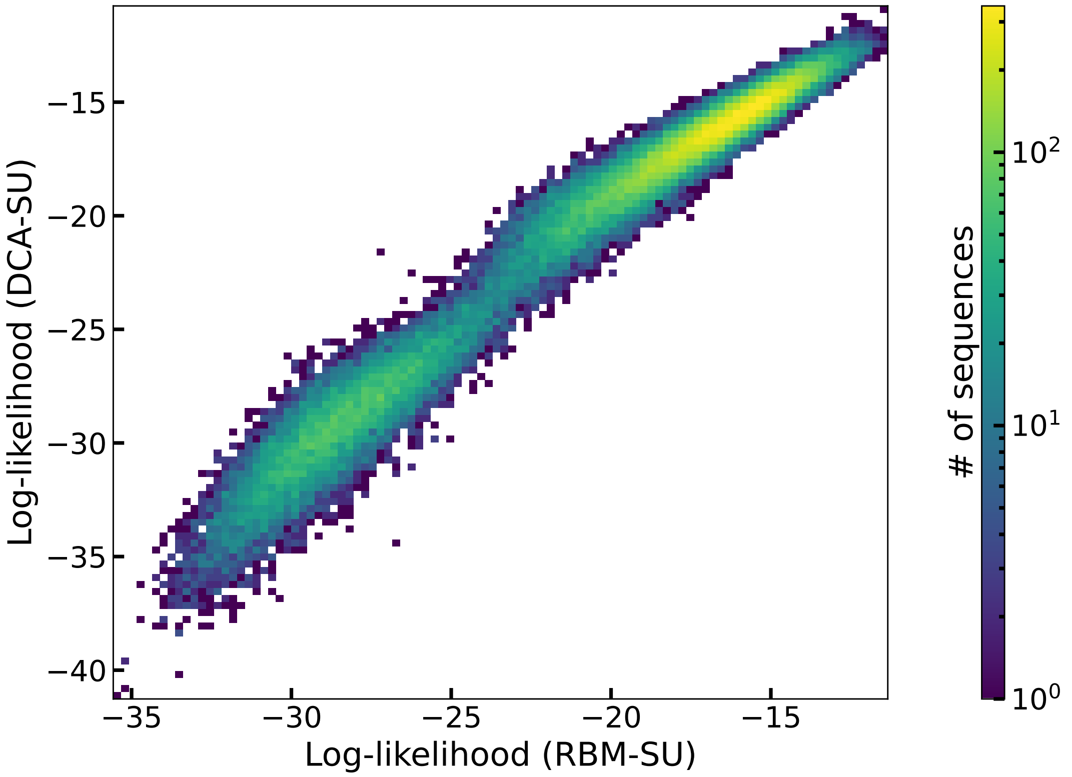


Figure S14: Log-likelihood of all unique single-loop aptamers observed at round 6, as computed by a DCA and an RBM model trained through persistent contrastive divergence. The corresponding linear fit resulted in a slope of 1.09 and an $R^2$ of 0.97.

#### S6 Additional supplementary figures and tables

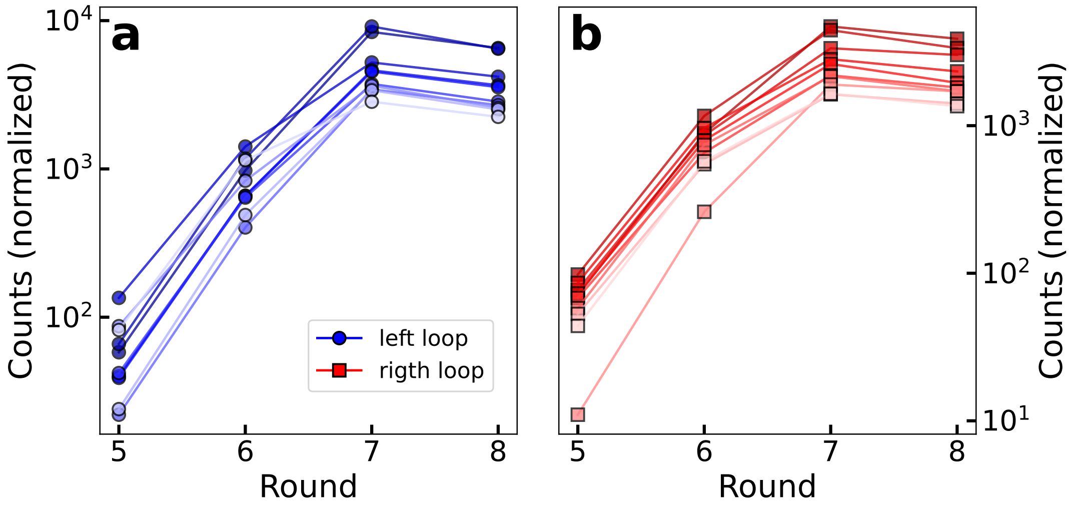


Figure S15: Evolution of counts of the 10 left (panel a) and right (panel b) aptamers with largest number of counts at round 8. Counts have been re-scaled by a factor so that the total number of counts in each round is constant.

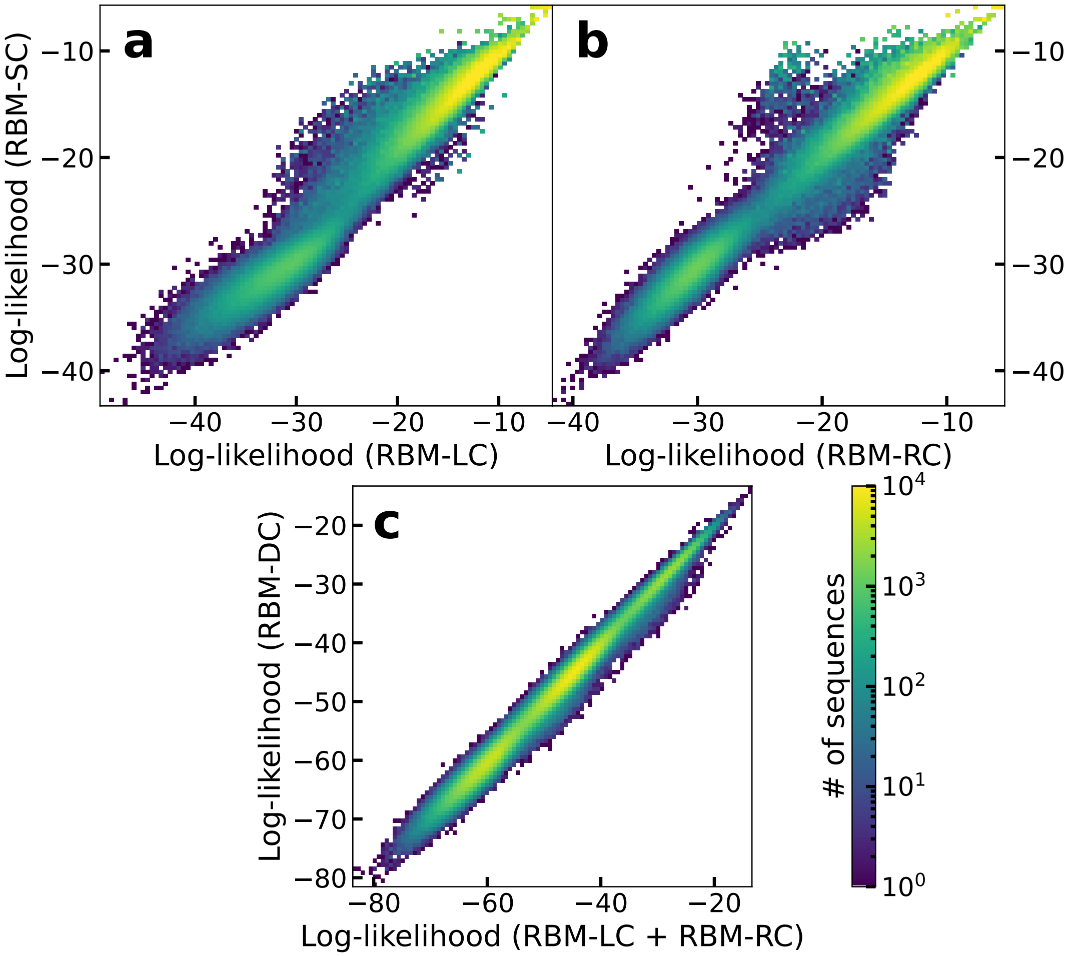


Figure S16: Log-likelihood computed with the RBM-SC model and with the RBM-LC model (trained on left single-loop sequences at round 8, see S3) in panel a or RBM-RC model (trained on right single-loop sequences at round 8, see S3) in panel b for the single-loop sequences observed at round 8. The slope and the $R^2$ values of the linear fit are respectively 0.96 and 0.98 for panel a, and 1.05 and 0.97 for panel b.

Panel c: log-likelihood computed with the RBM-DC model for the double-loop sequences observed at round 5, compared with the sum of the log-likelihood obtained by using RBM-LC to score the left loop and RBM-RC to score the right loop. The slope and the $R^2$ value of the linear fit are, respectively, 0.99 and 0.99.

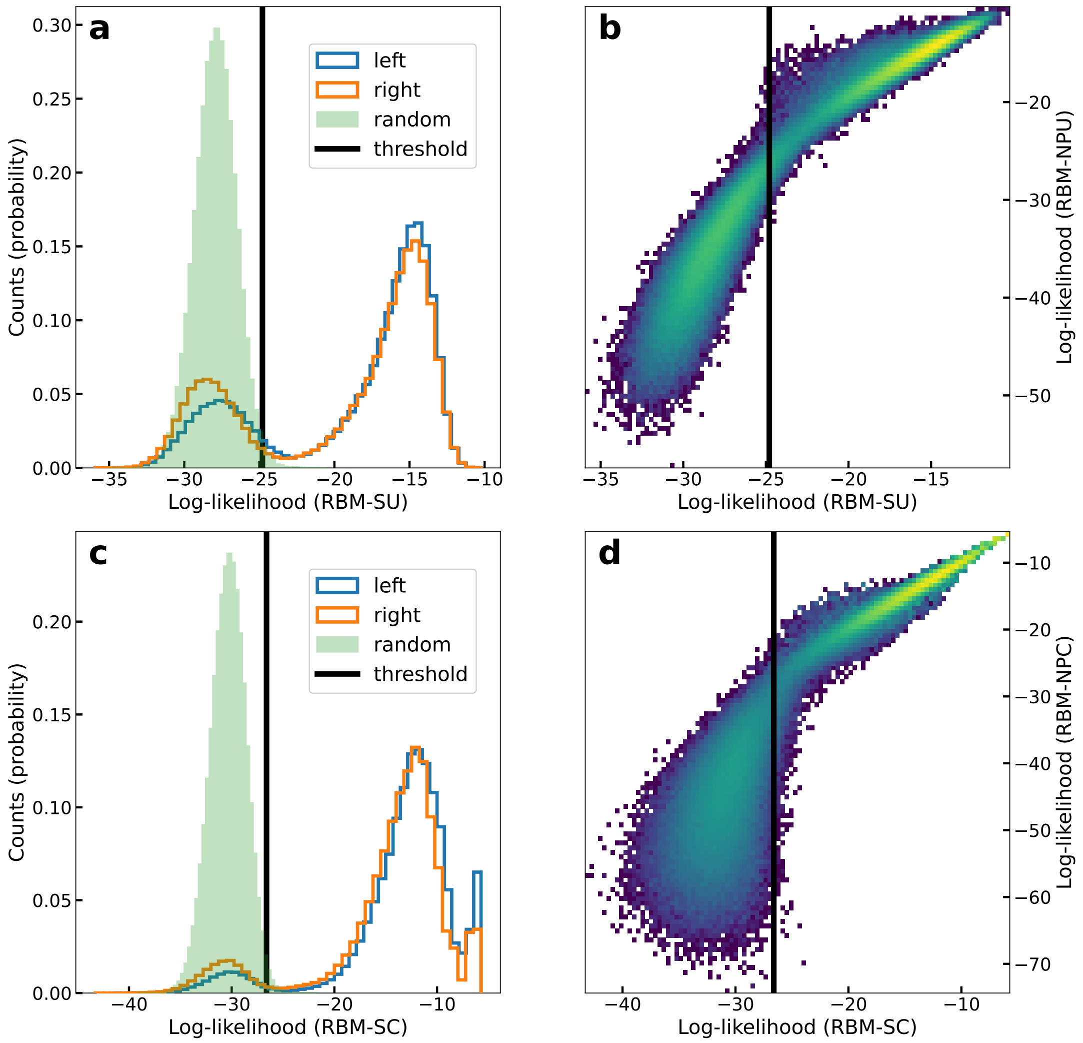


Figure S17: Left side: histograms of log-likelihoods of left (blue) and right (orange) loops computed with RBM-SU (panel a) or RBM-SC (panel c) for sequences observed in round 8 (unique in panel a, with their counts in panel b), together with that of $5 \cdot 10^5$ random uniform sequences (light green); the black line is the 99-quantile of the light green histogram, and parasite sequences are defined as those which have lower log-likelihood than the black line, while at the same time the other loop of the 40-nt aptamer has log-likelihood larger than the threshold.
Right side: log-likelihood of the RBM trained after excluding parasite sequences at round 8 (RBM-NPU for panel b, RBM-NPC for panel d) versus that of the RBM-SU (panel b) or RBM-SC (panel d) model. A linear fit for the points at the right-hand side of the black line (which is the same of panels a for panel b, and of panel c for panel d) gives a slope of 1.0 and a $R^2$ of 0.92 for panel b, and a slope of 1.0 and a $R^2$ of 0.96 for panel d. For points at the left-hand side of the black line the slope is 2.6 with an $R^2$ of 0.79 for panel b, and the slope is 2.0 with an $R^2$ of 0.33 for panel d.

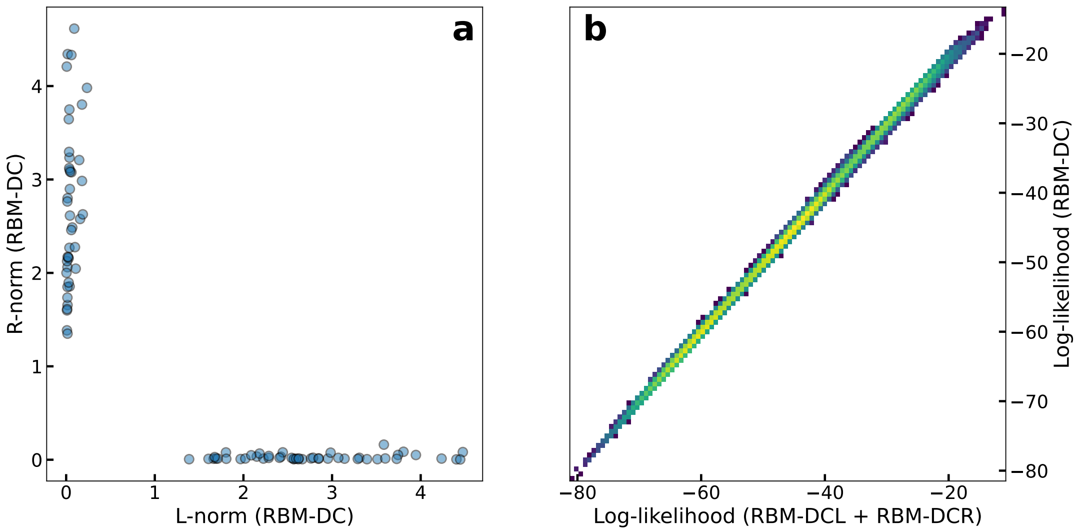


Figure S18: Panel a: Frobenius norms obtained for each weight of RBM-DC computed using only the first 20 visible units (L-norm in the x axis) or the last 20 visible units (R-norm in the y axis).
Panel b: RBM-DCL and RBM-DCR are two RBMs with 20 visible units used to score left and right loops. RBM-DCL (RBM-DCR) is obtained from RBM-DC by using only its first (last) 20 visible units and their fields, and the hidden units with L-norm > R-norm (R-norm > L-norm) with their potentials, ignoring their interactions with the last (first) 20 visible units. In this panel, we compare, for each unique double-loop sequence observed at round 5, the log-likelihood of the RBM-DC model with the sum of the log-likelihoods obtained by using RBM-DCL to score the left loop and RBM-DCR to score the right loop. The slope of the linear fit is 0.99 and the $R^2$ score is $> 0.99$.

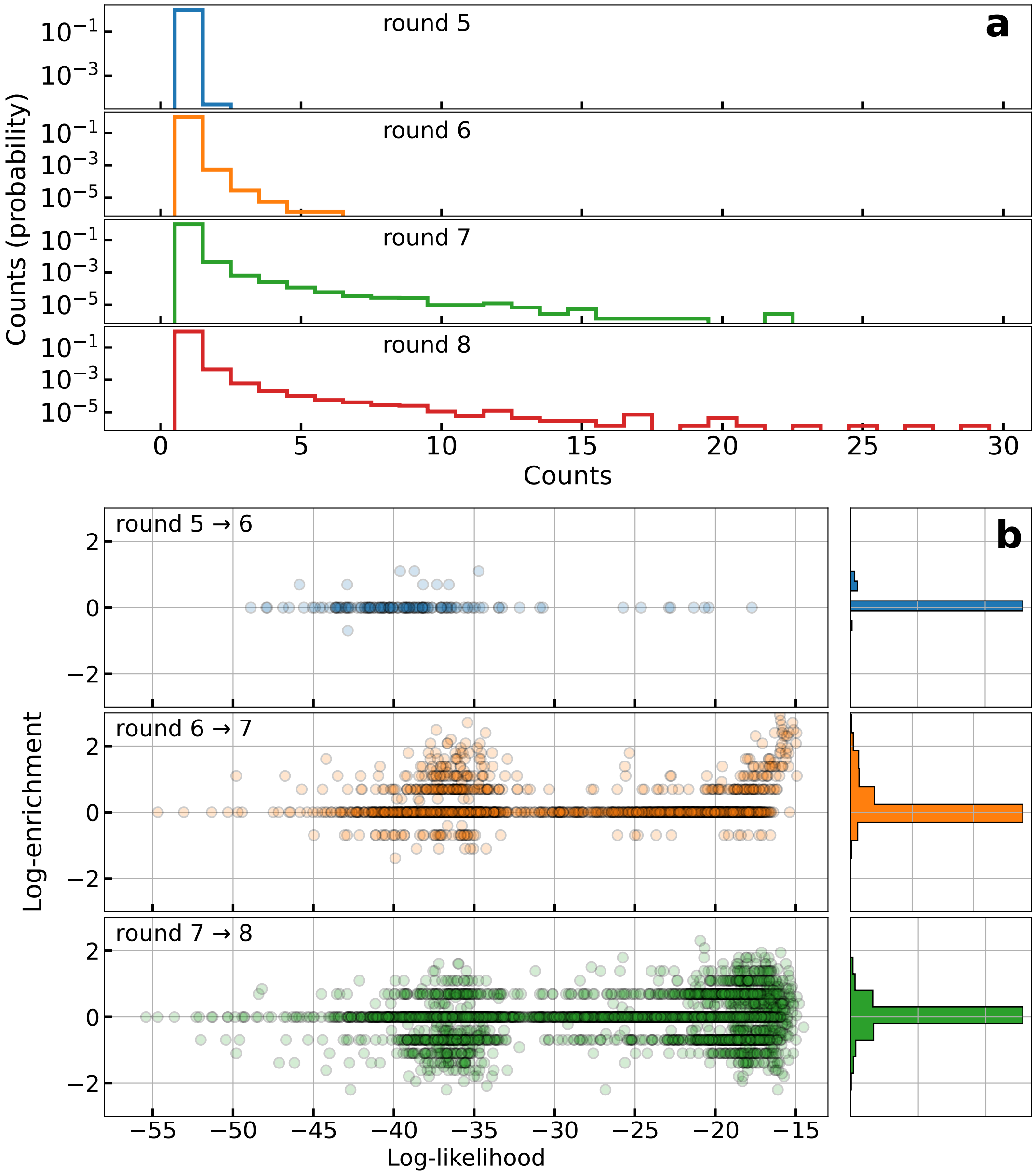


Figure S19: Panel a: probability density function of the counts observed for the double aptamers in each round. Notice the log scale on the y axis.
Panel b: for each pair of consecutive rounds, we plot here the logarithm of the ratio of counts of the sequences present in both rounds (left) and the corresponding histogram (right), against the log-likelihood of the sequence computed with the RBM-DC model.

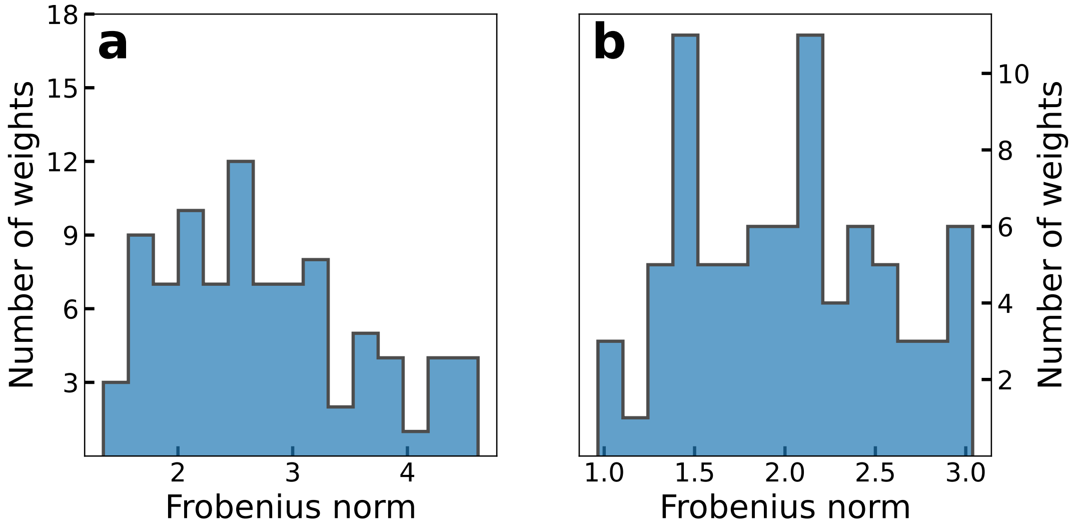


Figure S20: Panel a: Frobenius norms of the weights for RBM-DC. The logos corresponding to the 3 weights with largest Frobenius norm are given in Fig. 4a-c.
Panel b: Frobenius norms of the weights for RBM-SC. The logos corresponding to the weight with the 2nd largest Frobenius norm and the one with the 7th largest Frobenius norm are given in Fig. 4e-f.

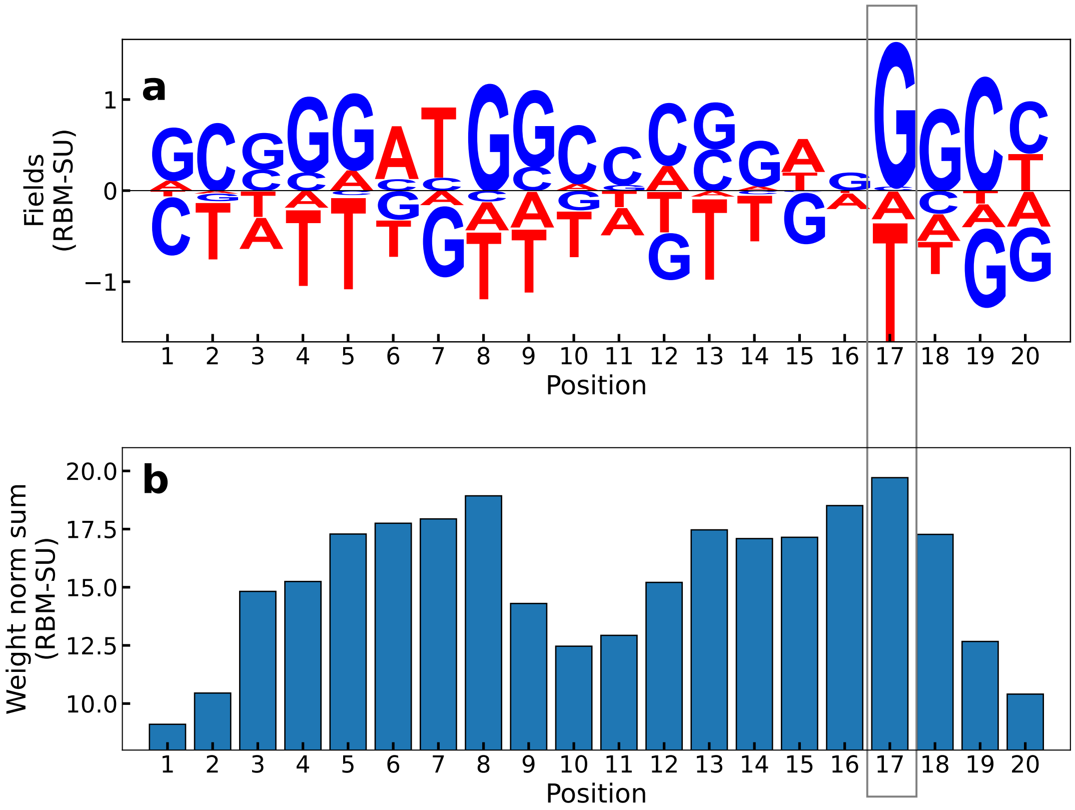


Figure S21: Panel a: fields of RBM-SU. The largest field (in norm) corresponds to position 17 (gray box), which is the one that in Fig. S25 determines the binding exosite.
Panel b: sum of the norms of each weight of RBM-SU, at fixed sequence position. The largest sum corresponds again to position 17 (gray box).

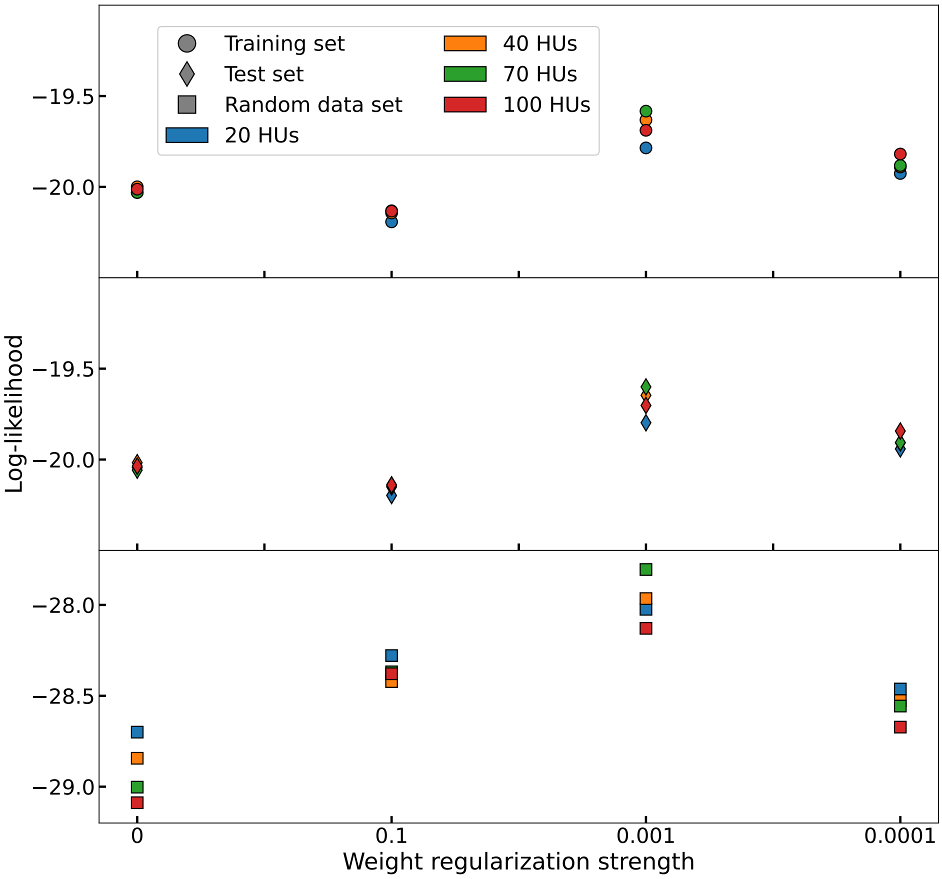


Figure S22: Average log-likelihoods computed with 16 RBMs trained with different choices of hidden unit numbers and weight regularization on the single aptamers obtained from the 8-th round. The scale on the y-axis is kept constant across the different sub-plots to highlight how the difference in average log-likelihoods are much smaller than the difference between the log-likelihood of training (and test) data and that of random sequences. The green circle at 0.001 regularization strength correspond to the RBM used in the paper (RBM-SU).

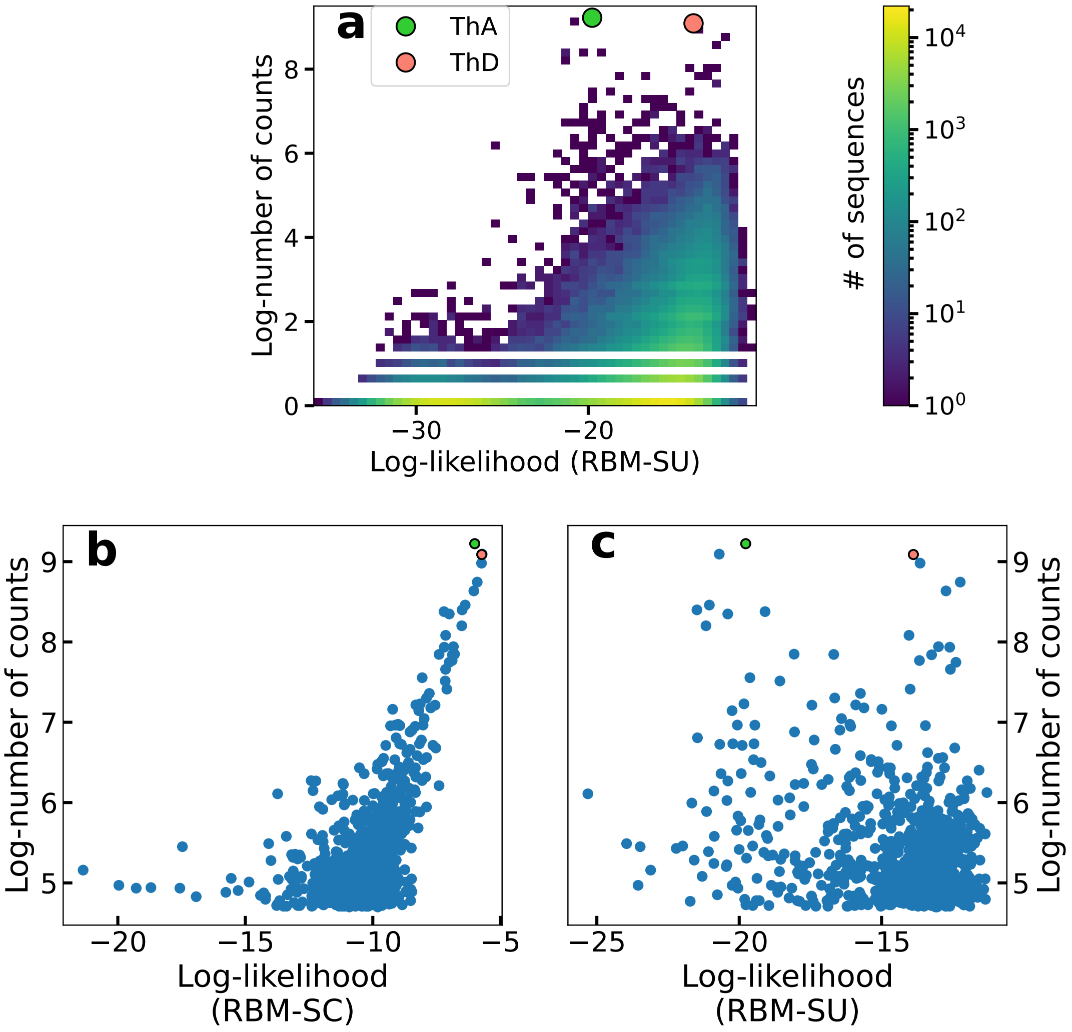


Figure S23: Panel a: Log-likelihoods (computed with RBM-SU) versus log number of counts for the unique single-loop sequences observed at round 8. ThA (counts: 10132, log-likelihood: -19.8) and ThD (counts: 8853, log-likelihood: -13.9) are highlighted with circles.
Panels b, c: Log-likelihoods computed with RBM-SC (for panel b) or RBM-SU (for panel c) versus log number of counts for the 1000 unique single-loop sequences observed at round 8 with highest number of counts.

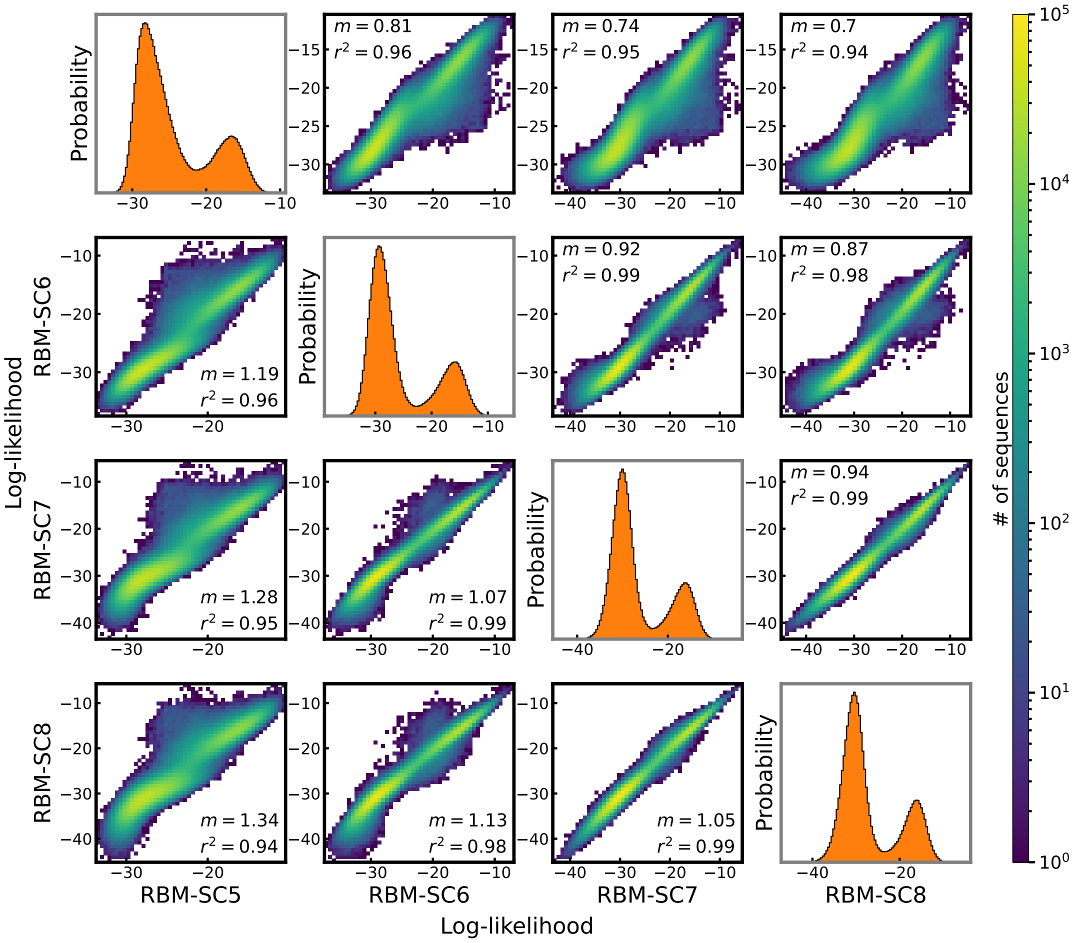


Figure S24: Comparison of the log-likelihoods computed with RBM-SC trained at different rounds (named RBM-SC5, RBM-SC6, RBM-SC7 and RBM-SC8 if trained respectively on sequences observed in round 5, 6, 7, 8). Plots on the diagonal are the distribution of the log-likelihoods of each RBM. The sequences used to prepare each histogram are the full set of sequences observed in round 5, 6, 7, or 8 (discarding counts). In each-non diagonal plot, the slope $m$ and the coefficient of determination $r^2$ for the linear fit are given.

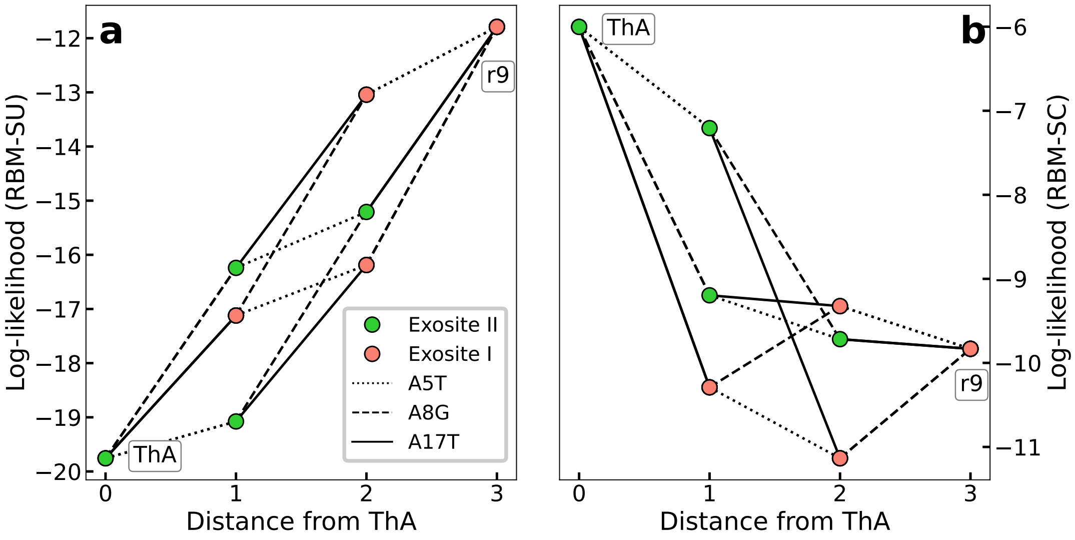


Figure S25: RBM-SU (panel a) or RBM-SC (panel b) log-likelihood versus distance from ThA for sequences p1 to p6 in Table 1. Different mutations are represented with different line styles: dotted lines for mutations involving position 5 (mutating A into T when going from ThA to r9), dashed lines for mutations involving position 8 (mutating A into G when going from ThA to r9), and solid lines for mutations involving position 17 (mutating A into T when going from ThA to r9).

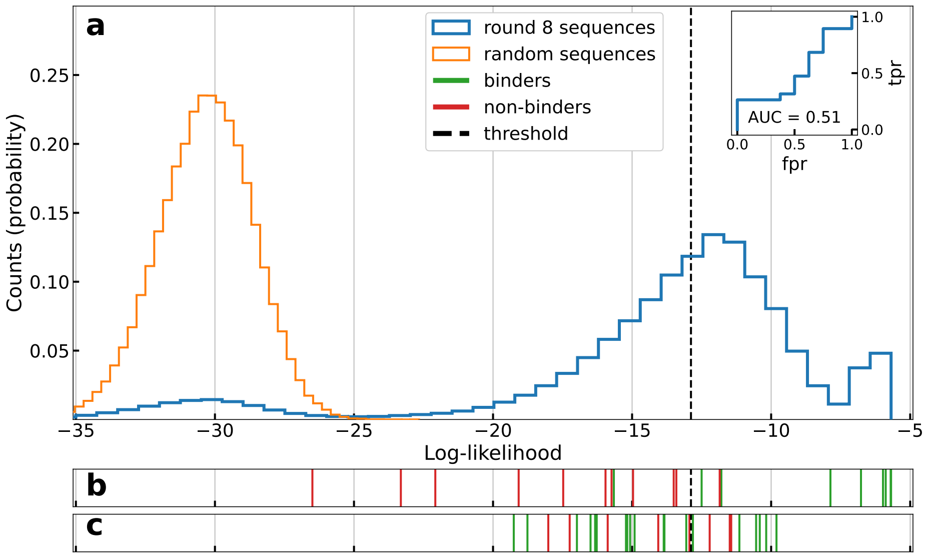


Figure S26: Panel a: Histogram of the log-likelihoods of all unique aptamers observed in the last round (blue line) and of uniformly random sequences (orange line), computed with RBM-SC trained on single-loop sequences from round 8, keeping information about the counts. Inset: AUC computed on the sequences generated by the RBM-SU model (panel c).
Panel b: Vertical lines locate the log-likelihoods of sequences experimentally validated to be binders (green) or non binders (red). Sequences taken from a preliminary set described in Suppl. Table S5. Results allows us to determine the binding/non binding threshold, shown with the black dashed line.
Panel c: same as panel b for sequences designed with the RBM-SU model, as described in Sec. 2.6 (see Table 1).

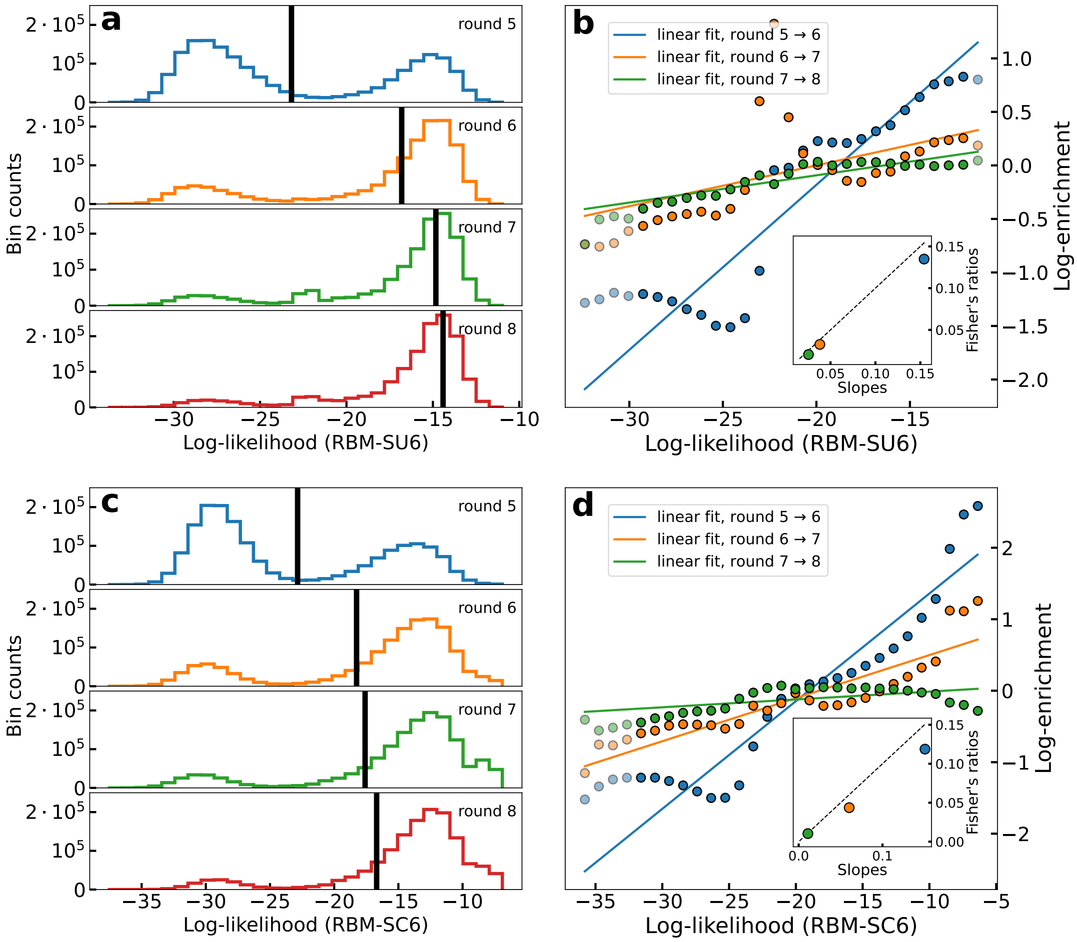


Figure S27: Relationship between log-enrichment and log-likelihoods of single-loop aptamers. Panels a, c show the histograms of log-likelihoods at each round, as computed by RBM-SU6 (panel a) and RBM-SC6 (panel c). Panels b, d show the scatter plot of log-enrichment of each bin in the left panels, and the corresponding log-likelihood. In the inset, the slope of each linear fit appearing in the main plot is compared with the same quantity estimated as a Fisher's ratio (see Sec. 2.3). The dashed black line is the $x = y$ line.

| Label | counts round 8 | Dist1 | Dist3 | Dist10 | Dist100 |
|---|---|---|---|---|---|
| r1 | 3 | 0 | 0 | 1 | 1 |
| r2 | 3 | 0 | 0 | 1 | 1 |
| r3 | 0 | 1 | 1 | 1 | 1 |
| r4 | 0 | 1 | 1 | 2 | 2 |
| r5 | 0 | 1 | 1 | 2 | 2 |
| r6 | 242 | 0 | 0 | 0 | 0 |
| r7 | 341 | 0 | 0 | 0 | 0 |
| r8 | 11 | 0 | 0 | 0 | 1 |
| r9 | 9 | 0 | 0 | 1 | 2 |
| r10 | 0 | 1 | 2 | 2 | 3 |
| r11 | 0 | 2 | 2 | 2 | 4 |
| r12 | 0 | 1 | 2 | 3 | 3 |
| r13 | 0 | 2 | 2 | 3 | 5 |
| r14 | 0 | 2 | 2 | 2 | 5 |
| r15 | 0 | 2 | 2 | 2 | 4 |
| r16 | 0 | 1 | 2 | 2 | 3 |
| r17 | 0 | 1 | 2 | 3 | 4 |
| r18 | 528 | 0 | 0 | 0 | 0 |
| r19 | 139 | 0 | 0 | 0 | 0 |
| r20 | 10 | 0 | 0 | 0 | 1 |
| r21 | 8 | 0 | 0 | 1 | 2 |
| r22 | 0 | 2 | 2 | 2 | 2 |
| r23 | 0 | 1 | 1 | 2 | 4 |
| r24 | 0 | 1 | 1 | 1 | 3 |
| r25 | 0 | 1 | 1 | 2 | 3 |
| r26 | 0 | 1 | 3 | 3 | 4 |
| r27 | 0 | 1 | 1 | 1 | 3 |

Table S7: For each sequence generated from RBM-SU trained on unique loop sequences observed in the last round, we provide here the distance from the closest single-loop aptamer observed at round 8 (column Dist1, 382094 sequences) and the number of counts of each sequence at round 8. Since a good binder is expected to be found close to a sequence with many counts, we also provide in the other columns (Dist3, Dist10, Dist100) the distance to the closest single-loop aptamer with at least, respectively, 3, 10 or 100 counts in round 8 (respectively 74785, 22332, and 1177 sequences).

| | Fig. 2c (slope) | Fig. 2c (Fisher's ratio) | Suppl. Fig. S24 |
|---|---|---|---|
| $\beta_5$ | - | - | 0.81 |
| $\beta_6$ | - | - | 1 |
| $\beta_7$ | - | - | 1.07 |
| $\beta_8$ | - | - | 1.13 |
| $\alpha_5$ | 0.16 | 0.14 | 0.19 |
| $\alpha_6$ | 0.07 | 0.06 | 0.07 |
| $\alpha_7$ | 0.01 | 0.01 | 0.05 |

Table S8: Ratios of the values $\beta_r$ (with $r = 5, 6, 7, 8$), see Eq. (3) can be estimated from the slopes given in Suppl. Fig. S24. Here we fixed the fitness scale so that $\beta_6 = 1$, or, equivalently, each value in this table is given in units of $\beta_6$. Coefficients $\alpha_r$ (with $r = 5, 6, 7$) can be obtained: (i) as the slopes in Fig. 2c (first column); (ii) using the Fisher's ratio, see inset of Fig. 2c (second column); (iii) from the slopes in Suppl. Fig. S24, since $\beta_{r+1} - \beta_r = \alpha_r$, see Eq. (3). Each of these methods has different noise sources, but the values obtained are in quite good agreement.

### References

[1] J. P. Barton, E. De Leonardis, A. Coucke, and S. Cocco. ACE: adaptive cluster expansion for maximum entropy graphical model inference. *Bioinformatics*, 32(20):3089–3097, 06 2016.

[2] M. Ekeberg, T. Hartonen, and E. Aurell. Fast pseudolikelihood maximization for direct-coupling analysis of protein structure from many homologous amino-acid sequences. *Journal of Computational Physics*, 276:341–356, 2014.

[3] K. He, X. Zhang, S. Ren, and J. Sun. Deep residual learning for image recognition. *Proceedings of the IEEE Computer Society Conference on Computer Vision and Pattern Recognition*, 2016-December:770–778, 2016.

[4] D. P. Kingma and M. Welling. Auto-encoding variational bayes. *CoRR*, abs/1312.6114, 2014.

[5] T. Miyato, T. Kataoka, M. Koyama, and Y. Yoshida. Spectral normalization for generative adversarial networks. *6th International Conference on Learning Representations, ICLR 2018 - Conference Track Proceedings*, 2018.

[6] R. Müller, S. Kornblith, and G. Hinton. When does label smoothing help? *Advances in Neural Information Processing Systems*, 32, 2019.

[7] E. van der Spoel, M. P. Rozing, J. J. Houwing-Duistermaat, P. Eline Slagboom, M. Beekman, A. J. M. de Craen, R. G. J. Westendorp, and D. van Heemst. Siamese Neural Networks for One-Shot Image Recognition. *ICML - Deep Learning Workshop*, 7(11):956–963, 2015.

[8] J. N. Zadeh, C. D. Steenberg, J. S. Bois, B. R. Wolfe, M. B. Pierce, A. R. Khan, R. M. Dirks, and N. A. Pierce. NUPACK: Analysis and design of nucleic acid systems. *Journal of Computational Chemistry*, 32(1):170–173, jan 2011.

[9] H. Zhang, I. Goodfellow, D. Metaxas, and A. Odena. Self-attention generative adversarial networks. *36th International Conference on Machine Learning, ICML 2019*, 2019-June:12744–12753, 2019.

[10] Y. Zhou, X. Qi, Y. Liu, F. Zhang, and H. Yan. Dna-nanoscaffold-assisted selection of femtomolar bivalent human alpha-thrombin aptamers with potent anticoagulant activity. *ChemBioChem*, 20(19):2494–2503, 2019.
